# Group II Intron-Like Reverse Transcriptases Function in Double-Strand Break Repair by Microhomology-Mediated End Joining

**DOI:** 10.1101/2022.03.14.484287

**Authors:** Seung Kuk Park, Georg Mohr, Jun Yao, Rick Russell, Alan M. Lambowitz

## Abstract

Bacteria encode free-standing reverse transcriptases (RTs) of unknown function that are closely related to group II intron-encoded RTs. Here, we found that a *Pseudomonas aeruginosa* group II intron-like RT (G2L4 RT) with YIDD instead of YADD at its active site functions in DNA repair in its native host and when transferred into *Escherichia coli*. G2L4 RT has biochemical activities strikingly similar to those of human DNA repair polymerase *θ* and uses them for translesion DNA synthesis and double-strand break repair (DSBR) via microhomology-mediated end-joining (MMEJ) *in vitro* and *in vivo*. We also found that a group II intron RT can function similarly to G2L4 RT in DNA repair, with reciprocal substitutions at the active site showing an I residue favors MMEJ and an A residue favors primer extension in both enzymes. The DNA repair functions of these enzymes utilize conserved structural features of non-LTR-retroelement RTs, including human LINE-1 and other eukaryotic non-LTR-retrotransposon RTs, suggesting such enzymes may have an inherent ability to function in DSBR in a wide range of organisms.

## Introduction

Reverse transcriptases (RTs) are best known for their crucial roles in the replication of human pathogens, such as retroviruses and hepatitis B virus, and as tools for biotechnological applications, such as high-throughput RNA sequencing (RNA-seq) and RT-qPCR (Martín-Alonso et al., 2021). However, RTs are found in all domains of life and are common in bacteria, where they are thought to have evolved from an RNA-dependent RNA polymerase (Lambowitz and Belfort, 2015). The most prevalent bacterial RTs are those encoded by mobile group II introns, retrotransposons that are evolutionary ancestors of spliceosomal introns and the spliceosome, as well as retrovirus and other retroelements in eukaryotes (Lambowitz and Zimmerly, 2011; Lambowitz and Belfort, 2015). Extant bacteria also harbor a variety of other RTs, all of which are closely related to group II intron RTs and some of which have been found to perform cellular functions. The latter include diversity-generating retroelement RTs, CRISPR-associated RTs, abortive phage infection RTs, and retron RTs, which were shown recently to function in phage defense systems (Liu et al., 2002; Wang et al., 2011; Silas et al., 2016; Gao et al., 2020; Millman et al., 2020; González-Delgado et al., 2021). In addition to these characterized enzymes, bacteria contain a wide variety of other unexplored group II intron-like RTs that are encoded by free-standing conserved ORFs in bacterial genomes and whose biochemical activities and biological functions remain unknown (Kojima and Kanehisa, 2008; Simon and Zimmerly, 2008; Zimmerly and Wu, 2015).

Group II intron and other bacterial RTs belong to a larger family of non-LTR-retroelement RTs, which includes human LINE-1 and other eukaryotic non-LTR-retrotransposons RTs (Xiong and Eickbush, 1990). These non-LTR-retroelement RTs are homologous to retroviral RTs but have distinctive conserved structural features that impact RT activity, including an N-terminal extension (NTE) with an RT0 loop region, two insertions, RT2a and RT3a, between universally conserved RT sequence blocks (RT1-7), and a larger thumb domain with three instead of two *α*-helices (Xiong and Eickbush, 1990; Blocker et al., 2005; Qu et al., 2016; Stamos et al., 2017; Haack et al., 2019) (Figure 1A). A crystal structure of a full-length group II intron RT (*Geobacillus stearothermophilus* GsI-IIC RT) in complex with template-primer and incoming dNTP showed that group II intron RTs are similar to retroviral RTs in folding into a hand-like structure with fingers, palm, and thumb forming a cleft that binds the template-primer at the RT active site, but with the NTE/RT0 loop and RT2a insertions contributing to tighter binding pockets for template/primer and incoming dNTP that could contribute to the relatively high fidelity and processivity of these enzymes (Mohr et al., 2013; Stamos et al., 2017) (Figure 1B). The NTE/RT0 loop also plays a key role in a proficient group II intron RT template-switching activity that is dependent upon a short base-pairing interaction between the 3’ ends of the donor and acceptor nucleic acids (Mohr et al., 2013; Stamos et al., 2017; Lentzsch et al., 2019, 2021).

**Figure 1.**
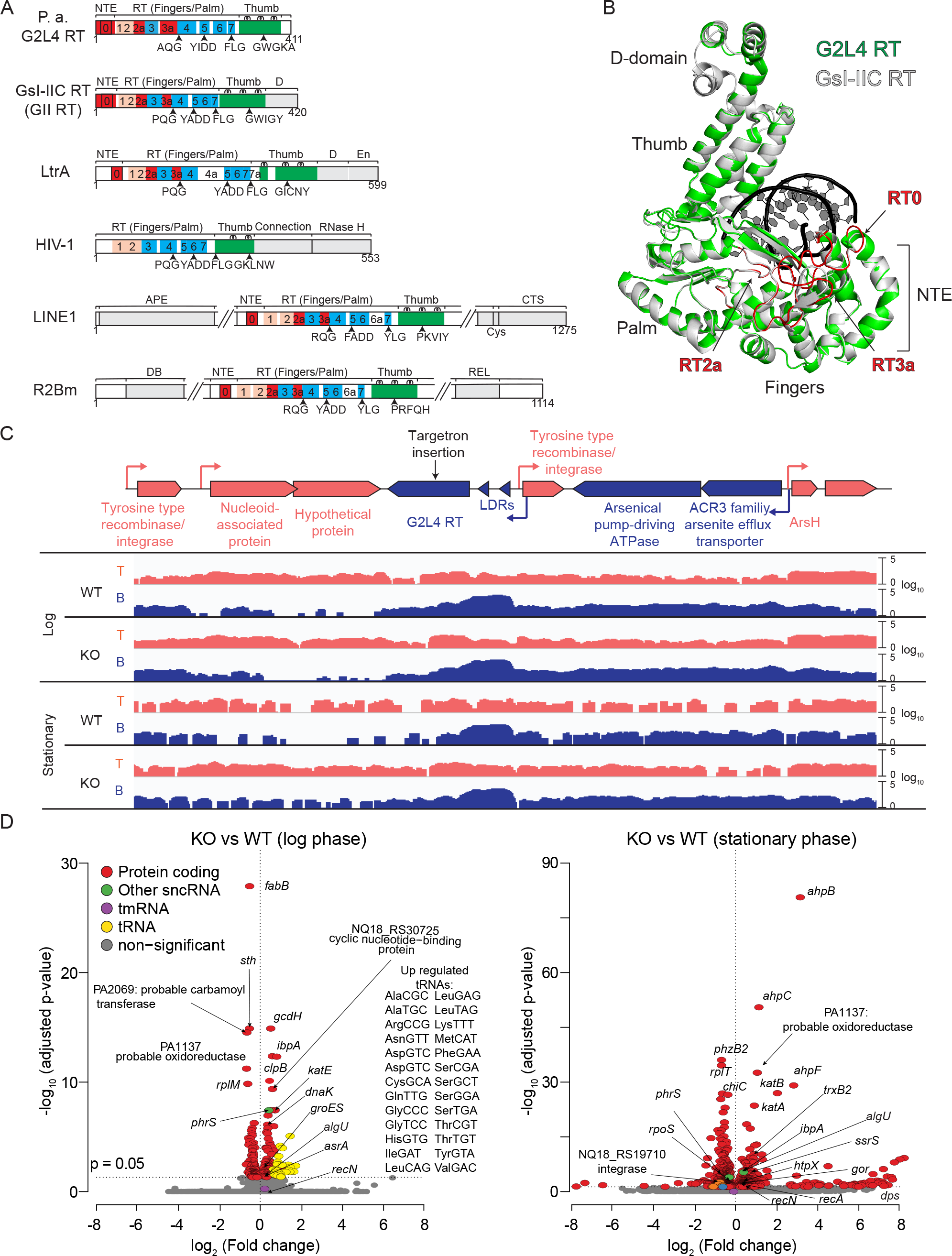
Characteristics of *P. aeruginosa* G2L4 RT and TGIRT-seq of cellular RNAs of G2L4 RT WT and knock out strains in log and stationary phases (A) Schematics comparing *P. aeruginosa* G2L4 RT, *G. stearothermophilus* GsI-IIC (GII) RT, and *Lactococcus lactis* Ll.LtrB group II intron RT (LtrA protein) with retrovirus HIV-1 RT and non-LTR retrotransposon human LINE-1 and *Bombyx mori* R2Bm RTs. Protein regions: NTE, N-terminal extension; RT1-7, conserved sequences blocks found in all RTs; RT0, RT2a and RT3a, insertions relative to retroviral RTs (red); fingers (pink); palm (blue); thumb (green); Other domains (gray): group II intron RT DNA-binding domain (D) and DNA endonuclease (En) domains; APE, apurinic endonuclease domain; CTS, conserved carboxy-terminal segment; Cys, cysteine-rich conserved sequence. (B) Three-dimensional model of G2L4 RT (green) constructed by I-TASSER (Yang and Zhang, 2015) superimposed on the crystal structure of GII RT (gray; PDB: 6AR1). Primer-template (black). (C) Map of *P. aeruginosa* AZPAE12409 genomic region encoding G2L4 RT and coverage plots based on TGIRT-seq of cellular RNAs in the WT and G2L4 RT KO strains in log and stationary phases. Genes with protein-coding sequences on the top and bottom strand are shown in red and blue, respectively, with predicted promoters identified by BacPP (de Avila e Silva et al., 2011) indicated by red and blue arrows, respectively. The G2L4 RT targetron insertion site into the top (antisense) strand of the G2L4 RT gene is indicated by a black arrow. LDRs, long direct repeats. Top and bottom strand read coverage on a log10 scale is shown below the map in the same colors. (D) Volcano plots showing differences in gene expression in G2L4 WT and KO strains in log and stationary phases. Read counts were DEseq2 normalized (Love et al., 2014). Significantly enriched genes in G2L4 RT KO versus the WT strain in log and stationary phases (adjusted p value *≤*0.05) are denoted in different colors: protein coding, red; tmRNA, purple, tRNA, yellow; other sncRNA, green; non-significant, gray.

All RTs have an active site containing a conserved F/YxDD motif, whose aspartate residues are critical for binding of catalytic Mg^2+^ ions (Argos, 1988; Xiong and Eickbush, 1990). In group II intron RTs, this motif is typically F/YADD, with the conserved alanine being part of the network of structural features that could impact fidelity and processivity (Stamos et al., 2017). However, in some families of bacterial RTs, this conserved alanine is replaced by a different conserved amino acid (I, V, M, H, S, or R) (Kojima and Kanehisa, 2008; Simon and Zimmerly, 2008; Zimmerly and Wu, 2015).

Here, we found via gene disruption and complementation that a *Pseudomonas aeruginosa* RT belonging to a family denoted Group II-Like 4 (G2L4 RT; Zimmerly and Wu, 2015) with a conserved YIDD at its active site functions in DNA repair in its native host. Further analyses showed that: (i) G2L4 RT functions in translesion DNA synthesis and DSBR via MMEJ *in vitro* and *in vivo*; (ii) a group II intron RT (the *G. stearotheromophilus* GsI-IIC RT, denoted GII RT) with YADD at its active site can function similarly in DNA repair; (iii) an I residue at the active site adapts these enzymes for MMEJ at the expense of primer extension; and (iv) the MMEJ activity of both enzymes is dependent upon the RT0 loop, a conserved structural feature of non-LTR-retroelement RTs. Our results demonstrate that RTs have the previously unsuspected ability to function in DSBR and suggest that non-LTR-retroelement RTs may have an inherent ability do so in a wide range of organisms.

## Results

### Identification and characteristics of G2L4 RTs

To identify candidate G2L4 RTs for detailed analysis, we performed a BLASTP search of GenBank using the sequence of a G2L4 RT (ABB74237) from *Nitrospira multiformis* ATCC 25196 as the search sequence. We identified ∼250 unique G2L4 RTs in gram negative *α*, *β*, *γ*, and a few *δ* proteobacteria. Among the *γ* proteobacteria, many G2L4 RTs were found in *Pseudomonas* spp., and we decided to focus on a member of this group (WP_034031052) found in *P. aeruginosa* strain AZPAE12409 because this strain was readily available and genetic and molecular biological methods are well-established for *P. aeruginosa*.

The genomic region encoding the G2L4 RT in *P. aeruginosa* AZPAE12409 has the characteristics of a horizontally transferred genetic element, including lower GC content and genes with codon usage that differs from that of neighboring host genes (Figure S1). In ∼70% of the cases, the G2L4 RT ORF was preceded by two palindromic ∼140-bp long direct-repeats (LDRs) separated by an ∼240-bp spacer, whose sequences were conserved in different *Pseudomonas* strains (Figure 1C and Figure S2). Further, we found that the G2L4 ORF and its upstream LDRs were inserted in different genomic regions in different *Pseudomonas* spp. strains, frequently in proximity to ORFs encoding putative tyrosine recombinases or other DNA integrases (Figure 1C), raising the possibility that the G2L4 RT might be associated with an independently mobile genetic element. Like group II intron and other non-LTR-retroelement RTs, G2L4 RT contains an N-terminal extension (NTE) with an RT0 loop and RT2a and RT3a insertions between conserved RT sequence blocks, which are absent in retroviral and other LTR-retroelement RTs (Figure 1A and Figure S3). The predicted secondary and tertiary structures of the G2L4 RT closely matched the known structure of GII RT, with the major differences being a longer RT3a insertion and small insertions downstream of RT6 and in the thumb domain (Figure 1B and Figure S3A). The RT0 loop, which plays a critical role in the template-switching activity of group II intron and other non-LTR-retro- element RTs (Jamburuthugoda and Eickbush, 2014; Lentzsch et al., 2019, 2021), is structurally similar between G2L4 and group II intron RTs but differs in having conserved serine residues (Figure S3B). G2L4 RTs also differs from group II intron RTs in lacking C-terminal DNA-binding (D) and DNA endonuclease (En) domains, which function in binding and cleaving DNA target sites during group II intron retrohoming (Figure 1A; San Filippo and Lambowitz, 2002).

### Analysis of G2L4 RT knock out strains

To investigate the function of the G2L4 RT in its native host, we used targetron mutagenesis (Karberg et al., 2001; Yao and Lambowitz, 2007) to disrupt the G2L4 RT gene in *P. aeruginosa* AZPAE12409 (Figure S4A). We obtained two G2L4 RT disruptants in which the targetron had inserted at the same site in the antisense orientation relative to the G2L4 ORF (Figure 1C and Figure S4B). After curing the targetron-expression plasmid, Southern hybridization and genomic DNA sequencing showed a single targetron insertion at the desired site in both knock out (KO) strains, with one disruptant (KO2) having no other changes and the other (KO1) having a single missense mutation in an ORF encoding cell division protein ZapA (Figure S4C and D and Supplementary File). The wild-type (WT) and both KO strains had similar growth rates through log and stationary phases in complete medium, indicating that G2L4 RT is not an essential protein in its native host (Figure S4E). We selected the KO2 strain lacking the secondary mutation for further analysis.

To assess the effect of the G2L4 RT disruption on gene expression, we analyzed the transcriptomes of the G2L4 WT and KO strains in log (15 h) and stationary (30 h) phases by using TGIRT-seq of rRNA-depleted, chemically fragmented whole-cell RNAs, an RNA-seq method that enables simultaneous profiling of all RNA biotypes without size selection (Figure S5A; Tables S1) (Nottingham et al., 2016; Boivin et al., 2018). In the resulting TGIRT-seq datasets, 70-80% of the reads mapped to protein-coding genes, with the remainder mapping to small non-coding RNAs (sncRNAs; Figure S5B). The most abundant sncRNAs were tRNAs, RNase P RNA, and tmRNA, a sncRNA that functions in releasing mRNAs from stalled ribosomes (Müller et al., 2021). In both the WT and KO strains, the proportion of tRNA reads decreased in stationary phase, while the proportion of tmRNA reads increased, consistent with its regulation by RpoS, a bacterial stress response sigma factor that is up-regulated in stationary phase (Figure S5B; Himeno et al., 2014).

Coverage plots showed relatively uniform shallow read depth on both strands over the G2L4 RT coding region, but with the bottom strand showing 20-fold higher read depth over the 140-bp LDRs preceding the RT (Figure 1C).

Volcano plots comparing the relative abundance of different RNAs in the WT and KO strains showed differences in both log and stationary phases, but with more differentially expressed genes and larger fold changes in stationary phase (Figure 1D). A notable difference in the KO strain in log phase was the higher expression level of tRNAs recognizing rare codons, including those used in the G2L4 RT ORF (Figure 1D and Figure S5C), raising the possibility that G2L4 RT synthesis might be subject to post-transcriptional regulation and implying that G2L4 RT disruption may directly or indirectly activate a pathway that leads to upregulation of tRNAs that recognize rare codons, possibly part of a global stress response. In stationary phase, the KO strain showed a significantly higher expression level of 6S RNA, a sncRNA that helps regulate σ^70^-dependent transcriptional responses (Figure S5D; Cavanagh and Wassarman, 2014).

Among the numerous differentially expressed protein-coding genes in stationary phase were three encoding transcriptional regulatory proteins. The gene encoding sigma factor AlgU, which induces osmotic, oxidative, and temperature stress responses, was expressed at higher levels in the KO than the WT strain in both log and stationary phases (Schurr and Deretic, 1997; Schnider-Keel et al., 2001); the gene encoding sigma factor RpoS, which is induced in stationary phase for tolerance of high osmolarity, DNA damage, and oxidative stress, was up regulated in stationary phase in both the WT and KO strains (Jaishankar and Srivastava, 2017); and the gene encoding the LexA repressor, a DNA damage sensor whose cleavage after interaction with RecA at double-strand breaks (DSBs) triggers an SOS response, was down regulated in stationary phase in both the WT and the KO strain (Figure 2A and Figures S6 and S7; Cirz et al., 2006; Jin et al., 2007; Simmons et al., 2008; Kreuzer, 2013).

**Figure 2.**
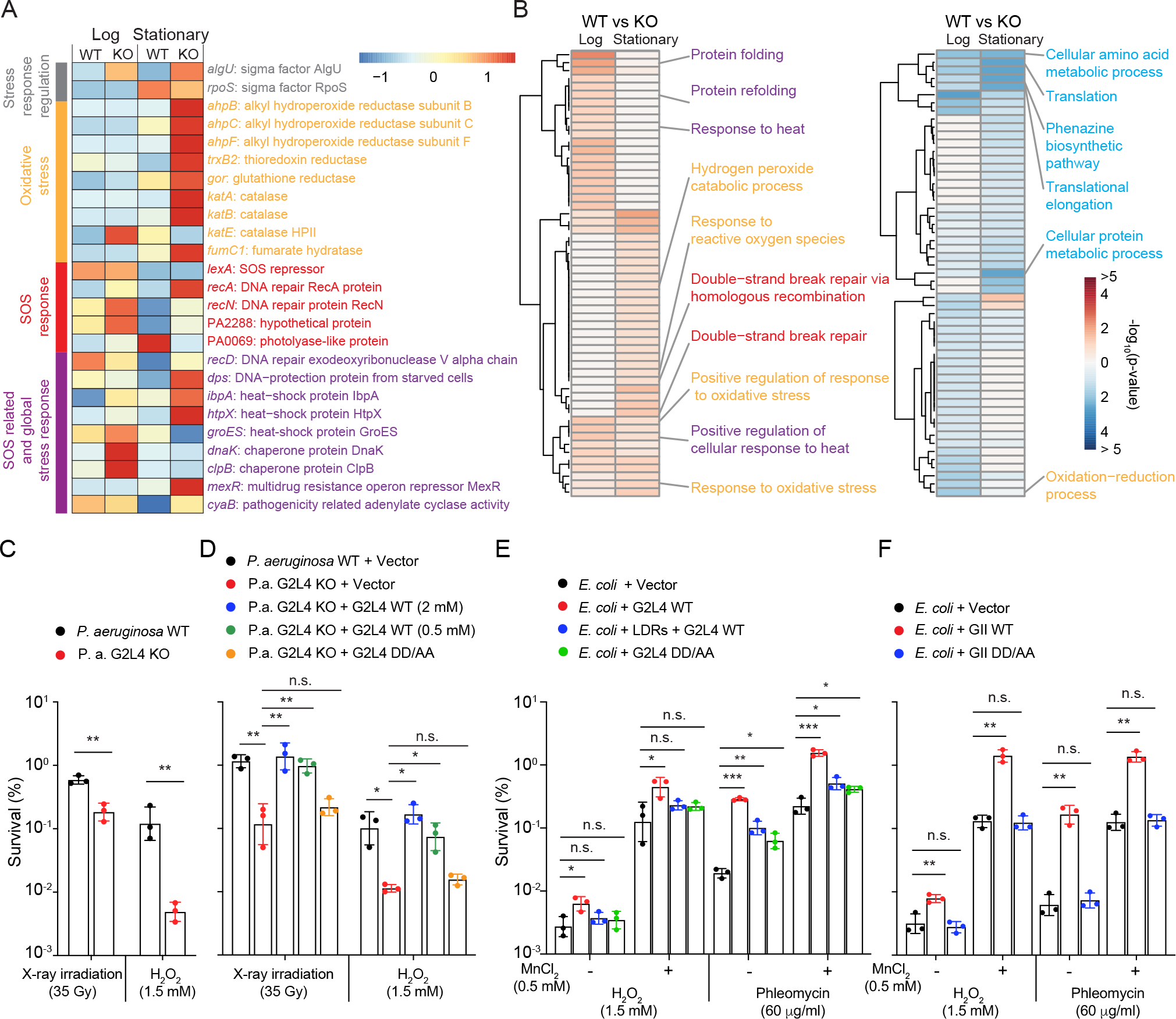
G2L4 RT knock out results in up regulation of oxidative stress and DNA damage response pathways and effect of G2L4 RT expression on DNA damage survival (A) Heatmaps comparing significantly regulated genes in cellular stress and DNA damage response pathways in WT versus G2L4 RT KO strains of *P. aeruginosa* AZPAE12409 in log and stationary phase. The heat maps are based on four replicates of TGIRT-seq of whole cell RNAs in log and stationary phase. The color scale for the heatmaps (top right) is based on the Z score for the mean of the sample-medians of the 4 replicates divided by the standard deviation of the samplemedians. Gene names are color coded by function: stress response regulation, gray; oxidative stress, orange; SOS response, red; SOS related global stress response, purple. Box plots showing differential expressions of genes in the heat map in WT and G2L4 RT KO strains are shown in Figure S6. (B) Heatmaps of up-regulated (left, red) and down-regulated (right, blue) GO terms in G2L4 RT KO versus WT strains in log and stationary phase. The color scale (bottom right) shows the - log10(p-value) of GO term enrichment for genes that were significantly up (log2FC>0, adjusted p-value *≤*0.05) or down (log2FC<0, adjusted p-value *≤*0.05) regulated in the G2L4 RT KO versus WT strains in log or stationary phases. GO term names are color coded based on the following groupings: oxidative stress, orange; SOS response, red; SOS related global stress response, purple; Translation and protein metabolic processes, blue. A complete GO terms heat map is shown in Figure S7. (C) and (D) Survival of *P. aeruginosa* AZPAE12409 WT and KO strains after X-ray irradiation (35 Gy) or H2O2 (1.5 mM) treatment without (C) or with (D) expression of WT or RT- deficient DD/AA mutant G2L4 RTs. RTs were expressed from plasmid pBL1 using an m-toluic acid-inducible promoter with the WT protein expressed using two different concentrations of m- toluic acid (0.5 and 2 mM) to vary the protein expression level. The expressed RT proteins have an N-terminal maltose-binding protein (MBP) tag, which improves the solubility and stability of group II intron RTs (Mohr et al., 2013), and a C-terminal 8X His tag used for detection in immunoblots (Figure S8), while the vector control expresses MBP with an 8X His tag. (E) Survival of *E. coli* HMS174 (DE3) expressing WT G2L4 and DD/AA mutant RTs after treatment with H2O2 or phleomycin with or without MnCl2 added to the growth medium compared to a vector control. The WT G2L4 RT gene was expressed with or without the upstream long direct repeats (LDRs). (F) Survival of *E. coli* HMS174 (DE3) expressing WT or DD/AA mutant GII RTs after treatment with H2O2 or phleomycin with or without MnCl2 added to the growth medium. GII RTs were expressed from pBL1 similarly to G2L4 RTs with an N-terminal MBP tag and a C-terminal 8X His tag (Figure S8). All experiments were repeated three times, with errors bar indicating the standard deviation. n.s., non-significant; p-values < 0.05, *; < 0.01, **; < 0.001, ***.

The altered expression levels of these transcriptional regulatory proteins were correlated with the up-regulation of pathways and genes related to DNA repair and oxidative stress in the KO compared to the WT strain in log and/or stationary phases, but more pronounced in the KO strain in stationary phase (Figure 2A and B). These changes included up-regulation of genes involved in both oxidative stress response *ahpB*, *aphC*, and *aphF* (alkyl hydroperoxide reductase), *trxB2* (thioredoxin reductase), *gor* (glutathione reductase), *katA*, *katB* and *katE* (catalase), *fumC1* (fumarate hydratase class II) and SOS-induced DNA damage response, including *recA* and *recN*, which encode proteins that function in double-strand break repair (DSBR), and PA2288 and PA0069, which encode a hypothetical protein and a photolyase-like protein, respectively, and are among 15 identified SOS response genes in *P. aeruginosa* (Figure 2A and Figure S6B; Cirz et al., 2006). The KO strain in stationary phase also showed up-regulation of the genes encoding the RecD helicase subunit of RecBCD, which functions in DNA recombination and repair, and DPS, a DNA-binding protein that protects DNA in starved cells and is part of the RpoS regulon (Loewen et al., 1998; Dong et al., 2008). Heat shock proteins genes (*ibpA*, *htpX* and *groES*), protein chaperone genes (*dnaK* and *clpB*), multidrug resistance repressor gene (*mexR*), and the pathogenicity related adenylate cyclase gene (*cyaB*) were also elevated in log and/or stationary phases in the KO compared to the WT strain (Figure 2B and Figure S6A). The exacerbated DNA damage responses overseen by the three transcriptional regulatory proteins (AlgU, RpoS, and LexA) in log and/or stationary phase in the KO compared to the WT strain suggested that G2L4 might function in DNA repair.

### G2L4 RT functions in DNA repair *in vivo*

To investigate if G2L4 RT functions in DNA repair in its native host, we tested survival of the *P. aeruginosa* WT and G2L4 RT KO strains after inducing DNA damage. We found that the KO strain was more sensitive than WT to both X-ray irradiation, which causes double-strand breaks (DSBs), and H2O2 which causes oxidative DNA damage and can also induce DNA breaks (Figure 2C; Driessens et al., 2009). For both treatments, survival was restored to at or near WT levels by expressing WT G2L4 RT from a plasmid using an m-toluic acid inducible promoter, but not in a vector control nor in cells expressing an RT-deficient mutant G2L4 RT in which the conserved aspartates at the RT active site were replaced with alanines (denoted DD/AA; Figure. 2D and Figure S8A). By varying the concentration of the m-toluic acid inducer, we confirmed rescue after X-ray irradiation by WT G2L4 RT over a range of protein expression levels, including the lower expression level seen for the G2L4 RT DD/AA mutant protein (Figure 2D and Figure S8B).

Similarly in *E. coli* HMS174 (DE3) *recA^-^*, WT G2L4 RT expressed from a plasmid increased resistance to H2O2 as well as to phleomycin, which also induces DSBs (Figure 2E; Merrikh et al., 2009). As Mn^2+^ was known to enhance the activities of DNA repair polymerases (Hutfilz et al., 2019) and Mn^2+^ levels were known to be lower in *E. coli* than in *Pseudomonas* spp. (Anjem et al., 2009; Cunrath et al., 2016), we tested whether adding MnCl2 to the growth medium increases survival from DNA damage in *E. coli* expressing G2L4 RT and found this was the case (Figure 2E). In the *E. coli* assays, the ability of G2L4 RT to significantly increase cell survival from H2O2 or phleomycin was decreased but not completely abolished for the RT-deficient DD/AA mutant (Figure 2E). The latter finding indicates that the RT activity of G2L4 RT plays a major role in DNA damage survival, but leaves open the possibility that other activities of G2L4 RT may also contribute (Figure 2D and E). Co-expression of G2L4 RT with the upstream LDRs modulated its ability to increase cell survival from H2O2- and phleomycin-treatments in *E. coli* without affecting the protein expression level (Figure 2E and Figure S8C). Collectively, these results indicate that G2L4 RT functions as a DNA repair polymerase in both its native host and *E. coli*.

### G2L4 and GII RT have robust DNA polymerase activity

To investigate if G2L4 RT has the enzymatic activities required to function as a DNA repair polymerase and how these activities might differ from those of a group II intron RT, we carried out parallel biochemical assays with G2L4 RT and GII RT, a thermostable group II intron RT that retains high activity at lower temperatures (Mohr et al., 2013). In addition to the WT enzymes, we tested both proteins with reciprocal I/A substitutions and DD/AA mutations at the RT active site. Based on pilot experiments showing that G2L4 RT prefers low salt concentrations (Figure S9) and the finding that supplemental Mn^2+^ increased DNA damage survival in *E. coli* expressing G2L4 RT (see above), the biochemical assays were done in reaction medium containing 20 mM NaCl and 10 mM MgCl2 at 37°C in the absence or presence of 1 mM MnCl2. These assay conditions are similar to those used for human DNA polymerase *θ*, a DNA repair polymerase that repairs double-strand breaks via MMEJ (Kent et al., 2015; Chandramouly et al., 2021).

First, we assayed the DNA polymerase and RT activities of the G2L4 and GII RTs by primer extension using 3’-blocked 50-nt DNA or RNA oligonucleotide templates of identical sequence with different length DNA primers annealed at their 3’ ends. These experiments showed that WT G2L4 had high primer extension activity on both the DNA and RNA templates with primers up to 5-nt long, but differed markedly from GII RT in being unable to efficiently use primers *≥*10 nt (Figure 3A and B). Time courses with 5-nt-DNA primers showed that WT G2L4 RT prefers DNA over RNA templates, with the rate of primer extension on both templates increased 4- to 6-fold in the presence of Mn^2+^ (Figure S10A and B). Parallel assays showed that WT GII RT could use both short and long primers (Figure 3C and D), but with time courses revealing some preference for shorter primers, particularly on the DNA template, and little effect of added Mn^2+^ on both templates (Figure S10C and D). As expected, the primer extension activity of both enzymes was abolished by DD/AA mutations at the RT active site (Figure 3A-D).

**Figure 3.**
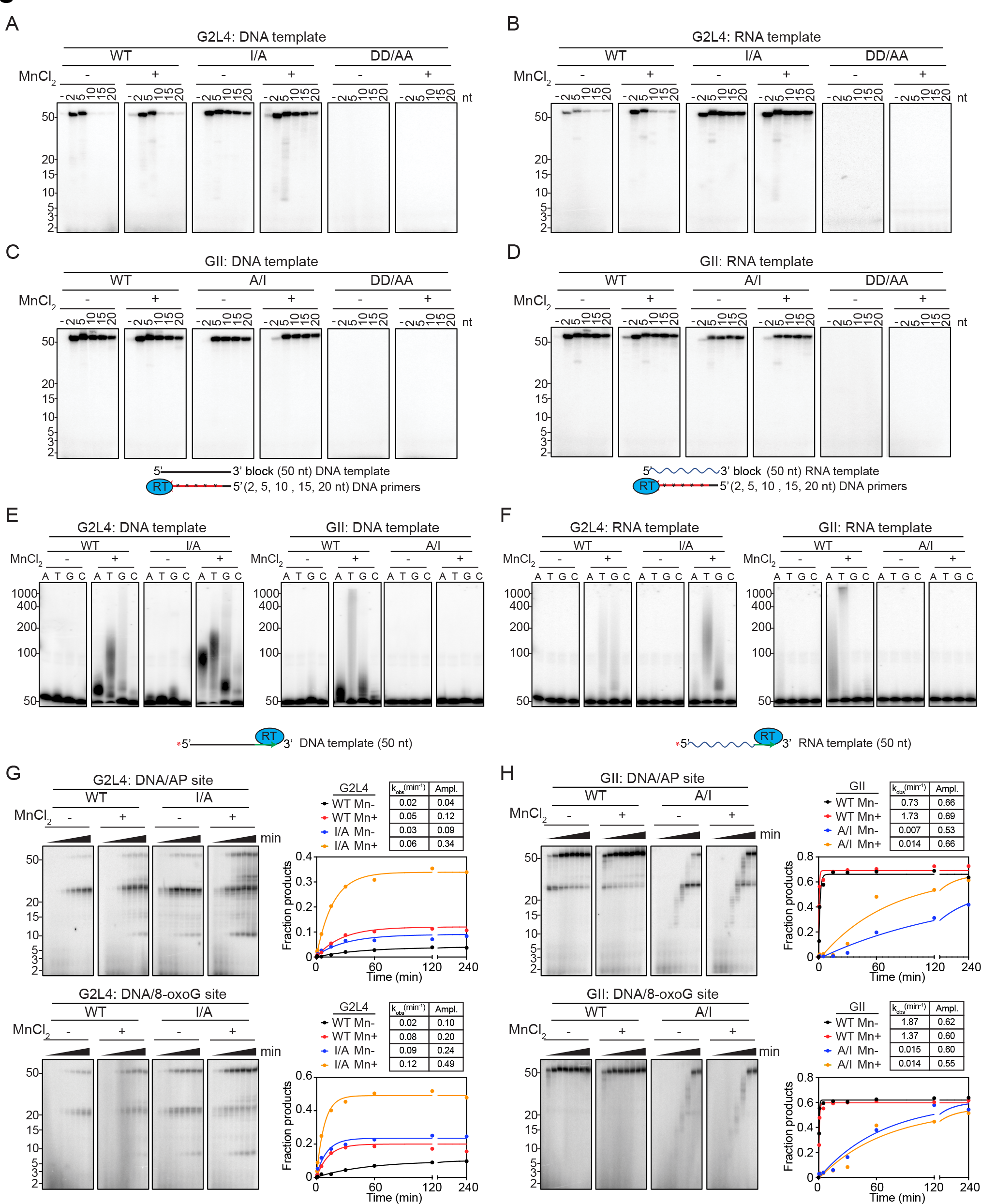
Biochemical activities of WT and mutant G2L4 and GII RTs (A-D) Primer extension. Reactions were done as described in Methods with 500 nM WT or mutant G2L4 and GII RTs and 250 nM 50-nt DNA or RNA templates with a 3’-blocking group (an inverted dT residue) annealed to primers of various lengths (100 µM 2 nt; 50 µM 5 nt; or 500 nM 10, 15, or 20 nt primers, respectively) in the presence or absence of 1 mM MnCl2. Reactions were initiated by adding 1 mM labeled dNTPs (an equimolar mix of 1 mM each dATP, dCTP, dGTP, and dTTP plus [*α*-^32^P]-dTTP) and incubated for 240 min at 37°C. Time courses are shown in Figure S10. (E and F) Terminal transferase. Reactions were done with 500 nM WT or mutant G2L4 or GII RTs, 10 nM 5’-labeled 50-nt DNA or RNA substrate without a 3’ blocking group and 1 mM of the indicated dNTP in the presence or absence of 1 mM MnCl2 for 20 min at 37°C. Time courses are shown in Figure S11. (G and H) Translesion DNA synthesis. Reaction time courses were done under the same condition as primer extension reactions with 500 nM WT or mutant G2L4 and GII RTs and 250 nM 50-nt 3’ blocked DNA or RNA templates containing an AP site or 8-oxoguanine positioned 23 nt from the 3’ end. The numbers to the left of the gels indicate the positions of size markers in a parallel lane. The tables above the plots show rate constants (*k*obs) and amplitudes (Ampl.) for production of the labeled 50-nt DNA product obtained by fitting the data to a first- order rate equation.

Notably, I/A substitution at the active site of the G2L4 RT increased the rate of primer extension and alleviated the strict requirement of G2L4 RT for short primers, enabling the use of primers up to 20 nt (Figure 3A and B; time courses Figure S10E and F). The mutant enzyme still showed a preference for DNA over RNA templates, but added Mn^2+^ had less of an effect. By contrast, the major effect of the reciprocal A/I substitution in the GII RT was to decrease the rate of primer extension (Figure 3C and D; time courses in Figure S10G and H). These findings indicate that the larger I residue at the active site in G2L4 RT dictates its strong preference for short DNA primers, while substitution of the smaller A residue enables G2L4 RT to use longer primers and increases the rate of primer extension on both DNA and RNA templates. The finding that both the G2L4 and GII RTs have robust DNA polymerase activity was expected for G2L4 RT functioning as a DNA repair polymerase and raised the possibility that the GII RT might also be capable of functioning as a DNA repair polymerase. Prompted by the biochemical assays, we confirmed that expression of GII RT in *E. coli* increased cell survival from treatment with H2O2 or phleomycin as well or better than G2L4 RT with the ability to do so inhibited by the DD/AA mutations at the RT active site (Figure 2F and Figure S8D).

### G2L4 and GII RT have Mn^2+^-stimulated terminal transferase activity

Human DNA repair polymerase *θ*, which functions in DSBR via MMEJ, has a Mn^2+^-dependent terminal transferase activity that extends single-stranded 3’-DNA overhangs at 5’-resected DSB sites until they can base pair with short complementary regions (microhomologies) in the 3’ overhang on the opposite side of the break (Kent et al., 2016). We found that WT G2L4 and GII RTs also have a Mn^2+^-stimulated terminal transferase activity with nucleotide preferences T=A>G>>C and with both enzymes showing a preference for single-stranded DNA over RNA substrates (Figure 3E and F; time courses in Figure S11). The G2L4 I/A mutation had little effect on terminal transferase activity, while the reciprocal A/I substitution in GII RT strongly inhibited this activity (Figure 3E and F).

### Both G2L4 and GII RT read through DNA lesions

Human DNA polymerase *θ* has a translesion DNA synthesis activity that enables it to bypass DNA lesions in damaged DNA (Seki et al., 2004). To investigate if G2L4 and GII RTs could read through DNA lesions, we performed primer extension assays using the 50-nt DNA template containing an AP (apurinic/apyrimidinic) site or 8-oxoguanine, which is produced by oxidative DNA damage (Sidorenko et al., 2007), positioned 23 nt from the 3’ end of the DNA. The primer extension reactions were done as above with a short 5-nt DNA primer, which can be used efficiently by both enzymes (see above).

We found that G2L4 RT could read through both lesions, with this ability being strongest for the I/A mutant in the presence Mn^2+^, likely reflecting the higher primer extension activity of the mutant protein (Figure 3G). WT GII RT, which has high primer extension activity, was even more efficient than G2L4 RT in reading though the lesion site in the presence or absence of Mn^2+^, and this ability was strongly inhibited by the A/I substitution, which inhibits primer extension activity (Figure 3H). These findings indicate that both the G2L4 and GII RTs can read through damaged DNA templates with the ability to do so favored by the smaller A residue at the RT active site, which enables higher primer extension activity.

### G2L4 and GII RT promote snap-back DNA synthesis by annealing short microhomologies between the 3’ end and upstream regions of DNA templates

Human DNA polymerase *θ* functions in DSBR by an error prone process (Alternative End Joining; Alt-EJ), which involves annealing short stretches of complementary nucleotides (microhomologies) between single-stranded 3’ overhangs resulting from 5’-DNA strand resection on both sides of a double-strand break and then using the 3’ ends of the annealed microhomology as primers to fill in the single-stranded gaps (Black et al., 2016; Ramsden et al., 2022). The ability of DNA polymerase *θ* to anneal short microhomologies enables a distinctive biochemical activity termed snap-back DNA replication in which the enzyme uses the unblocked 3’ end of a DNA template as a primer to initiate DNA synthesis at short stretches of complementary nucleotides located farther upstream (Kent et al., 2016; Black et al., 2019).

To assay the snap-back replication activity of G2L4 and GII RTs, we used 5’-labeled 50-nt DNA and RNA oligonucleotides identical to those used above for primer extension assays but with unblocked 3’ ends and no added primer (Figure 4A and B). The products of the reaction were analyzed in a non-denaturing 12% polyacrylamide gel, which makes it possible to distinguish snap back replication from MMEJ products in MMEJ assays below (Kent et al., 2015, 2016).

**Figure 4.**
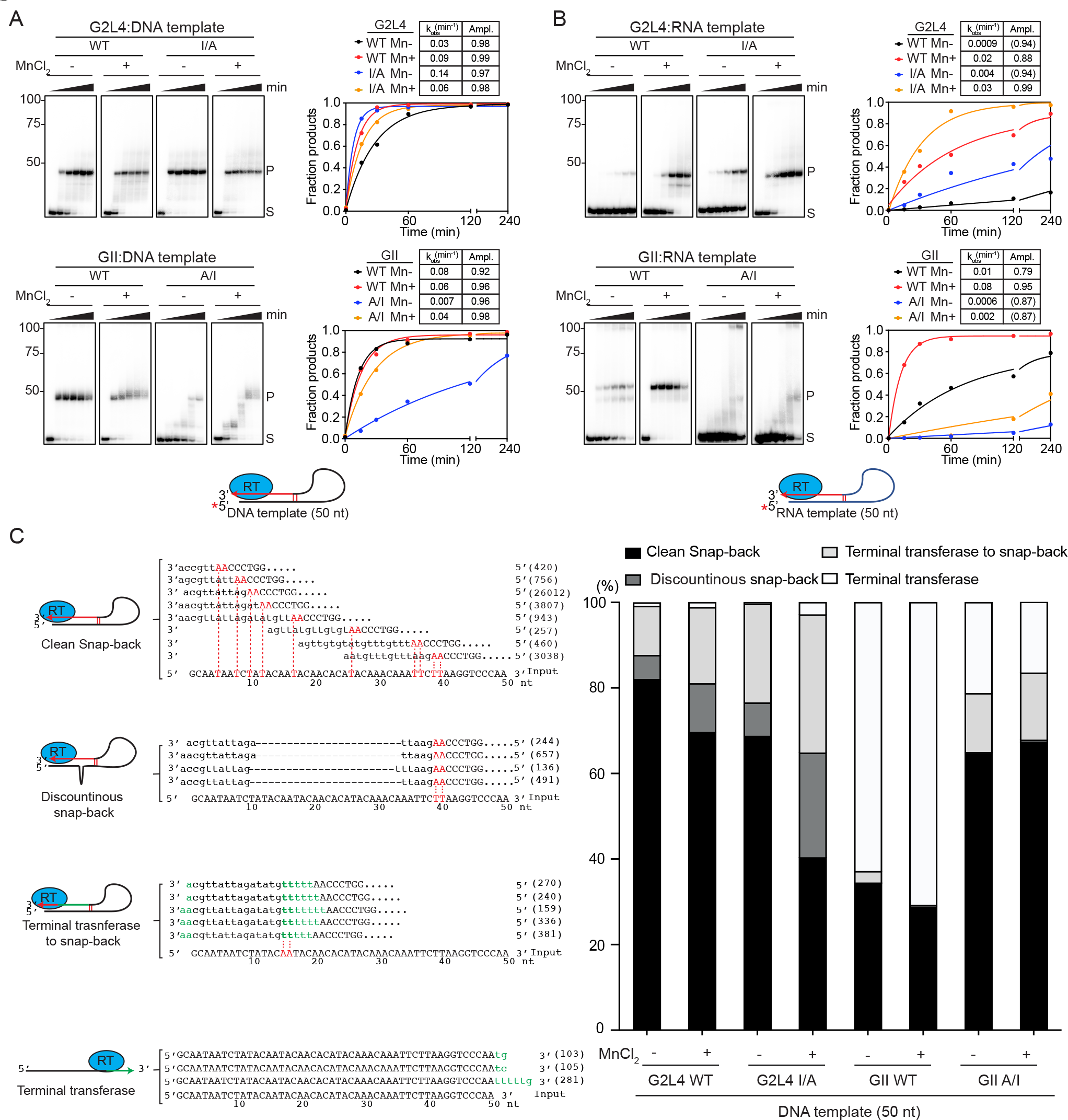
Snap-back DNA synthesis by WT and mutant G2L4 and GII RTs (A and B) Snap-back DNA synthesis assays. Reactions were done as described in Methods with 500 nM WT or mutant G2L4 and GII RTs with or without 1 mM MnCl2. The numbers to the left of the gels indicate the positions of size markers in a parallel lane. The plots show the fraction of substrate (S) that is converted to products extending up to the major product band (P), and the tables above the plots show rate constants (*k*obs) and amplitudes (Ampl.) obtained by fitting the data to a first-order rate equation. (C) High-throughput sequencing of snap-back DNA synthesis products. Reactions were done as above with 1 µM unlabeled DNA template for 3 h at 37°C. Products were sequenced with an Illumina MiSeq to obtain up to 1 million 2 x 75 nt paired end reads, and the sequences were aligned by clustalW (see Methods). Schematics and representative sequences of different categories of products obtained with WT G2L4 RT without MnCl2 are shown to the left. Template nucleic acid sequences are in upper case letters, and product sequences are in lower case letters. Nucleotides involved in short base-pairing interactions between the 3’ AA and internal regions of the DNA template are indicated in red, and nucleotides added to the 3’ end of the template by terminal transferase activity are indicated in green. The number of reads corresponding to the same sequence is indicated in parenthesis. The stacked bar graphs to the right show the proportions of different products obtained with WT and mutant G2L4 and GII RTs in the absence or presence of 1 mM MnCl2.

The WT and mutant G2L4 and GII RTs with reciprocal I/A substitution all gave products with electrophoretic mobility expected for snap-back DNA products, with all showing a preference for the DNA over the RNA template (Figure 4A and B). WT G2L4 and GII RTs and G2L4 I/A mutant RT had robust snap-back DNA synthesis activities on the DNA template, while the GII A/I mutant RT had lower activity and gave intermediate size products at short time points (Figure 4A, bottom) likely reflecting its decreased rate of primer extension. Addition of Mn^2+^ stimulated snap-back DNA synthesis on both DNA and RNA templates in most cases, the exceptions being WT GII and G2L4 I/A RTs, the two proteins with A at the active site, whose rates of product formation on DNA substrates were close to maximal with little if any effect by Mn^2+^ (Figure 4A and B).

We sequenced the products of snap-back replication on DNA substrates by using a thermostable group II intron reverse transcriptase (TGIRT)-based template switching method to obtain full-length DNA copies of the product flanked by Illumina adapter sequences, followed by sequencing on an Illumina MiSeq to obtain ∼1 million reads for each sample (see Methods). Based on the sequences, we classified the products into four categories, which we refer as Clean Snap Back, Discontinuous Snap Back, Terminal Transferase to Snap Back, and Terminal Transferase (Figure 4C and D). Clean Snap Back products were primed via base pairing between the 3’ A or AA residues of the template and upstream microhomology sites and continued uninterrupted to the 5’ end of the template. Remarkably, even a single A/T base pair was sufficient for priming. Discontinuous Snap Back and Terminal Transferase to Snap Back products were initiated similarly by the 3’ end of the template priming at short upstream microhomologies, but with the former having deletions due to the enzyme skipping over part of the template after the initial priming event and the latter containing insertions of non-templated nucleotides added by terminal transferase activity to the 3’ end of the template prior to annealing to upstream AA residues (Figure 4C). The remaining sequences contained non-coded nucleotides added by the terminal transferase activity to the 3’ end of the template without continuing to snap back DNA synthesis.

Comparison of the sequencing data for different proteins showed that the WT and I/A mutant G2L4 RTs produced the highest proportion of snap back products (>95% of all sequences), with WT G2L4 RT in the absence of Mn^2+^ giving the highest proportion of clean snap back products and the I/A mutation or presence of Mn^2+^ decreasing the proportion of clean snap back products (Figure 4C). WT GII RT produced fewer total and clean snap back products on the DNA substrate, but the A/I substitution increased the proportions of both to levels approaching those of WT G2L4 RT (Figure 4C). These findings show that both G2L4 and GII RTs have snap-back DNA synthesis activity, reflecting the ability to anneal and extend short microhomologies between the 3’ end and upstream regions of DNA templates and that this activity is favored by I at the RT active site.

### G2L4 and GII RT function in microhomology-mediated end-joining *in vitro*

Next, we tested whether the G2L4 and GII RTs could perform MMEJ in a classical DSBR assay using 5’-labeled partially double-stranded DNA substrates with 15-nt single-stranded 3’ overhangs (Figure 5). This assay requires both annealing of short microhomologies at the 3’ end of the singlestranded 3’ overhangs and using the 3’ ends of the base-paired strands as primers to fill in the resulting single-stranded DNA gaps (see schematics in Figure 5A and B). The DNA substrates were incubated with WT or mutant G2L4 and GII RT in the presence of dNTPs in time courses up to 240 min, and the products were analyzed by electrophoresis in a non-denaturing 12% polyacrylamide gel as done above for the snap-back DNA replication products.

**Figure. 5.**
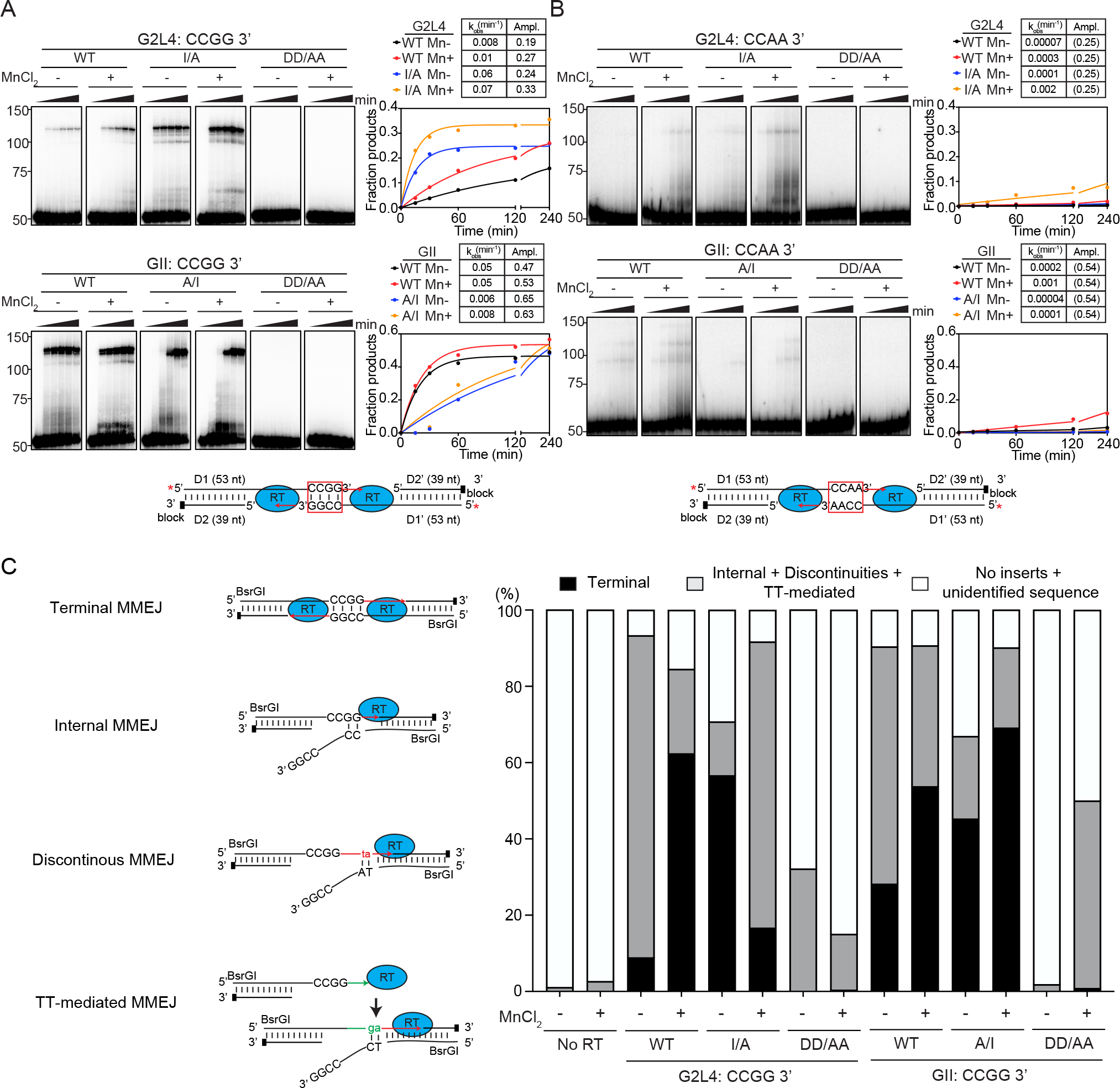
Microhomology-mediated end joining by WT and mutant G2L4 and GII RTs (A and B) Microhomology-mediated end joining (MMEJ) assays. Reactions were done with 500 nM WT or mutant G2L4 and GII RTs and pre-annealed double-stranded DNA substrates (denoted D1/D2 and D1’/D2’ for the left and right sides in the schematics) with 5’-labeled D1 and D1’ and 3’-blocked D2 and D2’) having a 3’ overhangs ending with complementary CCGG (panel A) or non-complementary CCAA (panel B) sequences. The numbers to the left of the gels indicate the positions of size markers in a parallel lane. The plots show the fraction of substrate converted to products running between the 100- and 150-nt size markers. Tables above the plots show rate constants (*k*obs) and amplitudes (Ampl.) obtained by fitting the data to a first-order rate equation. Values of Ampl. in parentheses indicate that the amplitude was fixed at the given value because the reaction did not reach an endpoint during the measurement time (see Methods). (C) High- throughput sequencing of MMEJ products. MMEJ reactions were done as above with 1 µM unlabeled DNA substrates having 3’ overhangs with complementary CCGG sequences for 4 h at 37°C. The products were cloned in *E. coli* using BsrGI sites at the ends of the oligonucleotides, and Illumina DNA sequencing libraries were prepared (see Methods). The products were sequenced on an Illumina MiSeq to obtain ∼1 million 2 x 150 paired end reads for each reaction, and the sequences were aligned by clustalW. Schematics and representative sequences of different categories of products obtained by WT G2L4 RT are shown to the left. Red and green indicate nucleotides at the 3’ end of nascent DNA or added by terminal transferase to the 3’ end of the nascent DNA, respectively, whose annealing to distal regions of the template strand results in discontinuities in DNA synthesis. The stacked bar graphs to the right show the proportions of different products obtained with WT and mutant G2L4 and GII RTs in the absence or presence of 1 mM MnCl2.

In assays with DNA substrates with 3’ overhangs having complementary CCGG-3’ sequences, both the WT G2L4 and I/A mutant RTs gave products of the size expected for MMEJ (∼102 bp, but running higher on the non-denaturing gel), with the rate of product formation higher for the mutant than the WT enzyme, and Mn^2+^ increasing activity of both enzymes (Figure 5A, top). The GII WT and A/I mutant RTs also gave MMEJ products with these substrates, but with the rate of product formation lower for the mutant than the WT enzyme, and Mn^2+^ having relatively little effect on the activity of both enzymes (Figure 5A, bottom). The slower rates of product formation for G2L4 and GII A/I mutant RTs, with I at the active site, likely reflects that primer extension activity is rate-limiting after annealing of the microhomologies. WT GII and G2L4 I/A mutant RTs, the two enzymes with A at the active site, also gave shorter products found by sequencing to result from filling in the single-stranded gap in one rather than both directions (see below). Similar results were obtained with DNA substrates having 3’ overhangs with complementary TTAA-3’ sequences, but with lower rates and amplitudes and different effects of added Mn^2+^ (Figure S12A). By contrast, little or no product was seen for any of the proteins with DNA substrates having 3’ overhangs with non-complementary CCAA-3’ sequences, confirming the dependence of the activity on base pairing of complementary 3’ overhangs (Figure 5B). Human DNA polymerase *θ* prefers MMEJ substrates with a 5’ phosphate on the resected strand (Kent et al., 2015), but this was not the case for the G2L4 or GII RTs (Figure S12B). For both G2L4 and GII RTs, DD/AA mutations abolished detectable MMEJ product formation, indicating that primer extension is required to stabilize the products after annealing of the microhomologies (Figure 5A and B, Figure S12A and B).

For high-throughput sequencing of the MMEJ products, we used DNA substrates with 3’ overhangs having complementary CCGG-3’ sequences. After the *in vitro* reaction, the MMEJ products were ligated into a plasmid via digestion at terminal BsrGI sites and cloned in *E. coli*, a step that enables completion and processing of gapped products, prior to sequencing on an Illumina MiSeq to obtain ∼1 million reads for each sample (see Methods). The results are summarized by stacked bar graphs, which show the proportions of different categories of products illustrated by schematic cartoons (Figure 5C).

As observed previously in similar assays with DNA polymerase *θ* (Kent et al., 2016), the MMEJ products could be divided into two major categories based on the length of the sequenced product: terminal (bidirectional) MMEJ products (82-86 nt), resulting from annealing of the 3’ microhomologies and filling in the single-stranded gaps in both directions, and internal (unidirectional) MMEJ products (<82 nt), resulting from annealing of the 3’ microhomologies and filling in only one of the single-stranded gaps. The shorter products also include those with discontinuities in DNA synthesis or in which 3’ nucleotides added by terminal transferase were used to initiate internally within the 3’ overhangs. WT G2L4 and GII A/I mutant RTs (the two enzymes with I at the active site) in the presence of Mn^2+^ were the combinations that gave the highest proportions of terminal MMEJ products (62-69%), followed by G2L4 I/A mutant RT in the absence of Mn^2+^ and WT GII RT in the presence of Mn^2+^ (Figure 5C). Notably, the DD/AA mutants of both enzymes gave no terminal MMEJ products, but did give internal MMEJ products above the levels of the no RT controls, possibly reflecting that the strand annealing activity of these enzymes can contribute to the formation of MMEJ products even in the absence of RT activity (Figure 5C). Collectively, these findings indicate that I at the active site in combination with Mn^2+^ favors annealing of microhomologies in a manner that enables gap filling via processive DNA synthesis in both directions.

### G2L4 and GII RT use somewhat different MMEJ mechanisms influenced by I/A substitutions at the RT active site

The finding that G2L4 RT differs from GII RT in being unable to efficiently use long (*≥*5 nt) DNA primers in primer extension assays (Figure 3A and B) suggested that the two enzymes might differ in their ability to bind and initiate DNA synthesis from longer annealed microhomologies and/or the extent to which this ability is influenced by a single-stranded region upstream of the microhomology, which is required for efficient MMEJ by human Pol *θ* (Kent et al., 2015, 2016; Black et al., 2019). To test these possibilities, we compared the MMEJ activities of WT and mutant G2L4 RTs using 3’-overhang DNA substrates that differ in the length of the 3’ microhomology and/or the length of the single-stranded gap upstream of the microhomology (Figure S13A-C). Among the substrates tested, the MMEJ activity of WT G2L4 was highest on those with a short (4 bp) microhomology and a long (17 nt) single-stranded gap preceding the microhomology, with a longer microhomology (10 bp) or shorter single-strand gap (6 nt) strongly decreasing activity and with substitution of A at the active site enabling higher activity with longer microhomologies and shorter single-stranded gaps (Figure S13A-C, top panels). By contrast, WT GII RT was better than G2L4 RT in binding and initiating DNA synthesis on substrates with longer microhomologies (10 bp) and shorter (6 nt) single-stranded gaps with the major effect of the A/I mutation being to decrease the rate but not the amplitude of product formation, reflecting the decreased rate of primer extension resulting from this mutation (Figure 13A-C bottom panels for GII RT compared to top panels for G2L4 RT). These results indicate that both G2L4 and GII RT function similarly to DNA polymerase *θ* in binding to single-stranded regions preceding microhomologies (Kent et al., 2016; Black et al., 2019), but with GII RT and the G2L4 I/A mutant, the two enzymes with an A residue at the RT active site, better able to bind directly to and initiate DNA synthesis from longer annealed microhomologies, mirroring the behavior of these proteins with longer annealed primers in primer-extension reactions.

### RT0 loop-dependent strand annealing contributes to MMEJ by G2L4 and GII RTs

The finding that G2L4 and GII RTs function in DSBR by annealing short microhomologies recalled that group II intron and other non-LTR-retroelement RTs have a proficient end-to-end template-switching activity that requires the annealing of short base-pairing interactions between the donor and acceptor nucleic acids (Bibillo and Eickbush, 2004; Mohr et al., 2013). Previous findings showed that this activity is dependent upon the RT0 loop, a distinctive conserved structural feature of non-LTR-retroelement RTs, with deletions in the RT0 loop inhibiting the template-switching activity but not the primer extension activity of both GII and insect R2 element RTs (Jamburuthugoda and Eickbush, 2014; Stamos et al., 2017; Lentzsch et al., 2019). A recent X-ray crystal structure of a template-switching complex of GII RT revealed the structural basis for this activity by showing that the annealing of short base-pairing interactions between the donor and acceptor nucleic acids occurs in a binding pocket that is formed by the RT0 and fingertips loops and is absent in retroviral RTs (Lentzsch et al., 2021).

To investigate if RT0 loop-dependent strand annealing contributes to MMEJ, we constructed a G2L4 mutant (G2L4 *Δ*RT0) in which the RT0 loop (amino acids 23-31) was replaced with a glycine residue and compared its biochemical activities to those of a previously described GII RT-*Δ*RT0 mutant. We found that the ΔRT0 mutants of both enzymes retained high primer extension activity on both DNA and RNA templates, with the RT0 loop mutation surprisingly enabling G2L4 RT to use the longer 20-nt DNA primer almost as efficiently as the 5-nt DNA primer (Figure 6A and B, S14A and B; time courses Figure S14C and D). Notably, the RT0 loop deletion in both RTs strongly inhibited the Mn^2+^-dependent terminal transferase activity on single-stranded DNA, suggesting that this mutation affects the ability to bind the 3’ end of a single-stranded DNA in a position to function as a primer at the RT active site (Figure 6C; time course gels Figure S15A and B). Because the ΔRT0 mutants retain high primer extension activity, MMEJ assays provide a means of assessing the contribution of the RT0 loop to the strand annealing activity used in MMEJ. For both enzymes, the ΔRT0 loop mutations strongly inhibited MMEJ (Figure 6D and Figure S15C for CCGG-3’ and TTAA-3’ microhomologies, respectively), indicating that the presence of the RT0 loop is crucial for the strand annealing activity of both G2L4 and GII RTs.

**Figure 6.**
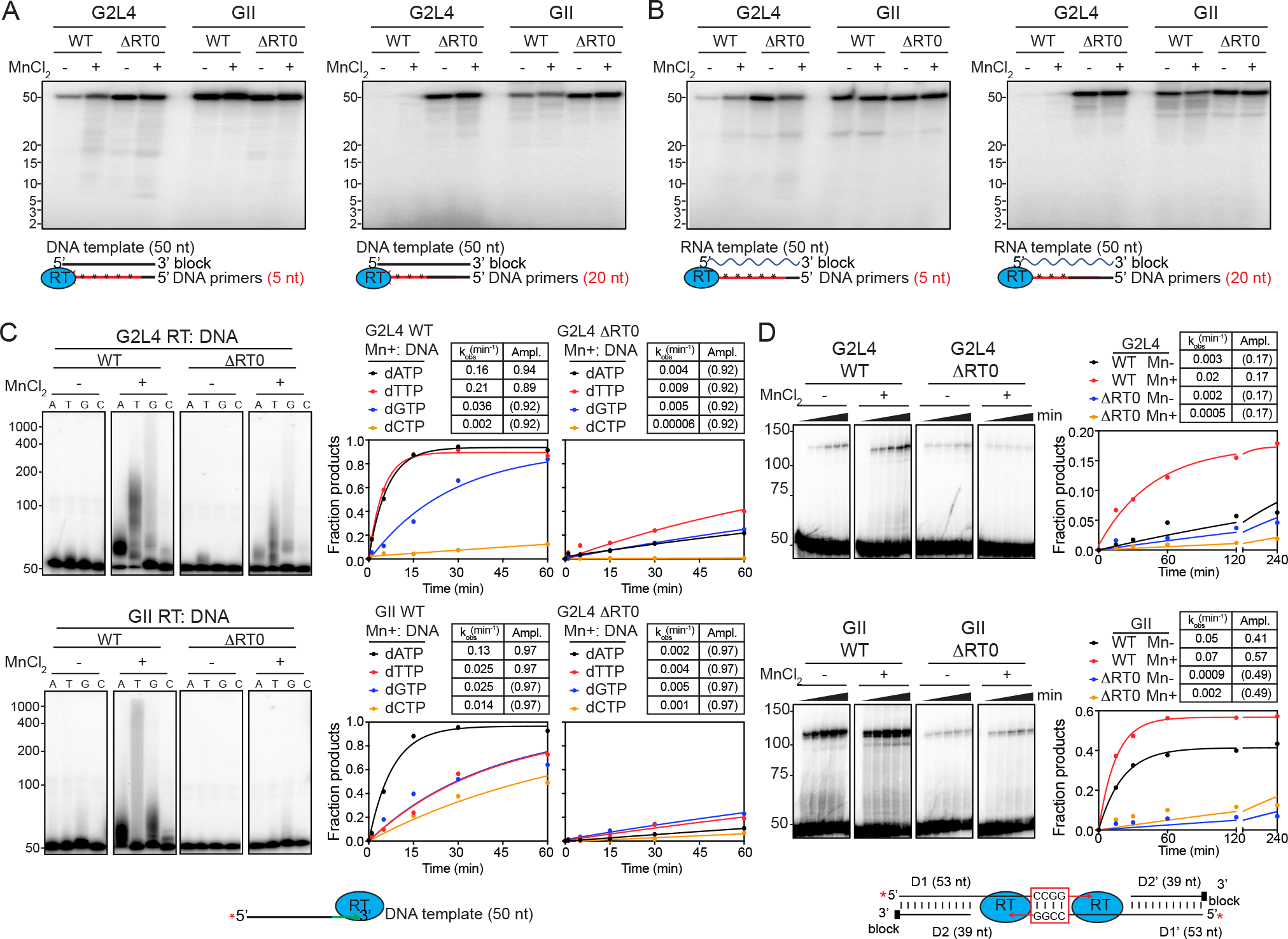
Effect of deletion of the RT0 loop on biochemical activities of G2L4 and GII RTs (A and B) Primer extension. Reactions were done as described in Methods and Figure 3 with 500 nM WT or mutant G2L4 and GII RTs with or without 1 mM MnCl2 for 20 min at 37°C. Time courses are shown in Figure S14. (C) Terminal transferase assays. Reactions were done as described in Methods with 500 nM WT or *Δ*RT0 G2L4 or GII RTs. The gel panels (left) show a single 20-min time point, and the plots to the right show time courses based on quantitation of all labeled products for gels shown in Figure S15A and B. (D) MMEJ assays. Reactions were done as in Figure 5A with WT or *Δ*RT0 G2L4 or GII RTs using DNA substrates having 3’ overhangs ending with a complementary CCGG sequences. Tables above the plots show the rate constants (*k*obs) and amplitudes (Ampl.) obtained from fits by a first-order rate equation. Values of Ampl. in parentheses indicate that the amplitude was fixed at the given value because the reaction did not reach an endpoint during the measurement time (see Methods).

### G2L4 RT repairs double-strand breaks in chromosomal DNA

Finally, to investigate how G2L4 and GII RTs repair DSBs in bacterial chromosomes, we used CRISPR-Cas9 (Chen et al., 2018) to introduce a targeted DSB in the *E. coli thyA* gene, which encodes thymidylate synthase and enables both positive and negative selection for *thyA* mutants (Figure 7A and S16A and B; Sangurdekar et al., 2011). As chromosomal DSBs are lethal in *E. coli*, we first tested whether the expression of WT or mutant G2L4 and GII RTs increases cell survival after co-expression of Cas9 and a guide RNAs directed to introduce a DSB at a site within the *thyA* gene of *E. coli* HMS174 (DE3) *recA*^-^, the same strain used for previous *E. coli* genetic assays (Figure S16C). Surviving bacteria were grown on 2X YT agar plates supplemented with thymine to enable growth of cells with *thyA* gene mutations. The results paralleled those of previous genetic and biochemical assays, with the frequency of surviving bacteria increased 2 to 4-fold by expression of WT G2L4 and GII RTs, to a lesser extent by mutant A/I and GII *Δ*RT0, and not significantly above background by G2L4 *Δ*RT0 or DD/AA mutants of both RTs (Figure 7B). Reciprocally, plating the treated cells on 2X YT + thymine agar plates with added trimethoprim, which selects for cells that harbor a mutant *thyA* gene, showed substantially increased frequencies of *thyA* mutations in cells expressing WT G2L4 and GII RTs and G2L4 I/A mutant RTs, to a lesser extent for GII I/A *Δ*RT0 RT, and not significantly above background for the other mutant RTs (Figure 7C). Western blots showed relatively high expression levels for all G2L4 and GII RT proteins, except for G2L4 *Δ*RT0 and DD/AA, possibly limiting the ability to detect low residual activity of the latter proteins (Figure S16C). These findings show that expression of G2L4 and GII RTs increases cell survival after induction of DSBs and that the surviving cells have increased frequencies of *thyA* mutations, as expected for DNA repair by MMEJ.

**Figure 7.**
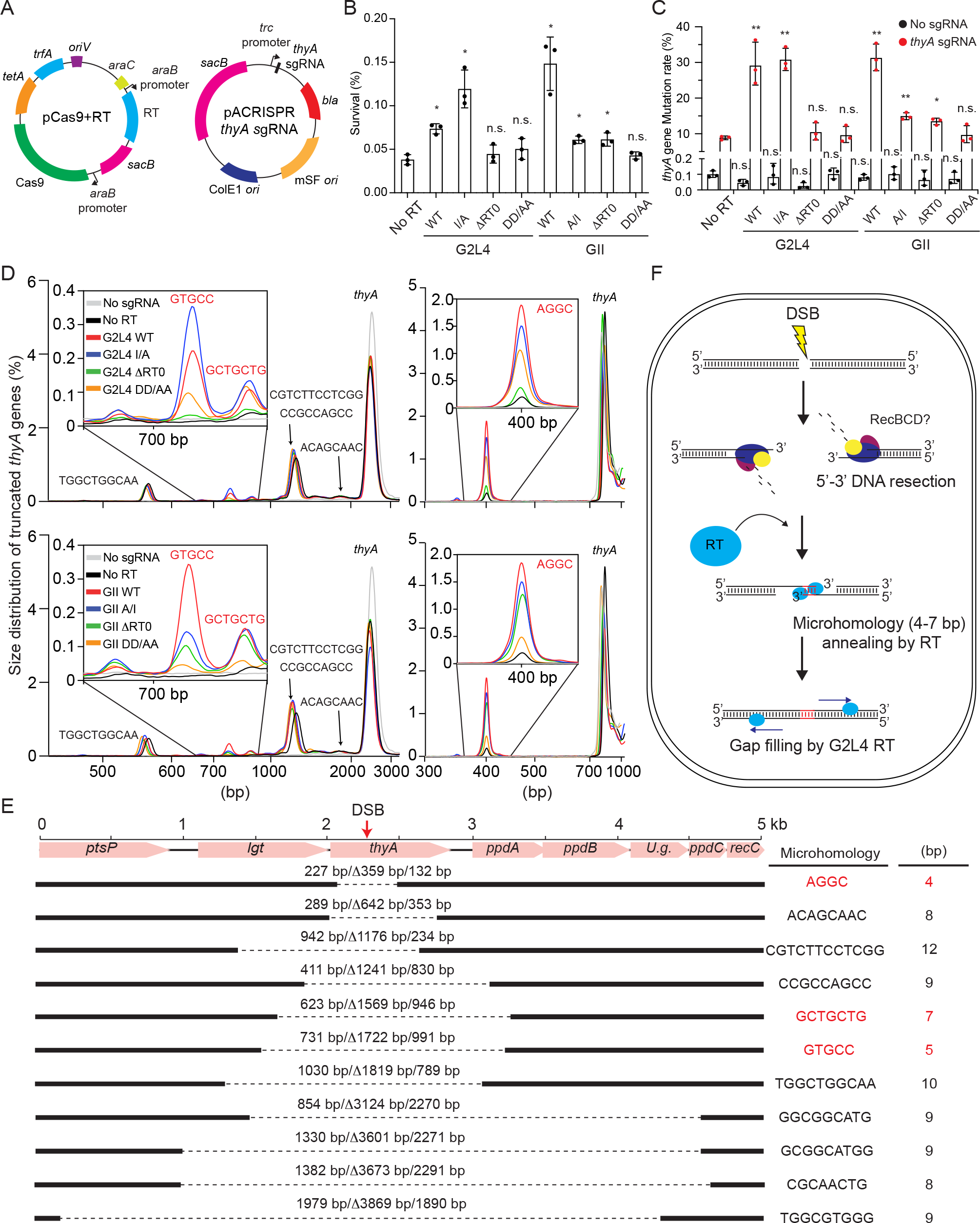
Repair of CRISPR-Cas9-induced double-strand breaks in the *E. coli thyA* gene by WT or mutant G2L4 and GII RTs (A) Schematic of plasmids used in the experiment. pCas9+RT (left) is a derivative of pCasPA that co-expresses Cas9 and G2L4 or GII RTs using separate arabinose-inducible *araB* promoters, and pACRISPR *thyA* sgRNA (right) is a derivative of pACRISPR that expresses *thyA* sgRNA from a constitutive *trc* promoter. *bla*, carbenicillin-resistance gene; *ColE1*, replication origin; *mSF*, a broad-host-range replication origin; *oriV*, replication origin; *sacB*, a counter-selectable marker; *tetA*, tetracycline-resistance gene; *trfA*, initiation of plasmid replication. (B) Cell survival after CRISPR-Cas9 induced DSBs in the *E. coli thyA* gene. Cell survival was measured in plating assays as the percentage of viable *E. coli* HMS174 (DE3) cells on medium containing thymine after introduction of pACRISPR *thyA* sgRNA into cells expressing Cas9 alone or together with WT or mutant G2L4 or GII RTs. (C) Frequency of *thyA* mutants after repair of CRISPR-Cas9 induced DSBs. The percentage of surviving cells with *thyA* mutations was measured in plating assays on medium containing thymine + trimethoprim (200 µg/mL) after introduction of pACRISPR *thyA* sgRNA into cells expressing Cas9 alone or together with WT or mutant G2L4 or GII RTs. The bar graphs in panels B and C show the average values for three repeats of the experiment with the error bars indicating the standard deviation. n.s., non-significant; *, p-value <0.05; **, p-value <0.01. (D) Bioanalyzer tracers of PCR products derived from 750 bp- or 2.5-kb regions encompassing the *thyA* gene in cells expressing WT or mutant G2L4 (top) or GII RTs (bottom) compared to no RT or no guide RNA controls. The bioanalyzer tracers were aligned via the peak corresponding to the full-length *thyA* gene. (E) Sequences of MMEJ products resulting from DSBR repairs in the presence of WT G2L4 RT and co-expression of Cas9 RT and *thyA* sgRNA. The region of the *E. coli* genome encoding the *thyA* gene was amplified by PCR, and PCR products were analyzed by Sanger sequencing. Microhomology sites whose use was increased by RT expression are indicated in red. (F) Model for the RT-CRISPR-Cas9 induced *thyA* gene DSBR. The Cas9-*thyA* sgRNA complex targets and cleaves the *thyA* gene in genomic DNA. After DNA cleavage, the RecBCD complex resects the 5’ strand on opposite sides of the break, resulting single-stranded 3’-DNA overhangs. G2L4 or GII RTs promote MMEJ by annealing microhomologies on opposite sides of the DSB using the 3’ ends of the annealed strands as primers to fill in the single-stranded gaps. The annealed microhomologies are depicted as being at the 3’ end of the 3’ overhangs as in the MMEJ assays reported here, but could also be at other positions within the 3’ overhangs.

To investigate whether DSBR occurred by MMEJ, we PCR amplified and sequenced the genomic region extending up to 2.5-kb upstream and downstream of the *thyA* gene before and after introduction of DSBs by using the *thyA*-directed guide RNA. Bioanalyzer traces showed that expression of the guide RNA resulted in a series of shorter products expected for DSBR after DNA resection resulting in deletions of chromosomal DNA in and around the *thyA* gene. The size distribution of these DSBR products was similar for all RTs, but the relative abundance of a subset of products (shown in insets) was increased substantially by the expression of WT and I/A mutant G2L4 and WT GII RTs and to a lesser extent but still above the no RT control for the *Δ*RT0 and DD/AA mutant RTs (Figure 7D top and bottom panels for G2L4 and GII RTs, respectively).

Sanger sequencing of the PCR products obtained after expression of WT G2L4 and GII RTs showed that those with repaired DSBs had deletions of 359 to 3,869 bp encompassing the region targeted by the guide RNA. All deletions had junction sequences that mapped to short (4 to 12 nt) sequence duplications in the genome. The subset of products enhanced by the G2L4 and GII RTs resulted from annealing of short microhomologies (4 to 7 nt) on either side of the break (Figure 7D and E, sequences shown in red), while those not enhanced by these enzymes resulted from annealing of longer microhomologies (8-12 nt, sequences shown in black), the latter presumably resulting from DSBR by endogenous *E. coli* enzymes. The smaller degree of enhancement at break points with short microhomologies by the *Δ*RT0 mutants RTs is consistent with the *in vitro* assays indicating a requirement for RT0-loop mediated strand anneal activity, and the residual enhancement by the DD/AA mutant RTs is consistent with the strand annealing activity of these enzymes contributing to MMEJ in the absence of RT activity. Collectively, these findings demonstrate that G2L4 and GII RTs function in repairing DSBs in bacterial chromosomes by the MMEJ mechanisms elucidated in the biochemical assays (Figure 7F).

## Discussion

Here we found that G2L4 RT, a free-standing genomically encoded group II intron-like RT, which has a conserved YIDD motif at its active site, functions in DNA damage repair in both its native *P. aeruginosa* host and when transferred into *E. coli*. The biochemical activities used by G2L4 RT for DNA repair include a robust DNA polymerase activity with the ability to read through abasic sites and oxidatively damaged bases, a Mn^2+^-stimulated terminal transferase activity, and the ability to anneal short microhomologies between 3’ overhangs of 5’ strand-resected double-stranded DNAs and use the annealed 3’ DNA ends as primers to fill in the single-stranded gaps. Surprisingly, we found that a group II intron RT with YADD at its active site could function similarly to G2L4 RT in DNA repair *in vitro* and *in vivo*. For both enzymes, a key biochemical activity for DSBR, the ability to anneal short microhomologies between single-stranded 3’-DNA overhangs, was inhibited by mutations in the RT0 loop, a distinctive conserved structural feature of non-LTR-retroelement RTs that is not required for RT activity but was shown previously to function in annealing short microhomologies between donor and acceptor nucleic acids in end-to-end template switching by non-LTR-retroelement RTs (Lentzsch et al., 2019, 2021). These findings and others from the literature discussed below suggest that non-LTR-retroelement RTs may have an inherent ability to function in DSBR in a wide variety of organisms.

Group II intron RTs were shown previously to copy DNA templates, but typically prefer RNA templates in primer extension assays, likely reflecting a steric preference against initiating from B-form DNA template/DNA primer duplexes that fit poorly into the RT active site (Stamos et al., 2017). Similar steric preferences likely contribute to the findings that both G2L4 and GII RT prefer shorter more malleable DNA primers with that preference being particularly stringent for G2L4 RT, which differed from GII RT in being unable to efficiently initiate DNA synthesis from primers

>5 nt (Figure 3). This more stringent preference for shorter primers was governed largely by the non-canonical I residue at the active site, with I/A substitution in G2L4 RT enabling it to use longer primers (Figure 3). Reciprocal substitutions showed that A at the active site enables a higher rate of primer extension and that I at the active site decreases the rate but not the processivity or amplitude of primer extension in both enzymes (Figure 3 and Figure S10).

The ability of both G2L4 and GII RT to read through DNA lesions, such as an abasic site or 8-oxoguanine, was unsurprising in light of previous studies, which showed that group II intron RTs differ from retroviral RTs in being able to read through and distinctively mis-incorporate at RNA post-transcriptional modifications that affect base pairing with the incoming dNTP, enabling GII RT (sold commercially as TGIRT-III) to be used for mapping base modifications in naturally occurring RNAs (Katibah et al., 2014) and DMS-induced modifications in RNA structure mapping (Zubradt et al., 2017). The findings that group II intron and group II intron-like RTs have robust DNA polymerase activity and perform a DNA-based function like DSBR suggest that additional biological functions and biotechnological applications of group II intron-related RTs may be on the horizon.

The combination of biochemical activities used by G2L4 RT for DSBR are remarkably similar to those of human DNA Pol *θ*, which has been shown to have both DNA polymerase activity and limited RT activity, the ability to read through DNA lesions in an error prone manner, a Mn^2+^dependent terminal transferase activity that enables extension of 3’ ends in search of microhomologies, and the ability to anneal short microhomologies between 3’-DNA overhangs and use the annealed 3’ ends as primers to fill in the single-stranded gaps (Seki et al., 2004; Kent et al., 2015, 2016; Black et al., 2019; Chandramouly et al., 2021). The similarities between Pol *θ* and G2L4 extend to the ability to switch between templated and non-template nucleotide addition during DNA synthesis, the ability to bind the 3’ terminus of single stranded DNA, a preference for short rather than long microhomologies, and a requirement for a single-stranded region upstream of the annealed microhomology (Figure S13A-C; Kent et al., 2016; Black et al., 2019). GII RT uses the same biochemical activities as G2L4 RT for DSBR, but is better able to bind and prime DNA synthesis from longer annealed microhomologies, a difference governed by the I/A substitutions at the RT active site (Figure S13 A-C). Our findings are consistent with G2L4 and GII RT using a MMEJ mechanism similar to that proposed for Pol *θ* in which the enzyme binds initially the 3’ end of the single-stranded 3’ overhang on one side of the DSB in the primer-binding site and uses it to search for a complementary microhomology in the opposite 3’ overhang (Kent et al., 2016; Black et al., 2019).

The RT0 loop was shown previously to play a key role in annealing short base-pairing interactions between the donor and acceptor nucleic acids in end-to-end template switching by group II intron and non-LTR-retrotransposon RTs (Jamburuthugoda and Eickbush, 2014; Mohr et al., 2013; Stamos et al., 2017; Lentzsch et al., 2019). Our findings that deletions in the RT0 loop of G2L4 and GII RT inhibit MMEJ without inhibiting primer extension activity (Figure 6 and 7) indicate that the RT0 loops of both enzymes also play a role in strand annealing of microhomologies during MMEJ. Mechanistically, MMEJ and template switching are analogous in requiring the annealing of short microhomologies between two nucleic acid substrates and using the 3’ end of one of the annealed strands as a primer for initiating DNA synthesis on the other. A difference, however, is that end-to-end template switching by group II intron RTs is optimal for annealing of a single base pair, while longer base-pairing interactions are inhibitory (Lentzsch et al., 2019), likely reflecting that the 3’ ends of the donor and acceptor nucleic acid bind after RT core closure in a tightly constrained binding pocket formed by the RT0 loop and the fingertips loop (Lentzsch et al., 2021). By contrast, the annealing of longer microhomologies, such as those typically used for MMEJ, is more akin to the mechanism used for binding and annealing primers for primer extension, as evidenced by the findings that WT G2L4 and GII A/I RT with I at the active site favors the use of shorter primers and microhomologies, while GII and G2L4 I/A RTs with A at the active site enables more efficient use of longer primers or microhomologies (Figure 3A and C, Figure S10, and Figure S13B and C). Although the RT0 loop deletions that inhibit strand annealing of G2L4 and GII RT do not inhibit the primer extension activity of these enzymes, they do inhibit terminal transferase activity (Figure S14 and S15), suggesting that the RT0 loop may be required for binding the 3’ end of the priming strand at the RT active site, potentially a critical first step for strand annealing by these enzymes (Kent et al., 2015).

Although the DNA repair activities of GII RT appear to be as good or better than those of G2L4 RT, mobile group II introns RTs bind group II intron RNAs co-transcriptionally in order to promote RNA splicing and remain tightly bound to the excised the intron RNA to promote reverse splicing into DNA target sites during retrohoming (Yang et al., 1996; Saldanha et al., 1999). Some group II intron RTs can mobilize other group II intron RNAs *in trans*, indicating that they remain functionally active as free proteins and might contribute to DNA repair in their host cells (Brown et al., 2014; Meng et al., 2005; Mohr et al., 2010; Zoschke et al., 2010). Other group II intron RTs, however, might be sequestered to at least some degree by binding to the intron RNA co-transcriptionally (Gu et al., 2010) and thus prevented from functioning in DNA repair in their host cells.

A likely evolutionary scenario is that free-standing bacterial RTs that perform host functions evolved from the RT of a mobile group II intron that integrated into a bacterial genome and became immobilized by mutations in the intron RNA. Because group II intron mobility is deleterious to the host cell, mutations that immobilize the intron RNA are favored by purifying selection resulting in numerous examples of mobility-compromised group II introns that remain integrated in bacterial genomes (Leclercq and Cordaux, 2012; Mohr et al., 2010; Robart and Zimmerly, 2005). After acquiring a host function that contributes to cell survival, the RT would be subject to positive selection to retain and enhance its ability to perform that function. Group II intron RTs that integrated into a pathogenicity island or became associated with other mobile elements could be disseminated to other bacteria, as suggested here for G2L4 RT (Figure S1).

In the case of G2L4 RT, the potential to function in DSBR may have pre-existed in the ancestral group II intron RT, with subsequent dissociation from the intron RNA enabling the protein to evolve to better perform that function with less constraint on biochemical activities that are required for intron mobility. Our results indicate that the substitution of an I residue for an A residue at the RT active site was a key adaptation that enabled G2L4 RT to better perform its host function in DSBR by favoring MMEJ at the expense of primer extension activity. The finding that the reciprocal A/I substitution in GII RT strongly inhibited primer extension activity suggests that additional changes were needed to accommodate the bulkier I at the active site with less effect on primer extension activity. A number of other group II intron-like bacterial RTs that evolved to perform host functions also have conserved substitutions in the F/YxDD motif at the RT active site, which may likewise enable them to better perform their host function, including M in diversity generating retroelement RTs associated with a higher frequency of nucleotide substitutions (Wu et al., 2018).

Finally, the close structural similarity between group II intron and other non-LTR-retroelement RTs and the finding that MMEJ by these RTs is dependent upon the RT0 loop, a conserved structural feature of non-LTR-retroelement RTs that is absent in retroviral RTs, suggests that nonLTR-retroelement RTs may have an inherent ability to function in DSBR in a wide variety of organisms. A driving force for the evolution of such an activity in non-LTR-retroelement RTs is suggested by the finding that the ability of both a bacterial group II intron and human LINE-1 element to proliferate to higher copy numbers in bacterial cells correlates with the ability of the bacterial host strain to repair DSBs, which are a side product of the retromobility of these elements (Lee et al., 2018). Numerous previous findings have shown that human LINE-1 elements have a close personal relationship with DSBs, including inducing them during retrotransposition and contributing to RNA-mediated DSB repair by using both LINE-1 endonuclease-induced and spontaneous DSB sites for templated insertions of processed pseudogenes and other cDNAs (Esnault et al., 2000; Morrish et al., 2002, 2007; Onozawa et al., 2014). Our findings extend the previously known connections between LINE-1 and other non-LTR-retrotransposon elements with DSBs by suggesting that non-LTR-retrotransposon RTs may function not only in producing cDNAs that are integrated at DSBs, but may also play an active role in repairing DSBs by mechanisms similar to those elucidated here for G2L4 and GII RTs. In this way, human LINE-1 and other non-LTR retroelement RTs may not only mitigate damage caused by their retrotransposition, but also provide a benefit to their host organisms in exchange for proliferating within their genomes.

## ACKNOWLEGMENTS

This work was supported by Welch Foundation grant F-1607 and NIH grant R35 GM136216 to AML. RR is supported by NIH grant R35 GM131777. We thank Kyle M. Miller (University of Texas at Austin) for use of his X-ray generator, Jennifer L. Stamos (University of Texas at Austin) for comments on the manuscript, and Douglas C. Wu (Invitae) for help with whole genome sequence analysis. The authors acknowledge the Texas Advanced Computing Center (TACC) at the University of Texas at Austin for providing high performance computing resources that have contributed to the research results reported in this paper (URL: http://www.tacc.utexas.edu).

## METHODS

### Bacterial strains

*Pseudomonas aeruginosa* AZPAE12409, which is naturally resistant to chloramphenicol (Cap^R^), was obtained from Entasis Therapeutics (Kos et al., 2015). *E. coli* HMS174 (DE3) (F^-^ *recA1 hsdR*(rK12^-^ mK12^+^) *Rif* ^R^ (DE3)) was purchased from Novagen. *E. coli* S17.1 (*recA pro hsdR* RP42-Tc::Mu-Km::Tn7 integrated into the chromosome Str^R^; Spc^R^; Tmp^R^) was purchased form ATCC. *E. coli* Rosetta 2 (F^-^ *ompT hsdS*B(rB^-^ mB^-^) *gal dcm* pRARE2 (Cap^R^)) and Rosetta 2 (DE3) (F^-^ *ompT hsdS*B(rB^-^ mB^-^) *gal dcm* (DE3) pRARE2 (Cap^R^)) were purchased from Novagen.

### DNA and RNA oligonucleotides

The DNA and RNA oligonucleotides used in this study are listed in Table S2. All were purchased in RNase-free, HPLC or PAGE-purified form from Integrated DNA Technologies (IDT). Oligonucleotides were 5’-labeled with [*γ*-^32^P]-ATP (6,000 Ci/mmol; Perkin Elmer) by using T4 polynucleotide kinase (New England Biolabs) and cleaned up by using an Oligo Clean & Concentrator or RNA Clean & Concentrator Kit (Zymo Research), all according to the manufacturers’ protocols.

### Recombinant plasmids

Recombinant plasmids used in this study are listed in Table S3. The targetron expression plasmid pBL1 is a derivative of the broad host range expression vector pJB866 (Blatny et al., 1997), which expresses the Ll.LtrB-*Δ*ORF targetron using an m-toluic acid-inducible promoter and carries a Tet^R^ marker (Yao and Lambowitz, 2007).

pBL1-MCS is a derivative of pBL1 used as an intermediate in the construction of plasmids that express G2L4 and GII RTs in *P. aeruginosa* and *E. coli*. It was derived from pBL1 by replacing the 3-kb XhoI + KpnI fragment containing the targetron cassette with a 42-nt DNA segment containing a multi-cloning site (KpnI/SpeI/BamHI/HindIII/BsrGI/XhoI) oligonucleotide synthesized by IDT.

pBL1-MBP-RT-8XHis plasmids used for expressing WT and mutant G2L4 and GII RTs in *P. aeruginosa* and *E. coli* were constructed by PCR amplifying the RT ORFs from pMal-RT plasmids (see below) with primers that introduce flanking KpnI and SpeI sites and then cloning the resulting ∼2.4-kb PCR products between the KpnI and SpeI sites of pBL1-MCS. The direct repeat region upstream of the G2L4 RT ORF in *P. aeruginosa* AZPAE12409 was inserted into pBL1MBPG2L4 RT-8XHis by PCR amplifying a 658-bp region of genomic DNA containing the Direct Repeats with Gibson forward and reverse primers that append flanking KpnI sites and inserting the KpnI-digested PCR product into the KpnI site of pBL1-MBPG2L4 RT-8XHis by using NEBuilder HiFi DNA Assembly (New England Biolabs) according to the manufacturer’s protocol. The proteins expressed from these plasmids have an N-terminal maltose-binding protein (MBP) tag, which stabilizes and increases the solubility of expressed group II intron RTs (Mohr et al., 2013), and a C-terminal 8XHis tag used for detection by immunoblotting.

pMal-RT plasmids used to express G2L4 and GII RTs in *E. coli* for protein purification were derivatives of pMal-c5X (New England Biolabs), which carries an Amp^R^ marker and uses an IPTG-inducible *tac* promoter to expresses recombinant proteins with a factor Xa cleavable maltose-binding protein (MBP) tag. pMal-GII RT WT and GII RT 23-41/4G (denoted GII RT *Δ*RT0) were described previously (Mohr et al., 2013; Stamos et al., 2017). pMal-G2L4 RT was constructed by cloning a G-Block containing a codon-optimized G2L4 RT ORF flanked by HpaI and BamHI sites (IDT) between the XmnI and BamHI sites of pMal-c5X. Other G2L4 and GII RT mutant plasmids were derived from pMal-G2L4 RT or pMal-GII RT by using a Q5 mutagenesis kit (New England Biolabs).

pKS-SacB used for cloning *in vitro* MMEJ products (Figure 5) is a derivative of pBluescriptII KS(+) (Agilent), which was constructed by PCR amplifying the *sacB* gene of pACRISPR (Addgene plasmid #113348) (Chen et al., 2018) with SacB forward and reverse primers that introduce flanking EcoRV sites, and then cloning the resulting PCR Product into the EcoRV site of pBluescriptII KS(+).

pCas9+RT ORF plasmids used for CRISPR-Cas9 *in vivo* DSBR assays in *E. coli* were constructed via an intermediate plasmid pCas9SX derived by replacing the lambda red ORF of pCasPA (Addgene #113347) (Chen et al., 2018) with a 47-bp DNA region with flanking SpeI and XbaI sites by using a Q5 mutagenesis kit (New England Biolabs). MBP-RT-8XHis ORFs were inserted into pCas9SX by PCR amplifying the pBL1-MBP-RT-8XHis ORFs with primers that introduce flanking SpeI and XbaI sites and cloning the PCR product between the corresponding sites of pCas9SX.

pACRISPR *thyA* sgRNA expressing plasmids were constructed as described (Chen et al., 2018). A sgRNA for the *thyA* gene was designed with on-line tools (http://chopchop.cbu.uib.no) and the corresponding DNA sequence was inserted as synthetic oligonucleotide into pACRISPR (IDT).

All insertions and PCR amplified regions of plasmid used in this study were confirmed by Sanger sequencing.

### Bioinformatic analysis of G2L4 RT ORF in AZPAE12409 *P. aeruginosa*

At the outset of this study, we used the protein sequence of a G2L4 RT (ABB74237) from *Nitrospira multiformis* ATCC 25196 to search Genbank using BLASTP (Altschul, 1997) and identified >100 G2L4 RT ORFs in gram negative *α*, *β*, *γ* and a few *δ* proteobacteriales from which a G2L4 protein (WP_034031052) from *P. aeruginosa* strain AZPAE12409 was selected for further analysis. More recent Genbank searches revealed ∼240 unique G2L4 RT proteins and a total of ∼500 G2L4 RTs including identical proteins in different strains. Analysis of the genomic neighborhood of G2L4 RTs with MUMmer3 (Kurtz et al., 2004) revealed two ∼140 bp direct repeats within 1kb upstream of the G2L4 RT ORF in ∼70% of the sequences (Supplemental File).

The GC content across the region of the *P. aeruginosa* AZPAE12409 genome containing the G2L4 RT ORF (Figure S1) was calculated across a 500-bp sliding window. The number of rare codons in the G2L4 RT and neighboring ORFs was determined from the *P. aeruginosa* PAO1 codon table (NCBI) and defined as codons that occur less than once per 1,000 codons. Promoters were predicted by using BacPP (bacterial promoter prediction, http://www.bacpp.bioinfoucs.com). The secondary structure of G2L4 RT was predicted by using HHpred (https://toolkit.tuebingen.mpg.de/tools/hhpred).

### Targetron gene knock-out of G2L4 RT in *P. aeruginosa*

Targetron disruption of the G2L4 RT ORF in *P. aeruginosa* AZPAE12409 was done by using the broad-host range targetron expression vector pBL1 with targetrons designed and constructed as described (https://sites.cns.utexas.edu/lambowitz/targetron-design; Yao and Lambowitz, 2007). Targetron plasmids were transformed into *E. coli* S17.1 and introduced into *P. aeruginosa* AZPAE12409 via conjugation. For this purpose, the *P. aeruginosa* recipient and *E. coli* donor carrying the pBL1 Tet^R^ targetron construct were grown separately in 50-mL conical tubes (Sarstedt) containing 5-mL Luria Bertani (LB) medium with tetracycline (25 μg/mL) added for the *E. coli* culture and shaken (250 rpm) at 37°C until O.D.600 = 0.3-0.4. The *P. aeruginosa* and *E. coli* cultures were then mixed at a 1:10 ratio, and cells were collected by filtration on a 25-mm diameter membrane filter (0.45-µm pore size; Millipore). For conjugation, the membrane was placed on a LB agar plate for 3 h at room temperature and then transferred to 5-mL of LB medium in a 50-mL conical tube (Sarstedt) and vortexed vigorously to separate the conjugating cells. Aliquots were plated on a LB agar plate containing tetracycline (100 μg/mL) and chloramphenicol (25 μg/mL) to which the *P. aeruginosa* strain is naturally resistant and incubated at 37°C for 14-16 h to select *P. aeruginosa* colonies carrying the Tet^R^ targetron plasmid. A single *P. aeruginosa* colony containing the targetron plasmid was picked and grown in LB medium containing tetracycline (100 μg/mL) overnight at 37°C. The culture was then diluted 1:100 into 5-mL LB medium plus tetracycline (100 μg/mL) in a 50-mL conical tube (Sarstedt) and incubated at 37°C with shaking (250 rpm) until O.D.600 = 0.3-0.4, at which point 2 mM m-toluic acid was added to induce targetron expression. After incubating at 30°C without shaking overnight, cells were plated on LB agar containing tetracycline (100 μg/mL), and disruptants were identified by colony PCR using primers flanking the predicted targetron insertion site in the G2L4 RT ORF. Twelve colonies were picked of which two (KO1 and KO2) contained the targetron insertion. Disruptants were confirmed by Southern hybridization and genomic sequencing on an Illumina MiSeq to obtain ∼1 million, 2 x 250 paired end reads at the Genome Sequencing and Analysis Facility (GSAF) at the University of Texas at Austin.

### Southern hybridization

Genomic DNA was isolated from WT and G2L4 knock-out *P. aeruginosa* AZPAE12409 by using a Quick-DNA Fungal/Bacterial Miniprep Kit (Zymo Research) according to the manufacturer’s protocol. The DNA was digested with PstI and EcoRI and run in a 1% agarose gel alongside a 1-kb Plus DNA Ladder (Invitrogen) that was 5’-labeled with [*γ*-^32^P]-ATP (6000 Ci/mmol; Perkin Elmer) usingT4 polynucleotide kinase (New England Biolabs). After electrophoresis, the gel was blotted onto an Amersham Hybond-XL (Cytiva) membrane by overnight capillary transfer. The membrane was washed 3 times with 25 mL of 6x SSC, dried, and UV irradiated to cross-link the DNA to the membrane (120 mJ; Stratalinker UV Crosslinker 2400). Hybridization was done with a 5’-labeled targetron probe (200 bp PCR product obtained using G2L4 RT targetron probe primers; Table S2) in a hybridization tube with Amersham Rapid-hybridization Buffer (Cytiva) for 2.5 h at 60°C. After washing twice with 2x SSC plus 0.1 % SDS, the membrane was dried and scanned with a phosphorimager (Typhoon FLA 9500; GE Healthcare).

### Genomic DNA sequence analysis of *P. aeruginosa* AZPAE12409 WT and G2L4 KO strains

Glycerol stocks of *P. aeruginosa* WT and G2L4 RT knock-out strains were inoculated into 5-mL LB medium in a 50-mL conical tube (Sarstedt) and incubated at 37°C for 16-18 h with shaking (200 rpm). The culture was then centrifuged at 4000 x g for 5 min, and genomic DNAs were extracted by using a Monarch Genomic DNA Purification Kit (New England Biolabs) according to the manufacturer’s protocol. 1 µg of each genomic DNA was submitted to the Genome Sequencing and Analysis Facility (GSAF) at the University of Texas at Austin and sequencing libraries were prepared and sequenced on an Illumina MiSeq V2 instrument to obtain ∼1 million 2 x 250 nt paired end reads per sample. Reads were mapped to the customized *P. aeruginosa* AZPAE12409 reference genome, which contains the targetron inserted at the designated location and the pBL1 vector used to express the targetron, using BWA with the default settings (Li and Durbin, 2010). The genomic DNA coverage was calculated as mean coverage of 500-bp bins along the genomic sequences and plotted using R. Variants were called using freeBayes on bam files from genomic alignment of the WT or KO dataset, with the following settings: --ploidy 1 --min-mapping-quality 30 --min-alternate-count 10 (Garrison and Marth, 2012). The statistic test on KO-specific variants (point mutations) against the WT was analyzed by VarScan (Koboldt et al., 2009).

### P. aeruginosa growth curves

Glycerol stocks of *P. aeruginosa* WT and G2L4 RT knock-out strains were streaked on LB agar and incubated at 37°C overnight. The next day, a single colony was inoculated into 5-mL LB medium in a 50-mL conical tube (Sarstedt) and incubated at 37°C overnight with shaking (200 rpm). A 1-mL aliquot of the overnight culture was then added to 100-mL LB in a 250-mL Erlenmeyer flask and incubated at 37°C with shaking (200 rpm). 0.5-mL samples of *P. aeruginosa* WT and G2L4 RT KO cultures were collected every 6 h for up to 72 h, serially diluted, and plated on LB agar. The plates were incubated overnight at 37°C, and colonies were counted to calculate colony forming units (CFUs).

### TGIRT-seq of *P. aeruginosa* WT and G2L4 RT disruptant whole-cell RNAs

*P. aeruginosa* WT and G2L4 RT knock-out strains were grown as described above, and 500 µL samples were collected at 15 and 30 h corresponding to log and mid-stationary phase, respectively. Total cellular RNA was extracted by using a Monarch Total RNA Miniprep kit, and rRNA-depleted by using riboPOOL (siTOOLs biotech), both according to the manufacturer’s protocols. After clean-up using an RNA Clean & Concentrator Kit (Zymo Research), the RNA was fragmented at 95°C for 5 min by using a Next Magnesium RNA Fragmentation Module (New England Biolabs), and cleaned up by using a MinElute PCR Purification Kit (Qiagen). TGIRT-seq libraries were prepared as described (Xu et al., 2019, 2021), and a 1-µL aliquot was analyzed on an Agilent 2100 Bioanalyzer using a High Sensitivity DNA kit to assess quality and concentration. The TGIRT-seq libraries were sequenced via Illumina NextSeq500 to obtain ∼20 million 2 x 75 bp paired end reads per sample at the University of Texas MD Anderson Cancer Center, Science Park.

Reads were mapped to both a *P. aeruginosa* AZPAE12409 reference genome, which was incomplete and computationally curated with only limited information about predicted genes, and to the model *P. aeruginosa* strain PAO1 reference genome, which was complete and had detailed gene annotation (Pseudomonas Genome Database) (Winsor et al., 2011). For read mapping, Illumina TruSeq adapters and PCR primer sequences were trimmed from the reads with Cutadapt v3.2 (sequencing quality score cut-off at 20; p-value <0.01), and reads <15-nt after trimming were discarded (Martin, 2011). The processed reads were mapped separately to the reference genomes for each of the *P. aeruginosa* strains by using Bowtie 2 v2.2.5 with local alignment (settings: --local -N 1 -D 20 -L 20 -X 1000 --no-mixed --no-discordant) and intersected with *P. aeruginosa* PAO1 and AZPAE12409 gene annotations by BEDTools v2.29.2 (Langmead and Salzberg, 2012; Quinlan, 2014). Finally, gene counts from the two *P. aeruginosa* strains were combined by using a customized R script. If a read pair mapped only to a PAO1 or AZPAE12409 gene, the gene annotation of the mapped strain was used, but if a read pair mapped to both the PAO1 and AZPAE12409 strains, the more complete gene annotation of the PAO1 strain was used.

Differential gene expression was analyzed by using DESeq2 and volcano plots were plotted using R (Love et al., 2014). GO term enrichment analysis was done by using the goseq package in R, and heatmaps were plotted by using the pheatmap package in R (https://CRAN.R-pro-ject.org/package=pheatmap) (Young et al., 2010). Coverage plots and read alignments were created by using Integrative Genomics Viewer v2.6.2 (IGV). Genes with >100 mapped reads were down-sampled to 100 mapped reads for visualization in IGV (Robinson et al., 2011).

### *P. aeruginosa* and *E. coli* cell survival assays

*P. aeruginosa* AZPAE12409 WT and KO strains, which had been electro-transformed with pBL1MBP-RT-8XHis plasmids or vector controls, were plated on LB medium containing tetracycline (100 μg/mL) and incubated overnight at 37°C. A single colony was picked and grown in LB containing tetracycline (100 μg/mL) overnight with shaking (200 rpm) at 37°C. The culture was then diluted 1:100 into 5-mL LB with tetracycline (100 μg/mL) in a 50-mL conical tube (Sarstedt) and incubated at 37°C with shaking (200 rpm) until O.D.600 = 1.0, at which point G2L4 RT expression was induced with m-toluic acid (2 mM final) for 2 h with shaking (200 rpm) at 37°C. *P. aeruginosa* WT and G2L4 RT KO strains lacking pBL1 were growth in the same medium as pBL1-containing strains with tetracycline until O.D.600 = 0.5-0.6. The *P. aeruginosa* WT and G2L4 RT KO strains or strains expressing WT or mutant G2L4 RTs from plasmids (see above) were diluted at 1:100 ratio into M63 minimal medium (22 mM KH2PO4, 40 mM K2HPO4, 15 mM (NH4)2SO4) supplemented with 2 mM MgSO4, 0.2% glucose and 0.5% casamino acids for cell survival assays describe below.

*E. coli* HMS174 (DE3) cells, which had been transformed with pBL1-MBP-RT-8XHis plasmids or vector controls, were processed similarly for cell survival assays except that tetracycline concentration was 25 μg/mL, protein expression was induced with m-toluic acid (2 mM final) at 18°C for 19-21 h with shaking (100 rpm), and after induction cells were diluted 1:100 in modified M9 minimal medium (33.7 mM Na2HPO4, 22 mM KH2PO4, 8.55 mM NaCl, 9.35 mM NH4Cl) supplemented with 0.4 % glucose, 2 mM MgSO4, 2 mM MgCl2, 0.1 mM CaCl2, 1 µg/mL thiamine, 2 mM m-toluic acid with or without 0.5 mM MnCl2.

For X-ray irradiation assays, 0.5 mL of the *P. aeruginosa* cells that had been diluted into M63 minimal medium were pipetted into single wells in a 24-well plate (Falcon) and exposed to 35 Gy X-rays using a 43855D RX-650 X-Ray Generator (Faxitron) according to the manufacturer’s protocol, while a second control plate containing *P. aeruginosa* cells was not irradiated. The X-ray irradiated and non-irradiated control cells were serially diluted in M63 medium and plated on LB agar plates for *P. aeruginosa* WT and G2L4 RT KO strains or LB agar plates containing tetracycline (100 µg/mL) for *P. aeruginosa* strains containing pBL1-MBP-RT-8XHis plasmids expressing WT or mutant G2L4 RTs.

For chemical cell survival assays, 1-mL of *P. aeruginosa* or *E. coli* cells diluted in M63 minimal medium as above were incubated with or without 1.5 mM hydrogen peroxide (Sigma-Aldrich) or 60 µg/mL phleomycin (InvivoGen) in 15-mL tubes (Sarstedt) for 1.5 h at 37°C with shaking (250 rpm). The cells were then serially diluted into minimal medium (M63 for P. aeruginosa and M9 for *E. coli*), and plated on LB agar plates or LB agar plates containing tetracycline (100 µg/mL for *P. aeruginosa* or 25 µg/mL for *E. coli*). After overnight incubation at 37°C, colonies were counted, and survival determined as the proportion of colonies surviving after DNA damage compared to untreated controls. p-values were calculated by student’s unpaired t-test in Prism 9.0 (GraphPad Software).

### Immunoblotting

Immunoblot analysis was done with parallel cultures to those used for cell survival assays. Instead of diluting into minimal medium for the cell survival assays, 5 mL of *P. aeruginosa* or *E. coli* HMS174 (DE3) cells expressing wild-type and mutant G2L4 and GII RTs in LB medium were centrifuged at 4°C, 4,000 x g for 10 min, and the pellets were lysed by resuspending in 300-µL of lysis buffer (20 mM Tris-HCl pH 7.5, 500 mM NaCl, 0.1% Triton X-100 and 20% glycerol). The lysed cells were transferred to a 1.5-mL microcentrifuge tube and sonicated three times for 5 sec each time at 30% amplitude using a Branson Sonifier 250 (Branson Ultrasonics) followed by centrifugation at 4°C, 15,500 x g for 15 min. Protein concentrations in the lysates were measured by Quick Start Bradford Protein Assay Kit (Bio-Rad) and SmartSpec Plus Spectrophotometer (Bio-Rad) according to manufacturer’s protocols, and 75 µL of the supernatant was transferred to a 1.5-mL tube, mixed with 25 µL 4X sample buffer (200 mM Tris-HCl pH 6.8, 400 mM DTT, 8 % SDS, 6 mM bromophenol blue, 40% glycerol), and incubated at 95°C for 5 min. Protein samples (50 µg) and Color Prestained Protein Standard, Broad Range ladder (10-250 kDa) (New England Biolabs) were loaded on a NuPAGE 4-12 % Bis-Tris gel, and electrophoresis was done in 1X MES running buffer (Thermo Fisher Scientific) at 150 V for 1 h by using an XCell Surelock Electrophoresis Cell according to the manufacturer’s protocol. For membrane transfer, an Immuno-Blot PVDF Membrane (Bio-Rad) was pre-soaked for 30 sec in 100% methanol, and membrane transfer was performed in 1X NuPAGE Transfer Buffer (Thermo Fisher Scientific) by using a Xcell II Blot Module according to the manufacturer’s protocol. The membrane was blocked by incubating in 15 mL of blocking solution (5% Blotting Grade Blocker Non-Fat Dry Milk (Bio-Rad) in 20 mM Tris-HCl pH 7.5, 150 mM NaCl, 0.1% Tween-20) in a rectangular tray at 4°C on an orbital shaker (55 rpm) for 1 h. For primary antibody treatment, 15 mL 6x His-Tag Monoclonal antibody (MA1-21315; Invitrogen) diluted 1:1500 in blocking solution supplemented with 0.1% NaN3 was added to the membrane and incubated at 4°C on an orbital shaker at 55 rpm for 14-16 h. The membrane was then washed three times with 15 mL 1X TBS-T with shaking (55 rpm) for 10 min at room temperature. For secondary antibody treatment, the membrane was incubated with Donkey antiMouse IgG (H+L) Cross-Adsorbed Secondary Antibody, HRP (SA1-100; Invitrogen) diluted 1:5000 in 1X TBS-T with shaking (55 rpm) for 1 h at room temperature, followed by three washes with 1X TBS-T. The antibody-treated membrane was then incubated with 15 mL Clarity Western ECL Substrate (Bio-Rad) at room temperature with shaking (55 rpm) for 5 min and exposed to CL-Xposure Film (Thermo Fisher Scientific), which was then developed with an X-ray film processor (Konica Minolta SRX-101A).

### Protein purification

Recombinant proteins used for biochemical assays were expressed from pMal-RT plasmids (see above and Table S3. For each protein preparation, *E. coli* Rossetta2 Cap^R^ cells (Novagen) containing freshly transformed expression constructs were plated on LB agar containing ampicillin (100 µg/mL) and chloramphenicol (25 µg/mL) and incubated at 37°C for 14-16 h. A single colony was inoculated into 20 mL of LB containing the same antibiotics and incubated in a 50-mL conical tube (Sarstedt) at 37 °C with shaking (250 rpm) for 14-16 h, then diluted 1:50 into 1 L of LB with the same antibiotics in a 4 L Erlenmeyer flask and incubated at 37°C with shaking (220 rpm) until the O.D.600 reached 0.8-1.0. Protein expression was induced by adding Isopropyl β-D-1-thiogalactopyranoside (IPTG) (100 µM for G2L4 RT constructs and 1 mM for GII RT constructs), followed by incubation at 18°C with shaking (100 rpm) for 19-21 hr. Bacteria were harvested at 4,000 x g for 25 min in a JLA-8.1000 rotor in an Avanti J-E centrifuge (Beckman), and the pellet was transferred to a 50-mL conical tube (Sarstedt) and resuspended in 45 mL of lysis buffer containing 20 mM Tris-HCl pH 7.5, 500 mM NaCl, 0.1% Triton X-100 and 20% glycerol. The resuspended cells were sonicated at 80% amplitude for three 1 min intervals with 1 min pause between bursts using a Branson Sonifier 250 (Branson Ultrasonics) followed by centrifugation at 15,500 x g for 25 min in a JA 25.50 rotor in an Avanti J-E centrifuge (Beckman). The supernatant was transferred to a 50-mL conical tube, and polyethyleneimine (final concentration of 0.04%) was added, mixed by inverting 2-3 times, and incubated on ice for 10 min to precipitate nucleic acids. Precipitates were removed by centrifugation as above, and the supernatant was filtered through a 0.2-µm pore-size nylon membrane (Fisher). The filtrate was loaded onto a 5-mL HiTrap MBP HP column (Cytiva) at a flow rate of 5 mL/min using an ÅKTA START FPLC (Cytiva). The column was washed with 5 column volumes of buffer A (20 mM Tris-HCl pH 7.5, 100 mM NaCl, 0.1% β-mercaptoethanol and 10% glycerol) followed by 5 column volumes of buffer B (20 mM Tris-HCl pH 7.5, 1.5 M NaCl, 0.1% Triton X-100, 0.1 % β-mercaptoethanol, 10% glycerol), and then again with 5 column volumes of buffer A. The protein was eluted from the column with 10 column volumes of elution buffer containing 20 mM Tris-HCl pH 7.5, 100 mM NaCl, 0.1% β-mercaptoethanol, 10% glycerol and 10 mM maltose. 2 mL fractions were collected. 15-µL of each column fraction was mixed with 5-µL 4X sample buffer and loaded on a NuPAGE 4-12% Bis-Tris gel and gel electrophoresis was performed as described above for Immunoblotting. The gel was stained with 0.25% Coomassie brilliant blue R (Sigma-Aldrich) to identify recombinant proteins. Peak fractions were pooled and loaded onto a 5-mL HiTrap Heparin HP column (Cytiva) at a flow rate of 5 mL/min using an ÅKTA START FPLC (Cytiva). The column was washed with 5 column volumes of buffer A. Bound proteins were eluted using 10 column volumes of a 0.1 to 1.5 M NaCl gradient collecting 2 mL fractions. Column fractions containing the protein were identified by SDS-PAGE and Coomassie staining (see above). Fractions containing protein were pooled and concentrated to 10 µM into storage buffer (20 mM Tris-HCl pH 7.5, 50 mM NaCl, 50% glycerol for G2L4 RTs and 20 mM Tris-HCl pH 7.5, 500 mM KCl, 50 % glycerol for GII RTs) with an Amicon Ultra-15 (30k) concentrator (Millipore) according to the manufacturer’s protocol.

### Primer extension assays

The templates for primer extension assays were 50-nt DNA or RNA oligonucleotides (see Table S2) with 3’-ends blocked by an inverted dT residue (IDT). Templates were annealed to DNA primers of different lengths (see Table S2) by mixing 1 µM of the template with of 400 µM of 2 nt, 200 µM of 5 nt DNA primers or 2 µM of 10, 15 or 20 nt DNA primers in 100 µL of TE (TrisEDTA; 10 mM Tris-HCl pH 7.5, 1 mM EDTA) and heating to 95°C for 3 min followed by cooling to 25°C at 0.1 °C/min in a T100 thermal cycler (Bio-Rad). Primer extension assays were performed either as time courses (up to 240 min) in 80 µL or as single time points (20 min) in 20 µL of reaction medium containing 20 mM NaCl, 10 mM MgCl2, and 20 mM Tris-HCl pH 7.5, with or without 1 mM MnCl2. For the reactions, 250 nM of the annealed template-primer substrate was pre-incubated with 500 nM WT or mutant G2L4 or GII RTs in reaction medium for 30 min at room temperature. Reactions were initiated by adding 1 mM dNTPs (final concentration; an equimolar mix of 1 mM each of dATP, dCTP, dGTP and dTTP) plus 1 µCi [*α*-^32^P]-dTTP (3000 Ci/mmol; Perkin Elmer) and incubating the reaction at 37°C for times indicated in Figure Legends for individual experiments. Aliquots (10 µL) were removed at each time point and quenched by adding 2 µL of 6X stop solution (25 mM EDTA (Sigma-Aldrich), 0.5 U/µL proteinase K (New England Biolabs)) and incubating for 15 min at 37°C. The samples were then mixed with an equal volume of 2X loading dye (95% formamide, 0.25% SDS, 25 mM EDTA and 0.1% xylene cyanole and bromophenol blue), and analyzed by electrophoresis in a denaturing 20% polyacrylamide gel, with 5’-labeled DNA primer and template oligonucleotides as size markers. The gel was scanned with a phosphorimager (Typhoon FLA 9500; GE Healthcare). The fractions of the labeled dTTP incorporated into the product were determined by quantitating bands using ImageQuant TL 8.1. This fraction was multiplied by the dTTP concentration and divided by the number of Ts per extension product to determine the concentration of extended products, which was then plotted as a fraction relative to the template concentration. Time course data were fit by a first-order rate equation using Prism 9.0. For reactions that were slow and progressed approximately linearly during the observation time, such that the reaction endpoint could not be determined from the data, the fitting was done with the endpoint (reaction amplitude) forced to the same value as for corresponding reactions that had defined endpoints. Amplitude values that were forced during fitting are indicated by parentheses in the tables above the corresponding plots.

### Terminal transferase assays

The substrates for terminal transferase assays were the same 50-nt DNA or RNA oligonucleotides used in primer extension assays (Table S2) but without 3’ blocking groups. Terminal transferase assay were done either as time courses (up to 60 min) in 80 µL or as single time points (20 min) in 20-µL of reaction medium. The RT (500 nM) was preincubated with 5’-labeled oligonucleotide substrate (10 nM) in reaction medium containing 20 mM NaCl, 10 mM MgCl2, and 20 mM TrisHCl pH 7.5 with or without 1 mM MnCl2 for 30 min at room temperature and initiated by adding 1 mM dNTPs (final concentration). The reactions were incubated at 37°C for times indicated for individual experiments, and quenched as described above for primer extension assay. The products were analyzed in a denaturing 6% polyacrylamide gel with a 5’-labeled RiboRuler Low Range RNA Ladder (Thermo Scientific) as size markers, and the gel was dried and scanned with a phosphorimager (Typhoon FLA 9500; GE Healthcare). The products were quantified by ImageQuant TL 8.1, and the data were analyzed as described above for primer extension assays.

### Snap-back replication assays

The substrates for snap-back replication assays were 50-nt DNA or RNA oligonucleotides (see Table S2) without 3’ blockers. Snap-back replication assays were done as time courses by preincubating 10 nM 5’-labeled 50-nt DNA or RNA oligonucleotides with 500 nM enzyme in 80 μl of reaction medium (see above) for 30 min at room temperature. Reactions were initiated by adding 1 mM dNTPs (equimolar mix of 1 mM each of dATP, dCTP, dGTP and dTTP) and 10 µL portions were taken at time points (up to 240 min). The reactions were quenched by adding 2 µL of 6X stop solution (see above for primer extension assays) and analyzed by electrophoresis in a non-denaturing 12% polyacrylamide gel with 5’-labeled Low Molecular Weight DNA Ladder (New England Biolabs) run in a parallel lane. After electrophoresis, the gel was dried and scanned with a phosphorimager (Typhoon FLA 9500; GE Healthcare) The products were quantified by ImageQuant TL 8.1, and data were analyzed as described above for primer extension assays.

For high-throughput sequencing of snap-back replication products, the reactions were scaled up to use 1 µM of the 50-nt DNA oligonucleotide substrate and 1 µM RT together with 1 mM dNTPs in 100 μl of reaction medium (see above) and incubated at 37°C for 3 h. The reaction was terminated by adding proteinase K and EDTA, and the products were cleaned-up with an Oligo Clean & Concentrator Kits (Zymo Research). Nucleic acids concentrations were determined with a Nanodrop One (Thermo Scientific), and 50 ng of product was used to prepare TGIRT-seq libraries for high-throughput sequencing.

High-throughput sequencing libraries were constructed by using a variation of the TGIRT-seq method (Xu et al., 2019, 2021). First strand DNA synthesis was initiated at the 3’ end of the snapback DNA product by template switching from an RNA/DNA heteroduplex consisting of a 34-nt RNA containing an Illumina R2 adapter sequence annealed to a complementary 35-nt DNA leaving a single-nucleotide 3’ overhang (an equimolar mix of A, C, G, and T) that can base pair to the 3’ end of the DNA product, resulting in a full-length DNA copy of the product with an the reverse complement of the Illumina R2 adapter (denoted R2R) seamlessly linked to its 3’ end. After clean up using a MinElute PCR Purification Kit (Qiagen), the second-strand DNA synthesis was done by annealing 200 nM of a snap-back-specific R1R DNA oligonucleotide whose 3’ end was complementary to 3’ end of the DNA product, followed by a single-cycle of PCR using Phusion High-Fidelity PCR Master Mix with HF Buffer (New England Biolabs) (98°C for 10 sec pre-denaturation followed by 98°C for 5 sec, 60°C for 10 sec, and 72°C for 15 sec). After two rounds of clean up with 1.4 X AMPure XP beads (Beckman Coulter) with elution in 25 µL of double-distilled H2O, the products were amplified by PCR using Phusion High-Fidelity PCR Master Mix with HF Buffer (New England Biolabs) with 200 nM of Illumina multiplex primer and 200 nM of Illumina index barcode primers (98°C for 10 sec pre-denaturation followed by 12 cycles of 98°C for 5 sec, 60°C 10 sec, 72°C 15 sec). The resulting TGIRT-seq libraries were cleaned up by using 1.4 X AMPure XP beads (Beckman Coulter) and eluted in 25 µL double-distilled H2O, with 1 µL analyzed on an Agilent 2100 Bioanalyzer using a High Sensitivity DNA chip (Agilent) to assess product profiles and concentrations. The remainder of the library was sequenced on an Illumina MiSeq V2 instrument to obtain ∼1 million 2 x 75 nt paired end reads per sample at the Genome Sequencing and Analysis Facility (GSAF) at the University of Texas at Austin.

To analyze the product sequences, Illumina TruSeq adapters and PCR primer sequences were trimmed from the reads with Cutadapt v3.2 (sequencing quality score cut-off at 20; p-value <0.01), and reads <15-nt after trimming were discarded (Martin, 2011). After merging the trimmed pair-ended reads by using BBMerge (Bushnell et al., 2017), the template sequence (5’-GCAATAATCTATACAATACAACACATACAAACAAATTCTTAAGGTCCCAA-3’) was trimmed from the 5’ ends of the merged reads by using Cutadapt, and downstream sequences were sorted to collect unique sequences. These unique sequences were then aligned to the reverse complement of the template sequence (5’-TTGGGACCTTAAGAATTTGTTTGTATGTGTTGTATTGTATAGATTATTGC-3’) using CLUSTALW (https://www.genome.jp/tools-bin/clustalw) with default settings and manually adjusted to correct minor misalignments introduced by CLUSTALW.

### Microhomology-mediated end-joining assays

Initial MMEJ assays were by done using a double-stranded DNA with a 15-nt single-stranded 3’ overhang with or without a 3’ terminal self-complementary microhomology sequence mimicking 5’-strand resected double-stranded DNAs on either side of a double-strand break. A 5’-labeled 53nt oligonucleotide (D1) ending with a 4-nt microhomology or a control lacking the microhomology was annealed to an unlabeled complementary 39-nt DNA oligonucleotide with a 3’ blocker (D2) at a ratio of 1:2 in 100 µL of TE by heating to 95°C for 3 min followed by cooling to 25°C at 0.1 °C/min in a T100 thermal cycler (Bio-Rad). In Figures, the left- and right-hand are denoted D1/D2, and D1’/D2’, respectively. For MMEJ assays, 10 nM of the annealed MMEJ substrate was preincubated with 500 nM enzyme in 80 μL of reaction medium for 30 min at room temperature. The reaction was initiated by adding 1 mM dNTPs, incubated at 37°C for times indicated for individual experiments (up to 240 min), and quenched as described above for the primer extension assays. The products were analyzed by electrophoresis in a non-denaturing 12% polyacrylamide gel against a 5’-labeled Low Molecular Weight DNA Ladder (New England Biolabs) and quantitated with a phosphorimager (Typhoon FLA 9500; GE Healthcare). Some experiments used substrates that varied the length and sequence of the 3’ microhomology (4-bp CCGG or TTAA; 10-bp CCCCCGGGGG; non-complementary CCAA) or changed the length of the single-stranded gap flanking the microhomology by varying the length of the D2 oligonucleotide (27, 33 and 39 nt). The products were quantified by ImageQuant TL 8.1, and data were analyzed as described above for primer extension assays.

For high-throughput sequencing of MMEJ products, the reactions were scaled up to use 1 µM of unlabeled annealed dsDNAs and 1 µM of enzyme in 50 µL of reaction medium. The samples were incubated at 37°C for 4 h, and terminated by adding 10 µL of 6X stop solution and incubating for 15 min at 37°C. The products were cleaned up using 1.8 X AMPure XP Beads and eluted with 25 µL of double-distilled H2O, digested with BsrGI (New England Biolabs), and ligated into the BsrGI site of BsrGI-linearized pKS-SacB. After transforming the ligated plasmids into *E. coli* HMS174 (DE3), cells were incubated overnight in 5 mL of LB containing 50 µg/mL carbenicillin and 6% sucrose, to select cells containing plasmids in which the *sacB* gene was inactivated by an insertion into the BsrGI site. The plasmids were isolated by using a Monarch Plasmid Miniprep kit (New England Biolabs) according to the manufacturer’s protocol and PCR amplified using primers MMEJ R1 and MMEJ R2R, which are complementary to sequences that flank the *sacB* BsrGI cleavage site and added Illumina R1 and R2R sequences to either end of the PCR product. The PCR was done with 1-2 ng of plasmid and 200 nM of each primer in Phusion High-Fidelity PCR Master Mix (New England Biolabs) with pre-denaturation at 98°C for 5 sec followed by 12 cycles of 98°C for 5 sec, 65°C for 10 sec, and 72°C for 15 sec. After PCR amplification, the products were cleaned up with 0.4 x AMPure XP Beads to remove the plasmid, followed by 1.4 X AMPure XP Beads clean-up to remove the primers. The PCR products were eluted in 25 µL double-distilled H2O, and 1 µL was analyzed on an Agilent 2100 Bioanalyzer with a High Sensitivity DNA chip to confirm the product and determine product concentration. For sequencing, Illumina multiplex and bar code primers were added by PCR (1 µL of MMEJ product and 200 nM primers in Phusion High-Fidelity PCR Master Mix (New England Biolabs) with 98°C, 5 sec pre-denaturation followed by 12 cycles of 98°C for 5 sec, 60°C for 10 sec, and 72°C for 15 sec. After 1.4 X AMPure XP Beads clean-up to remove primer dimers, 1-µL of the product was analyzing on an Agilent 2100 Bioanalyzer with High Sensitivity DNA chip to assess the product profile and concentration, and the libraries were sequenced on an Illumina MiSeq v2 instrument to obtain ∼1 million 2 x 150 nt paired end read of each sample at the University of Texas MD Anderson Cancer Center, Science Park.

For the analysis of product sequences, Illumina TruSeq adapters and PCR primer sequences were trimmed from the reads with Cutadapt v3.2 (sequencing quality score cut-off at 20; p-value <0.01), and reads <15 nt after trimming were discarded (Martin, 2011). Trimmed pair-ended reads were then merged by using BBMerge (Bushnell et al., 2017). Sequences between two BsrGI sites that were longer than 45 nt were analyzed by using a customized R script to categorize the type of the MMEJ products. Merged reads of 82-86 nt that contain the template sequence on one end and its reverse complement on the other end were classified as terminal MMEJ products. Merged reads shorter than 82 nt or longer than 86 nt that contain template sequence only on one end were categorized as internal, discontinuous or terminal transfer-mediated MMEJ products.

### CRISPR-Cas9/RT induced *thyA* DSBR assay

The ability of WT and mutant G2L4 and GII RTs to repair double-strand breaks in *E. coli* HMS174 (DE3) was assessed by using a using a CRISPR-Cas9 introduce a DSB into the *E. coli thyA* gene. CRISPR-Cas9 components and G2L4 and GII RTs were expressed in *E. coli* HMS174 (DE3) using a two-plasmid system based on that described by Chen et al., 2018. First, pCas9+RT Tet^R^-based plasmids (Figure 7A), which express *Streptococcus pyogenes* Cas9 and WT or mutant G2L4 and GII RTs using two independent L-arabinose-inducible promoters, were transformed into *E. coli* HMS174 (DE3) chemically competent cells via heat shock, and the transformed cells were incubated overnight in 5 mL of LB containing tetracycline (25 µg/mL) in a 50-mL conical tube (Sarstedt) at 37°C with shaking (250 rpm) for 14-16 h. Then, 1 mL of the culture was transferred into 100 mL LB containing tetracycline (25 µg/mL) and incubated at 37°C until O.D.600=1.0. Expression of the RT and Cas9 proteins was induced by adding L-arabinose (2 mg/mL) and incubating at 18°C with shaking (100 rpm) for 19-21 h. To introduce the second plasmid pACRISPR *thyA* sgRNA expressing *thyA* guide RNAs targeted to a site in the *E. coli thyA* gene, 25 mL of the overnight culture was centrifuged at 4000 x g for 15 min at 4°C, and the cell pellet was gently resuspended in 20 mL of ice-cold 10% glycerol, and then centrifuged and resuspended in 10% glycerol twice more and the final pellet resuspended in 500 µL of ice-cold 10% glycerol. 50-µL portions of the cells were then electroporated with 1 µg pACRISPR vector or pACRISPR*thyA* sgRNA plasmids in a 2-mm cuvette at 3.0 kv, 200 Ω, 25 µF using a Gene Pulser Xcell Electroporation System (Bio-Rad). The bacteria were recovered in 1-mL fresh SOC medium supplemented with 2 mg/mL L-arabinose, 0.5 mM MnCl2, 25 µg/mL tetracycline, 100 µg/mL thymine and incubated at 37°C with shaking (250 rpm) for 1 h. To determine survival and mutation frequencies, the cells were serially diluted in SOC medium and plated on 2X yeast tryptone (YT) plates supplemented with 2 mg/mL L-arabinose, 10 mM MgCl2, 0.5 mM MnCl2, 25 µg/mL tetracycline, 50 µg/mL carbenicillin and 100 µg/mL thymine with or without trimethoprim (200 µg/mL), the latter to select against cells containing a functional *thyA* gene. Colonies were counted after 16-48 h at 37°C. p-values were calculated by student’s unpaired t-test in Prism 9.0.

To sequence *E. coli* chromosomal *thyA* genes with repaired DSBs, a 100-µL portion of the culture after electroporation of the *thyA* sgRNA plasmid was inoculated into 5 mL 2X YT medium containing 2 mg/mL L-arabinose, 10 mM MgCl2, 0.5 mM MnCl2, 25 µg/mL tetracycline, 50µg/mL carbenicillin and 100 µg/mL thymine plus 200 µg/mL trimethoprim to selected for *thyA* mutants, and incubated in a 50-mL conical tube (Sarstedt) at 250 rpm and 37°C for 60-72 h. The cells were then centrifuged at 4000 x g for 10 min, and genomic DNA was extracted by using a Monarch Genomic DNA Purification Kit (New England Biolabs). The *thyA* gene region was amplified from the genomic DNA (5 ng) with Phusion High-Fidelity PCR Master Mix (New England Biolabs) using 200 nM of *thyA* forward and reverse primers that give amplicons of 750 bp, 2.5 kb, or 5 kb centered on the *thyA* gene DSB site. PCR conditions were 98°C for 10 sec pre-denaturation followed by 25 cycles of 98°C for 10 sec, 60°C for 30 sec, and 72°C for 30 sec extension for the 750-bp amplicon or 72 °C for 1 min for extension for the 2.5 or 5 kb amplicons. The PCR products were analyzed by electrophoresis at 120 V in agarose gels containing Tris-acetate-EDTA buffer (2% agarose for the 750 bp amplicons and 1% agarose for the 2and 2.5-kb amplicons), and the gel regions that contain *thyA* genes with deletions were extracted by using a Monarch DNA gel extract kit (New England Biolabs). 1 ng of extracted PCR products were analyzed by an Agilent High Sensitivity DNA kit on an Agilent 2100 Bioanalyzer (Agilent), and the remainder was cloned by using PCR Cloning Kit (New England Biolabs) and analyzed by Sanger sequencing.

### Data access

Datasets for *P. aeruginosa* whole genome sequencing, TGIRT-seq, and sequencing of Snap-Back DNA synthesis and MMEJ products in biochemical experiments have been deposited in the Sequence Read Archive (SRA) under accession numbers PRJNA814398. A gene counts table, dataset metadata file, and scripts used for data processing and plotting have been deposited in GitHub: https://github.com/reykeryao/Seung.

---

**Figure S1.**
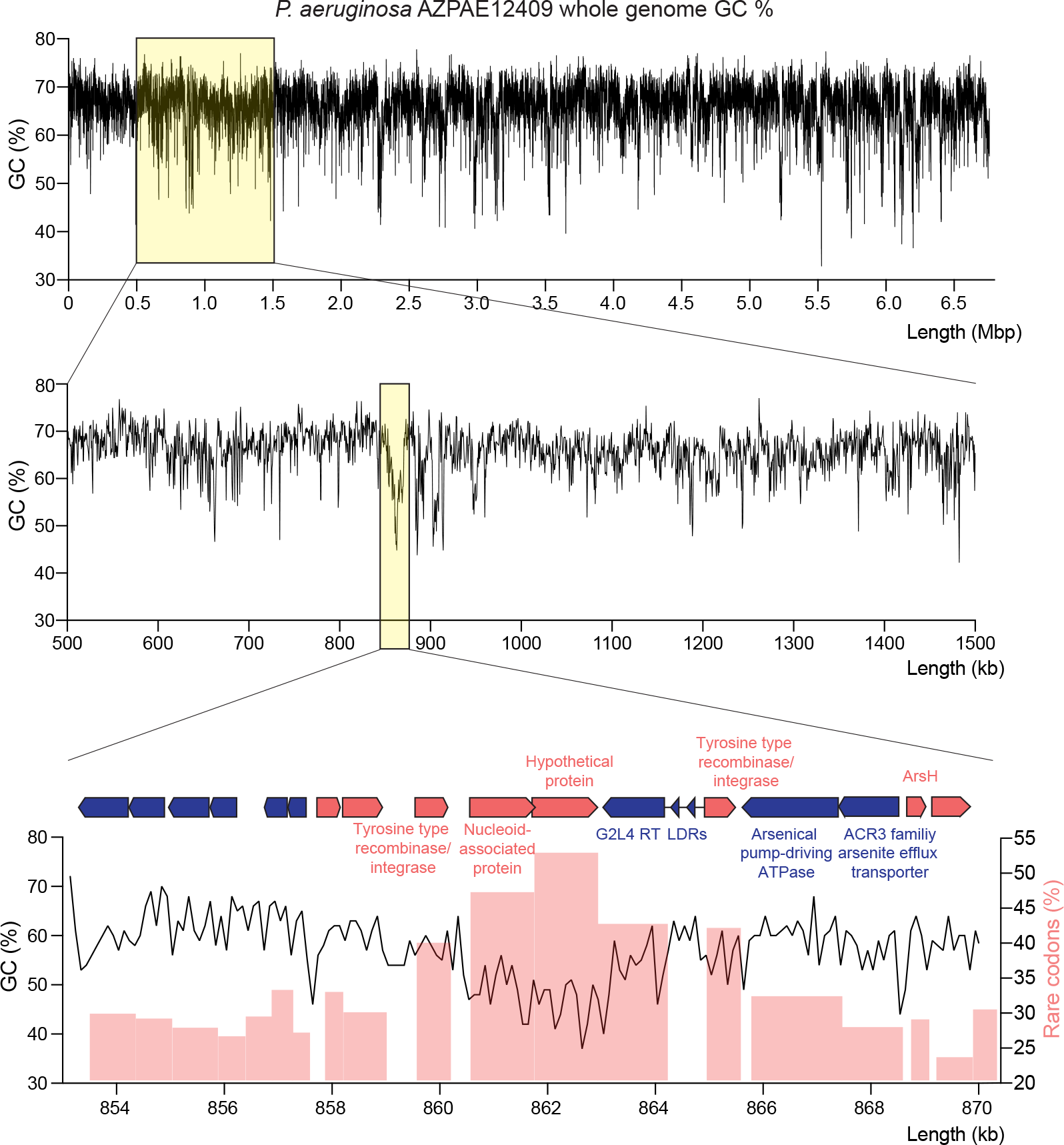
Characteristics of the genomic region surrounding the G2L4 RT ORF in *P. aeru- ginosa* AZPAE12409 The top panel shows the GC content of the *P. aeruginosa* AZPAE12409 genome with the yellow box indicating a 1-Mb region containing the G2L4 RT gene. The middle panel shows the GC content of this region with a second yellow box indicating an ∼20-kb region centered on the G2L4 RT gene. The 6.1-kb region encoding the G2L4 RT has a relatively low GC content compared to other regions of the genome. The bottom panels show plots of GC content (black line) and rare codon usage (red bars) for the region shown in the map above. Rare codons were defined as codons that are used at frequencies ≤1% of the average codon use in the *P. aeruginosa* PAO1 reference strain: UUU, UUA, UUG, CUU, CUA, AUU, GUU, GUA, UCU, UCA, CCU, CCA, ACU, ACA, ACG, GCU, GCA, UAU, CAU, CAA, AAU, AAA, UGU, UGC, CGU, CGA, ACU, AGA, AGG, GGU, GGA, GGG); LDRs, 140-bp direct repeats upstream of the G2L4 RT ORF.

**Figure S2.**
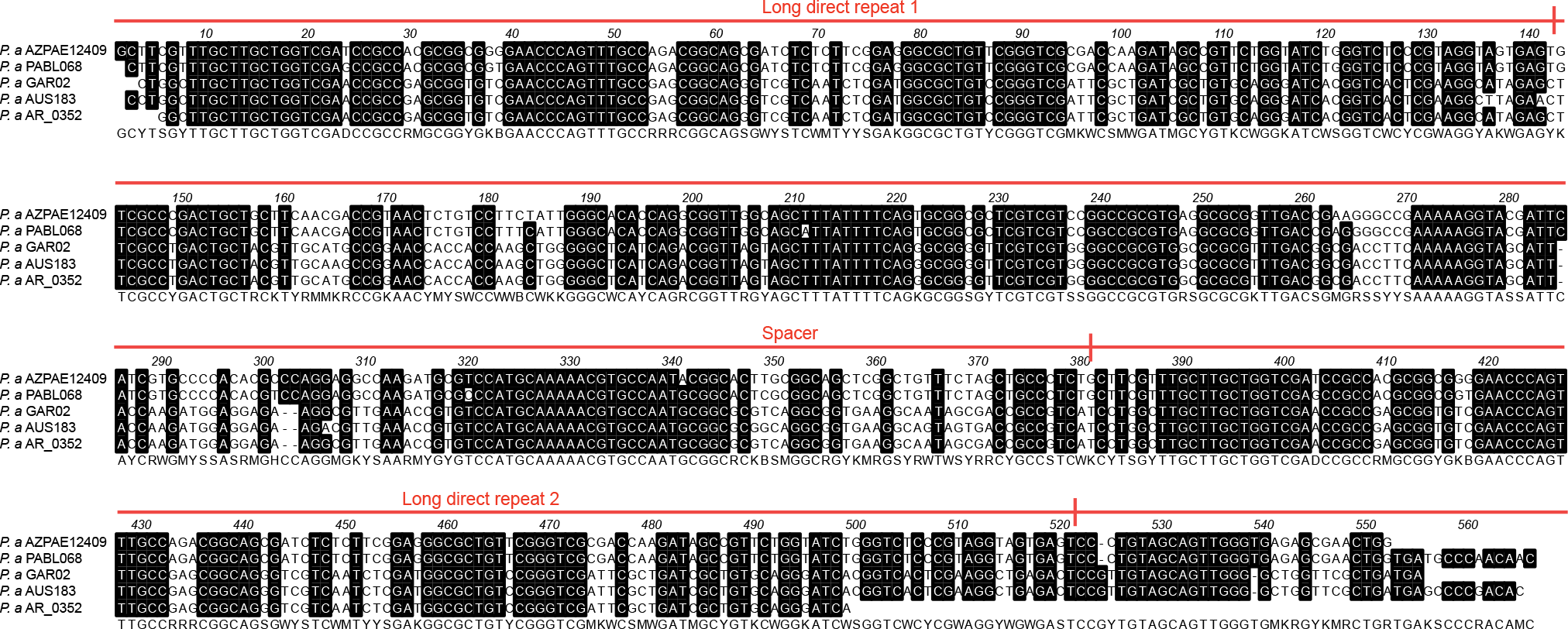
Nucleotide sequence alignment of the 140-bp long direct repeats and 240-bp spacer in different *P. aeruginosa* strains ClustalW alignment of the region upstream of the G2L4 RT containing the two long direct repeats and spacer from five *P. aeruginosa* strains. The position of the repeats and the spacer are indicated above the alignment. Identical nucleotides (>80%) are shown as white letters on black background. A consensus sequence is shown at the bottom. The sources for the aligned sequences were: *P. aeruginosa* AZPAE12409 whole genome sequencing data GCA_000797005.1; *P. aeruginosa* PABL068, whole genome sequencing data GCA_003411275.2; *P. aeruginosa* GAR02, Genebank accession NZ_JABUGS010000001; *P. aeruginosa* AUS183, Genebank accession NZ_NSZP01000001.1.; *P. aeruginosa* AR_0352, Genebank accession QMGJ01000001.1.

**Figure S3.**
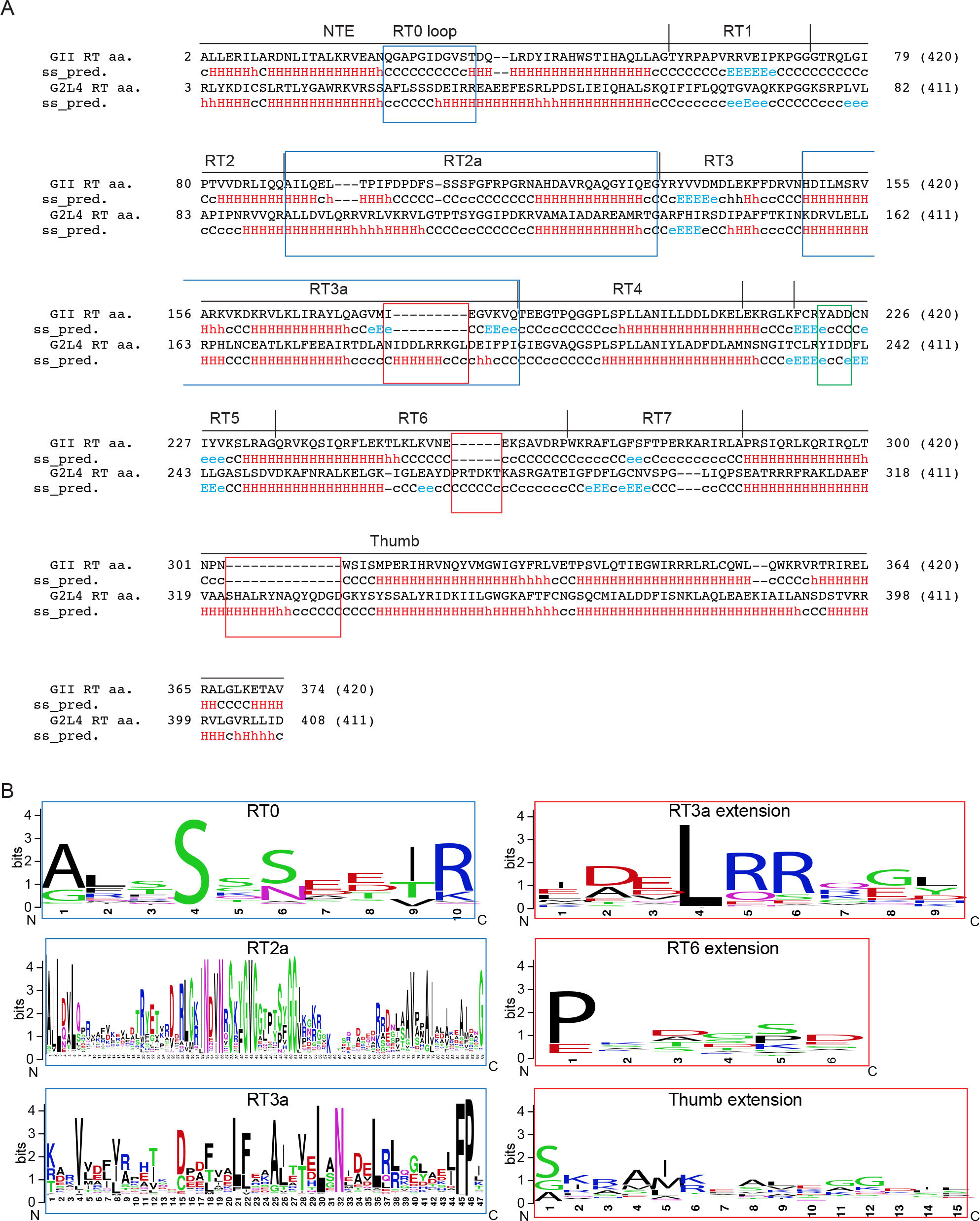
Predicted secondary structure and amino acid sequence alignment of G2L4 RT and GII RT (A) Sequence alignment and predicted secondary structures of the G2L4 and GII RTs. Secondary structure was predicted by using HHPred (Zimmermann et al., 2018). Conserved sequence motifs found in all RTs (RT1 to 7) and the thumb domain are delineated on top. The RT0 loop and the RT2a and RT3a insertions are in blue boxes. The YxDD motif at the RT active site motif is boxed in green. Red boxes indicate sequence additions in G2L4 RT relative to the GII RT. *α*-helices, H/h (red); *β*-sheets, E/e (blue); Coiled coil, C/c (black). Upper- and lower-case letters indicate higher and lower confidence predictions, respectively (Gabler et al., 2020). (B) Weblogos. ∼250 unique G2L4 RTs were aligned with ClustalW and the alignment was manually optimized. Sequences corresponding to those in the boxes shown in panel (A) were used to generate sequence logos with default parameters (Crooks et al., 2004). Sequence logos for the RT0 loop, RT2a and RT3a in G2L4 RTs are shown on the left, and sequence logos for regions of the G2L4 RT that are not present in GII RT are shown on the right.

**Figure S4.**
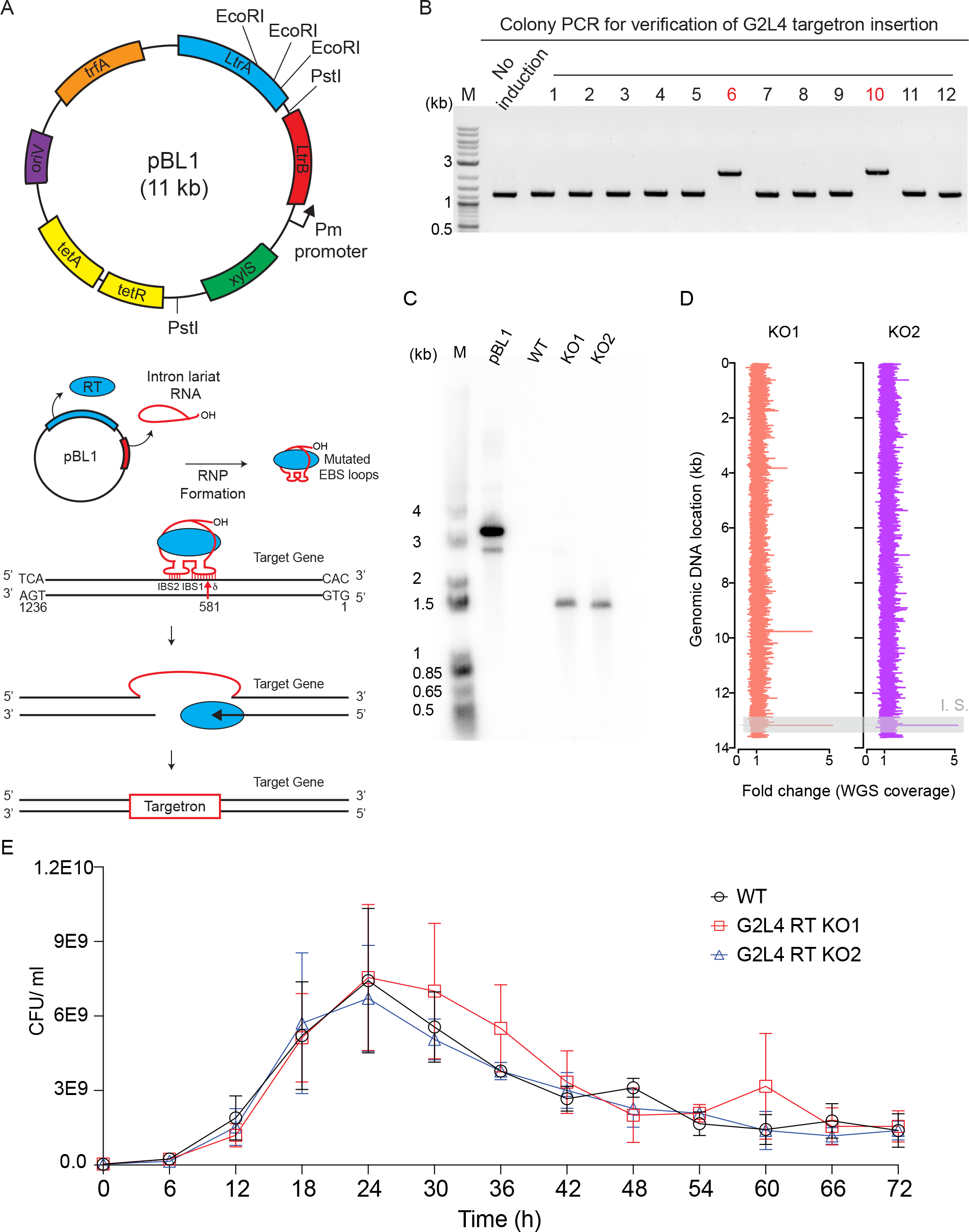
Disruption of the G2L4 RT ORF by targetron insertion (A) Schematic of the targetron gene knock-out method. A map of the pBL1 plasmid used for tar- getron expression is shown at the top. Abbreviations: Pm promoter, m-toluic acid-inducible pro- moter; LtrA, Ll.LtrB group II intron RT; LtrB, Ll.LtrB-*Δ*ORF group II intron; *oriV*, replication origin; *tetA*, tetracycline-resistance gene; *tetR*, tetracycline repressor; *trfA*, initiation of plasmid replication; xylS, cognate regulator for the Pm promoter. After transforming the targetron donor plasmid into *P. aeruginosa* AZPAE12409, targetron expression was induced overnight with 2 mM m-toluic acid at 30°C. The LtrA protein and the spliced intron lariat RNA form an RNP complex that targets the G2L4 RT ORF via base-pairing between the intron RNA’s EBS and *δ* sequences and the antisense strand (nucleotides 578-592) of the chromosomal G2L4 RT gene followed by insertion of the intron within the G2L4 RT gene (nucleotide 581; red arrow). (B) Disruptants in which the targetron inserted in the G2L4 RT ORF were identified by colony PCR, using primers flanking the intron-insertion site (Yao and Lambowitz, 2007), with colonies 6 and 10 giving larger PCR products reflecting the targetron insertion. (C) Southern hybridization showing a single tar- getron insertion in the G2L4 RT ORF. Genomic DNA of *P. aeruginosa* AZPAE12409 WT and G2L4 RT KO strains was digested with PstI and EcoRI, which is predicted to generate a 1.5-kb band containing the integrated targetron. The pBL1 vector digested with the same enzymes pro- duced a 3.6-kb DNA fragment. The digested DNAs were run in a 1% agarose gel against a 5’- labeled 1-kb Plus DNA Ladder (Invitrogen), and Southern hybridization using a 5’-labeled tar- getron probe was done as described in Methods. (D) Fold change of coverage between WT and knockout (KO) strains. The fold change was calculated as the ratio of coverage in the WT and KO in 500-bp bins arranged by the order of contig numbers. The line highlighted in gray corresponds to contig 110, which contains the targetron insertion site (I.S.) in the KO strains. (E) Growth curve of *P. aeruginosa* AZPAE12409 WT and G2L4 RT KO strains. Single colonies of each strain were inoculated in Luria-Bertani (LB) media and incubated at 37°C overnight. The next day, cultures were diluted 1:100 into fresh LB, and incubated further at 37°C, 200 rpm. Cells were harvested every 6 h up to 72 h, serially diluted, and plated on LB agar plates to determine the number of colony forming units (CFUs)/mL. The error bars indicate the standard deviation for three repeats using separate cultures (see Methods).

**Figure S5.**
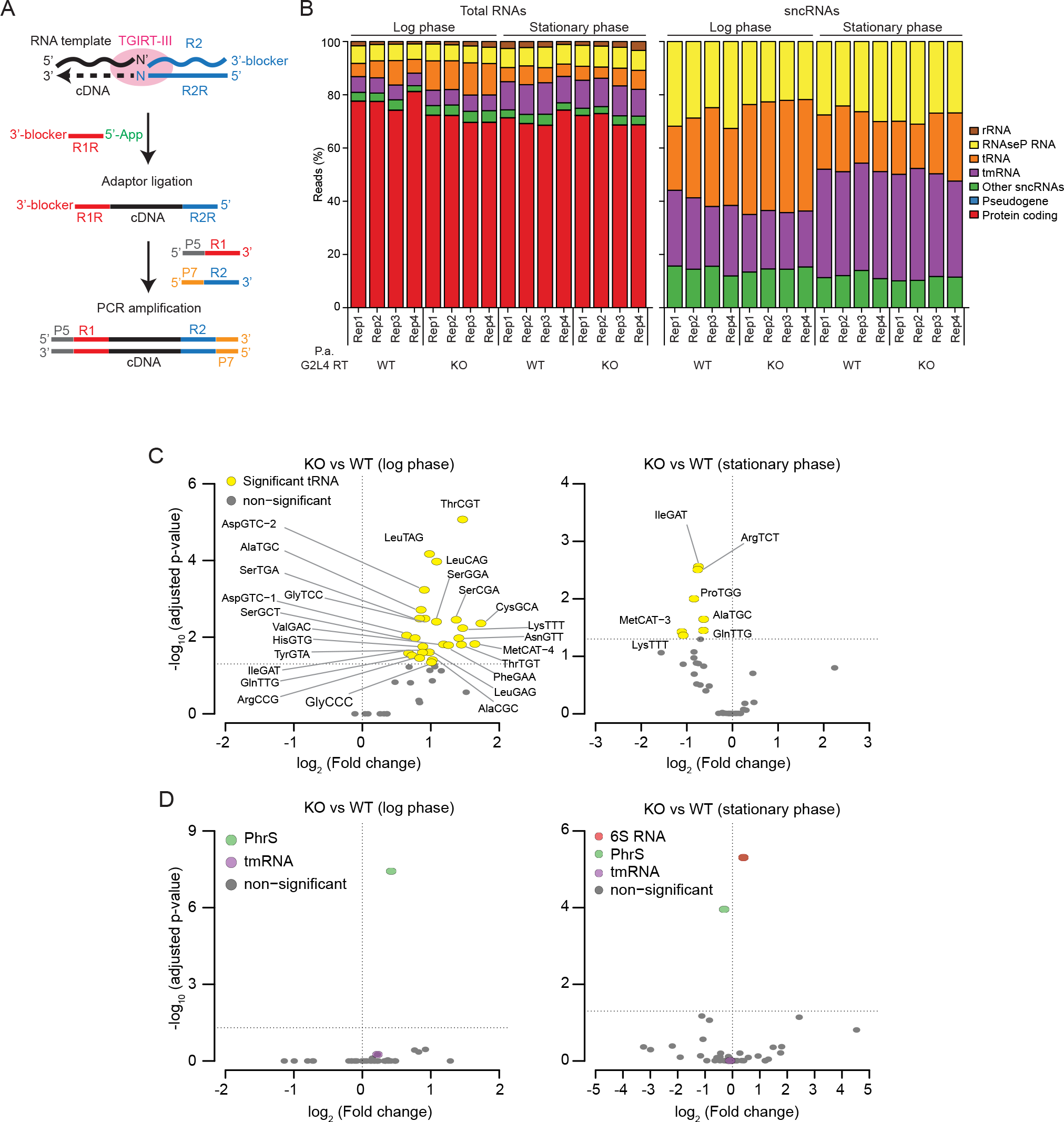
TGIRT-seq of total cellular RNAs from G2L4 WT and KO strains in log and stationary phase(A) Schematic of the TGIRT-seq library preparation method for chemically fragmented, rRNA- depleted bacterial RNAs. First, the TGIRT-III enzyme initiates reverse transcription by template switching to the 3’ end of a target RNA from a synthetic RNA/DNA starter duplex that contains an RNA-seq adapter sequence (the reverse complement of an Illumina Read 2 sequence denoted R2R). The starter duplex has a 1-nt 3’ DNA overhang (an equimolar mix of A, C, G, and T residues, denoted N) that directs template switching by base pairing to the 3’ nucleotide of an RNA template (denoted N’) with concomitant addition of the R2R sequence to the 5’ end of the cDNA. Next, a second RNA-seq adapter (the reverse complement of an Illumina Read 1 sequence denoted R1R) is added to the 3’ end of the completed cDNA by using thermostable 5’ App RNA/DNA ligase, followed by minimal PCR with primers that add Illumina multiplex and index barcode sequences. (B) Stacked bar graph showing the proportions of total reads (left) and sncRNA reads (right) for four replicates each of TGIRT-seq of the P.a. AZPAE12409 WT and G2L4 RT KO strains in log and stationary phases. (C) Volcano plots showing differential expression of tRNAs between G2L4 RT WT and KO strains in log and stationary phases. (D) Volcano plots showing differential ex- pression of sncRNAs between G2L4 RT WT and KO strains in log and stationary phases. Colored ovals in the Volcano plots indicate RNAs that show significantly different expression levels be- tween the KO and wild-type strains (p*≤* 0.05). PhrS, a sncRNA that had a significantly increased expression level in log phase of the G2L4 RT KO strain, stimulates synthesis of a *Pseudomonas* quinolone signal that regulates cellular processes (iron acquisition, cytotoxicity, outer-membrane vesicle biogenesis, and host immune modulatory activities) (Sonnleitner et al., 2011; Lin et al., 2018). Data were normalized by DESeq2.

**Figure S6.**
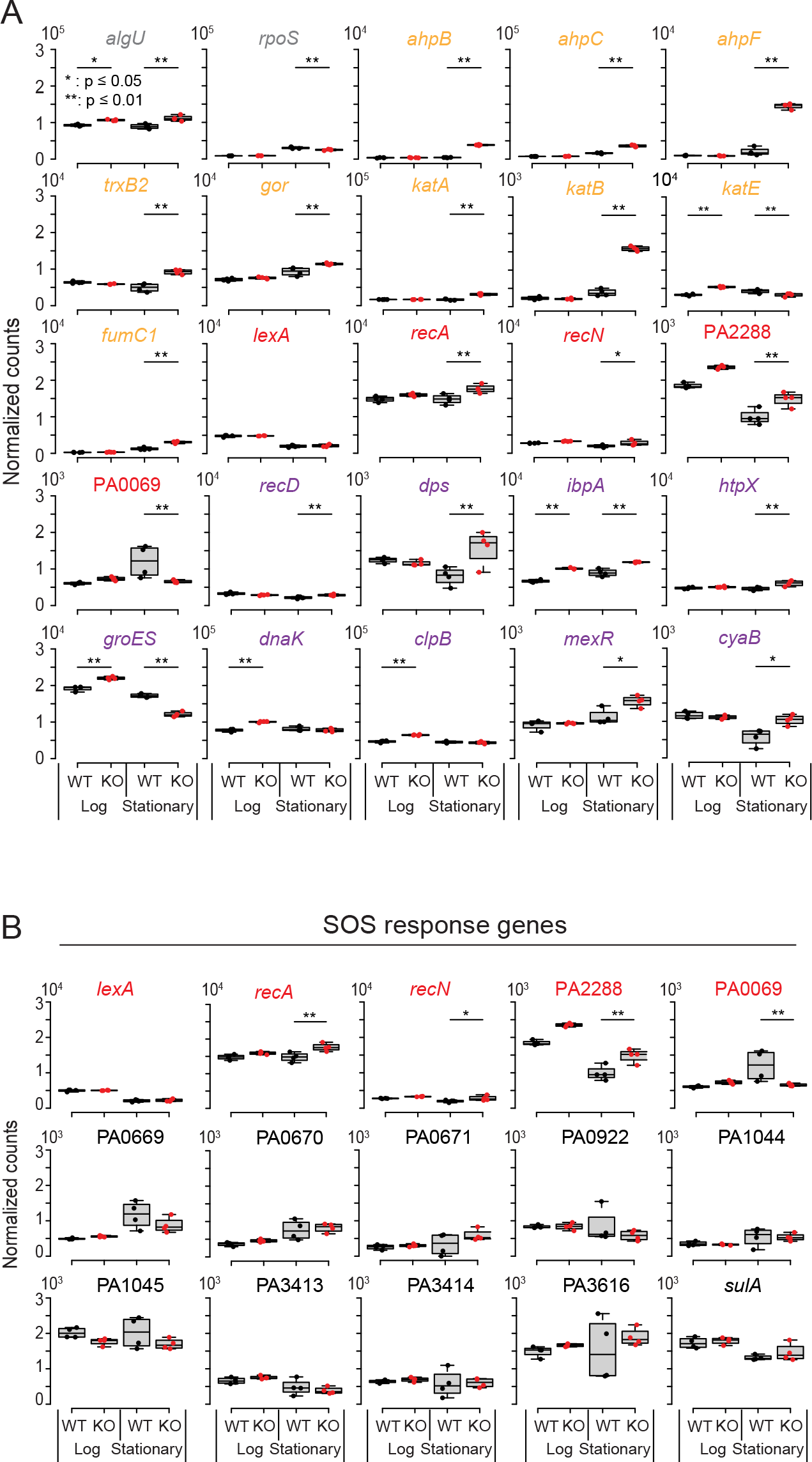
Box plots comparing gene expression levels in the G2L4 RT WT and KO strains during log and stationary phases (A) Box plots of genes showing significantly changed expression levels in the G2L4 RT WT and KO strains during log and stationary phases for four TGIRT-seq repeats. Reads for protein coding genes were normalized with DESeq2. p-values <0.05, *; <0.01, **. (B) Box plots of differential expression of SOS response genes in the G2L4 RT WT and KO strains during log and stationary phases. Gene names are color coded by function: stress response regulation, gray; oxidative stress, orange; SOS response, red; SOS related global stress response, purple.

**Figure S7.**
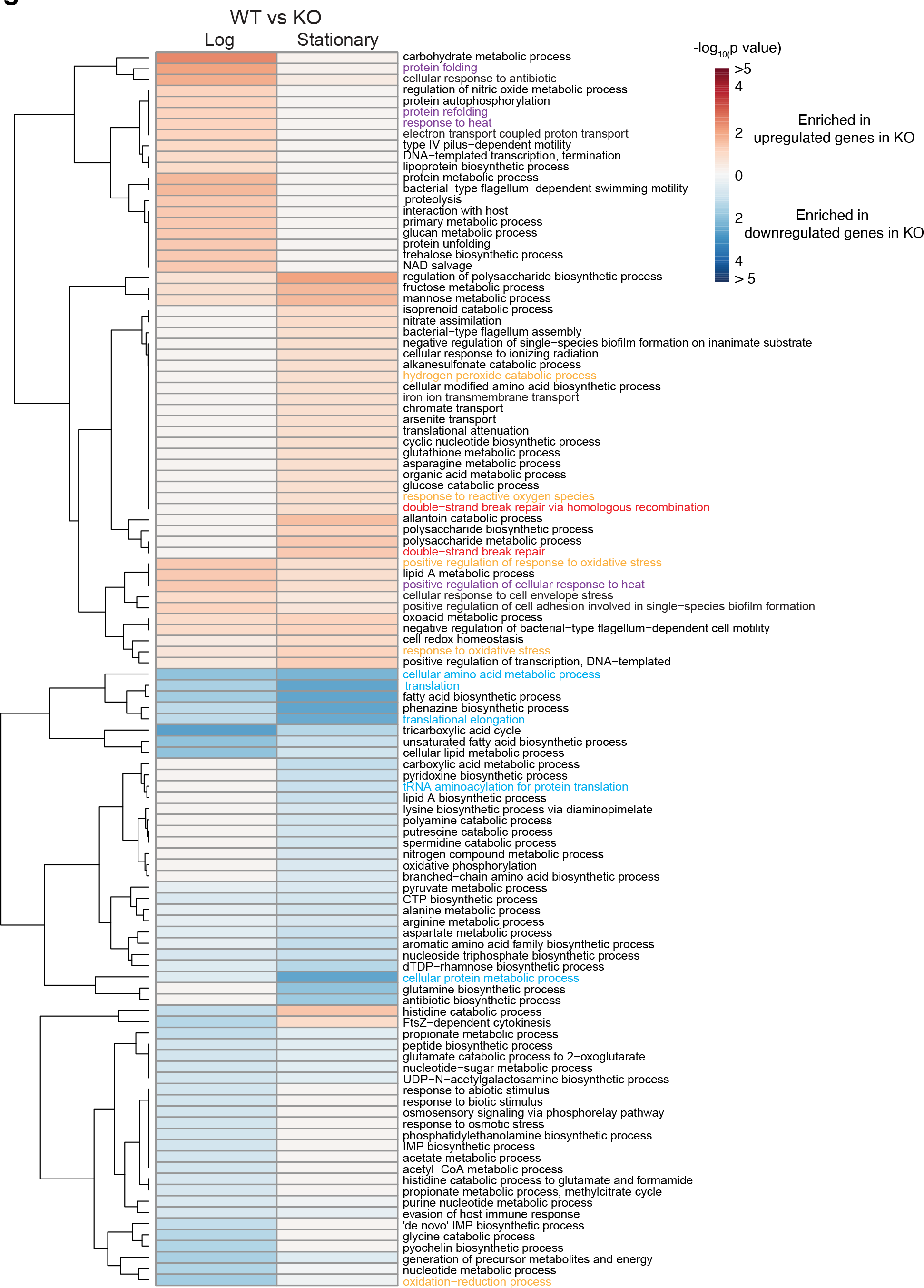
Heat map showing significantly enriched GO terms based on TGIRT-seq of whole cell RNA preparations from the G2L4 RT WT and KO strains in log and stationary phases Heatmap of significantly enriched GO terms in the G2L4 RT KO versus WT strains in log phase (left) and stationary phase (right). The color scale (top right) shows -log10(p-value) of GO term enrichment: up regulated (red; log2FC>0 and adjusted p-value *≤*0.05) and down regulated (blue; log2FC<0 and adjusted p-value *≤*0.05). Only significantly enriched GO terms are included (p-value *≤*0.05). Names of some GO terms are color coded by function: oxidative stress, orange; SOS re-sponse, red; SOS related global stress response, purple; Translation and protein and amino acids metabolic processes, blue.

**Figure S8.**
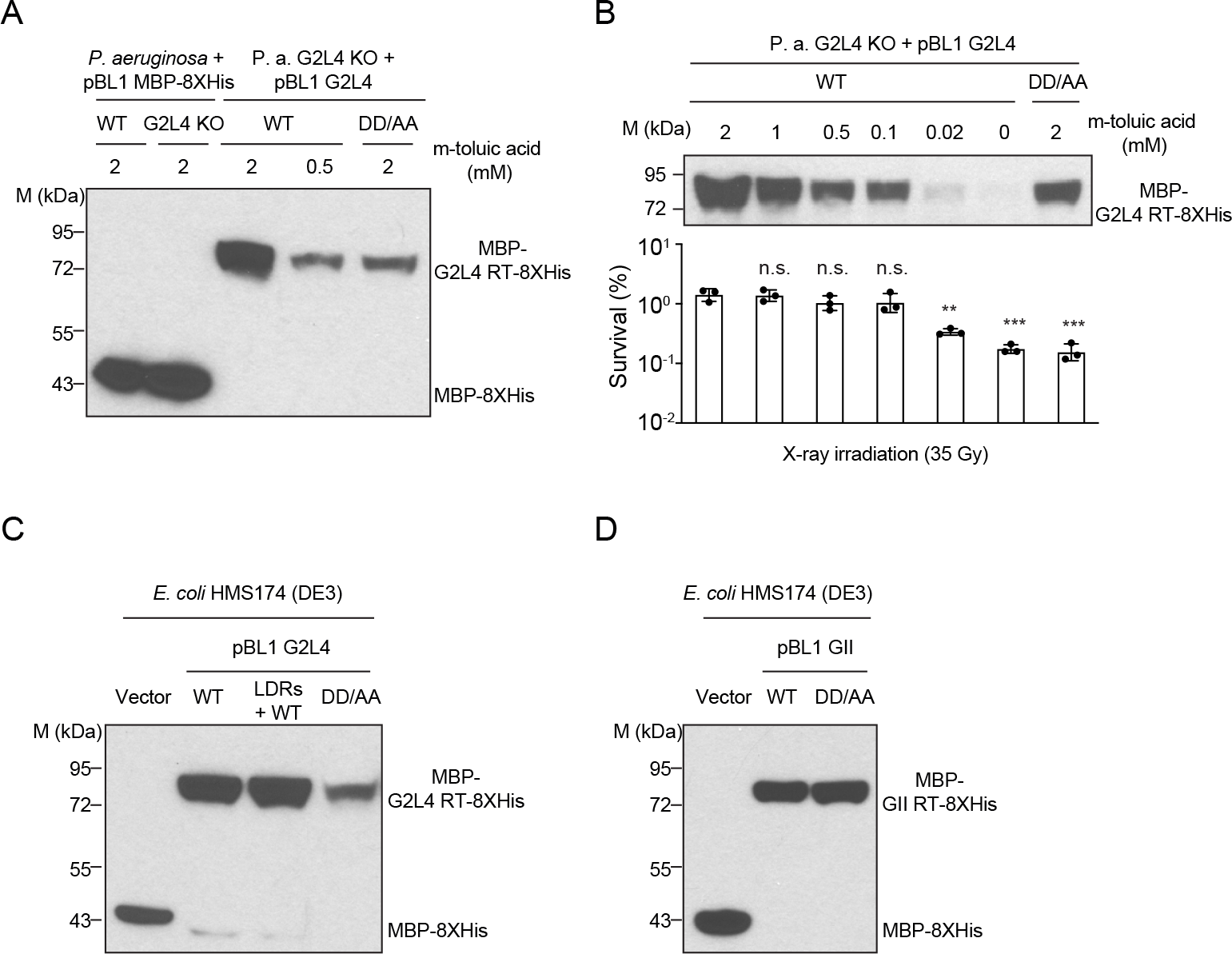
Immunoblots showing expression levels of WT and mutant G2L4 and GII RT in *P. aeruginosa* AZPAE12409 and *E. coli* HMS174 (DE3) (A) Immunoblot of WT and mutant G2L4 RTs expressed in *P. aeruginosa* AZPAE12409 WT and KO in a parallel culture to that used for cell survival assays from pBL1-MBP-G2L4-8XHis versus a vector control expressing MBP with a C-terminal 8XHis tag. Concentrations of m-toluic acid used to induce protein expression are shown above each lane. (B) X-ray irradiation cell survival assays and correlated immunoblots at different expression levels of G2L4 RT induced by different concentrations of m-toluic acid. (C and D) Immunoblots of WT and mutant G2L4 and GII RTs expressed in *E. coli* HMS174 (DE3) in a parallel culture to that used for cell survival assays from pBL1-MBP-GII RT-8XHis after induction with 2 mM m-toluic acid. Immunoblots were probed with an *α*-6XHis-tag primary antibody (Invitrogen) that recognizes the 8XHis Tag (see Methods).

**Figure S9.**
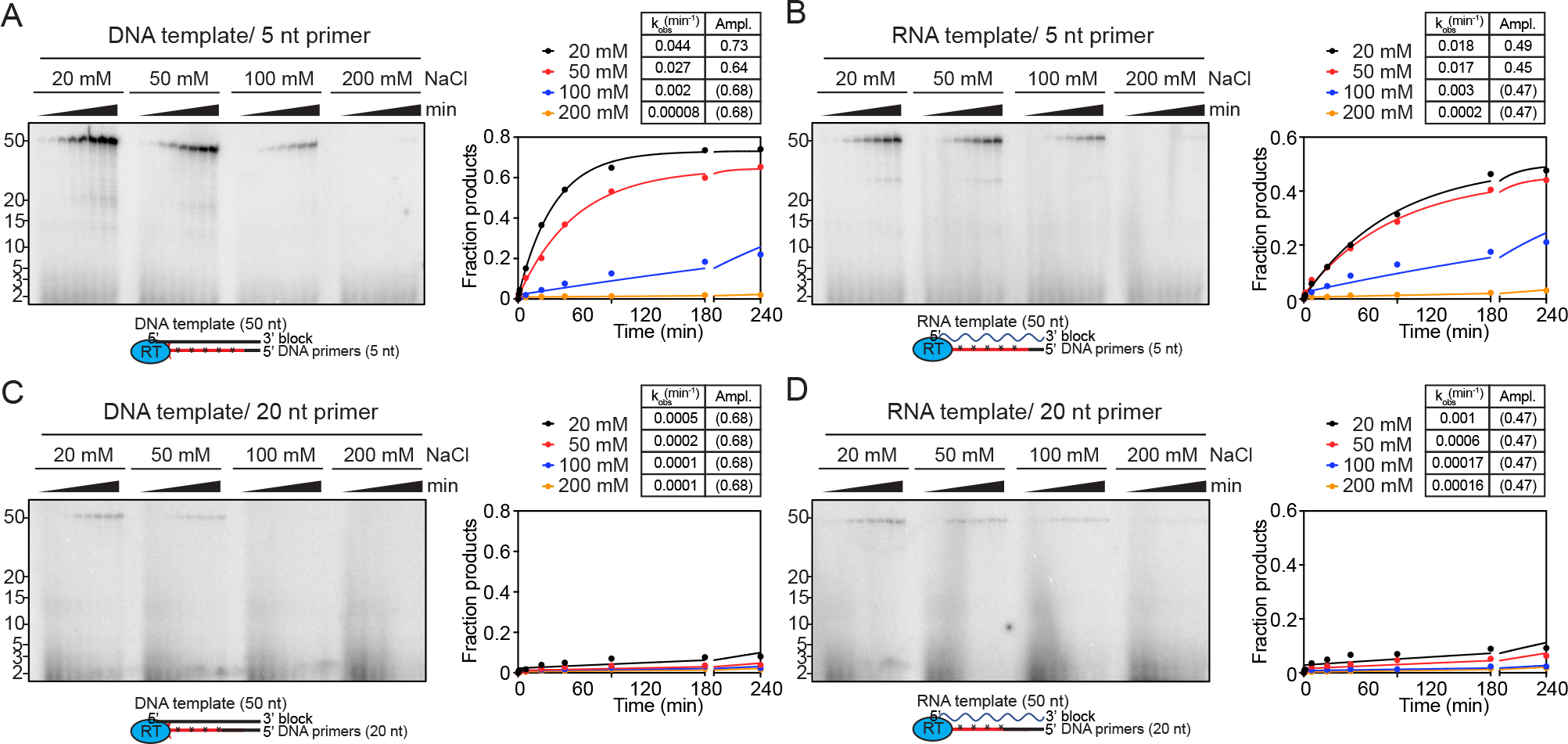
Salt dependence of WT G2L4 RT primer extension activity Primer extension assays were done as described in Methods with 3’-blocked DNA or RNA tem- plates and DNA primers of 5 nt or 20 nt. (A and B) Primer extension activity of WT G2L4 RT with 50-nt DNA or RNA templates and 5-nt DNA primer. (C and D) Primer extension activity of WT G2L4 RT with 50-nt DNA or RNA templates and 20-nt DNA primer. For all panels, the numbers to the left of the gels indicate the positions of size markers in a parallel lane. Tables above the plots show the rate constants (*k*obs) and amplitudes (Ampl.) for production of the labeled 50-nt DNA product obtained by fitting the data to a first-order rate equation. Values of Ampl. in paren- theses indicate that the amplitude was fixed at the given value because the reaction did not reach an endpoint during the measurement time (see Methods).

**Figure S10.**
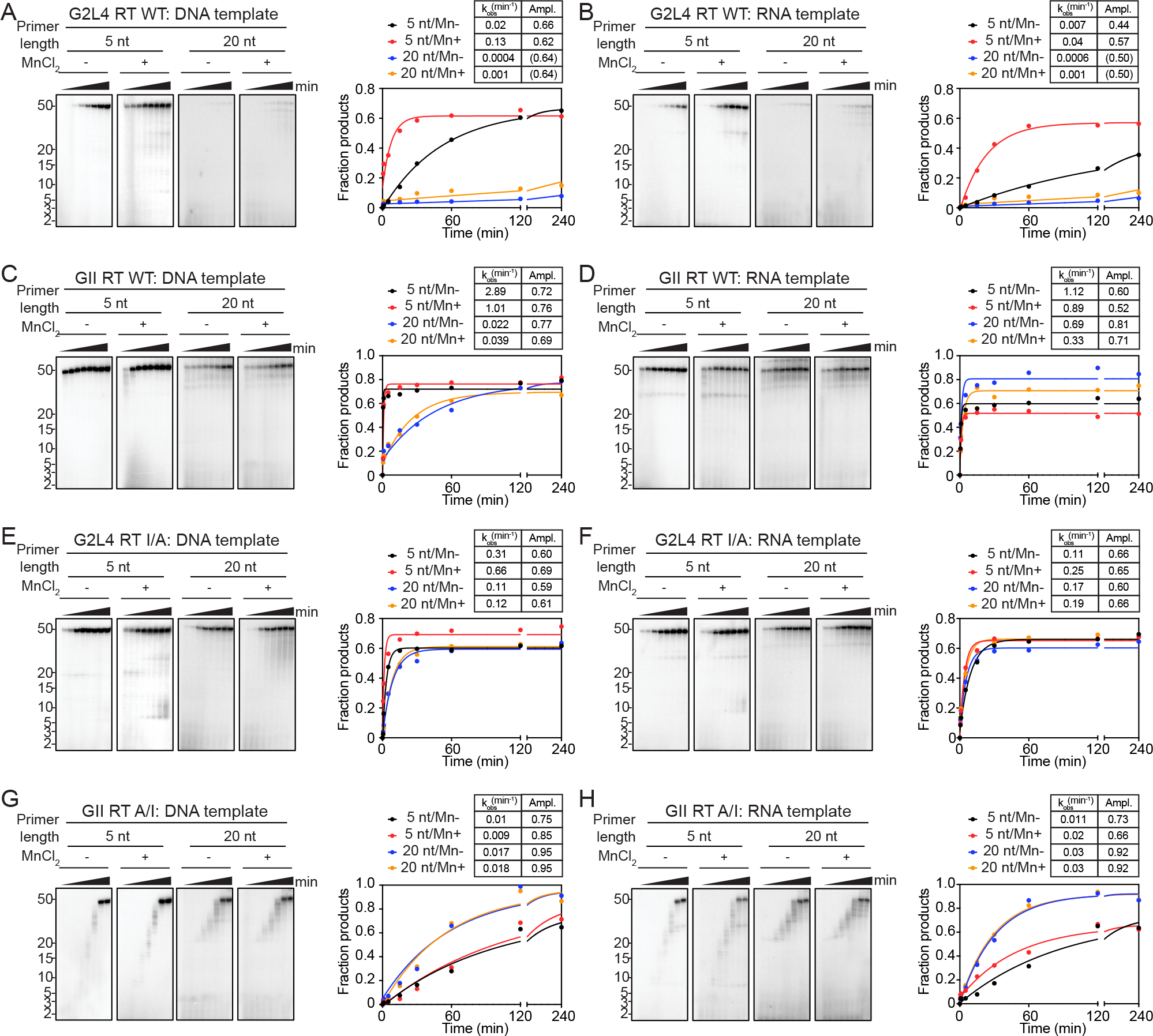
Effects of primer length and MnCl2 on the primer extension activity of WT and mutant G2L4 and GII RTs Primer extension reactions were done as described in Methods with 3’-blocked DNA or RNA tem- plates and 5- or 20-nt DNA primers in the presence or absence of 1 mM MnCl2. (A and B) Primer extension activity of WT G2L4 RT on DNA and RNA templates. (C and D) Primer extension activity of WT GII RT on DNA and RNA templates. (E and F) Primer extension activity of G2L4 I/A mutant RT on DNA and RNA templates. (G and H) Primer extension activity of GII RT A/I mutant RT on DNA and RNA templates. For all panels, the numbers to the left of the gel indicate the positions of size markers in a parallel lane. Tables above the plots show the rate constants (*k*obs) and amplitudes (Ampl.) for the production of the labeled 50-nt DNA product in panels A-F and for production of labeled products larger than 5 or 20 nt in panels G and H, respectively, obtained by fitting the data to a first-order rate equation. Values of Ampl. in parentheses indicate that the amplitude was fixed at the given value because the reaction did not reach an endpoint during the measurement time (see Methods).

**Figure S11.**
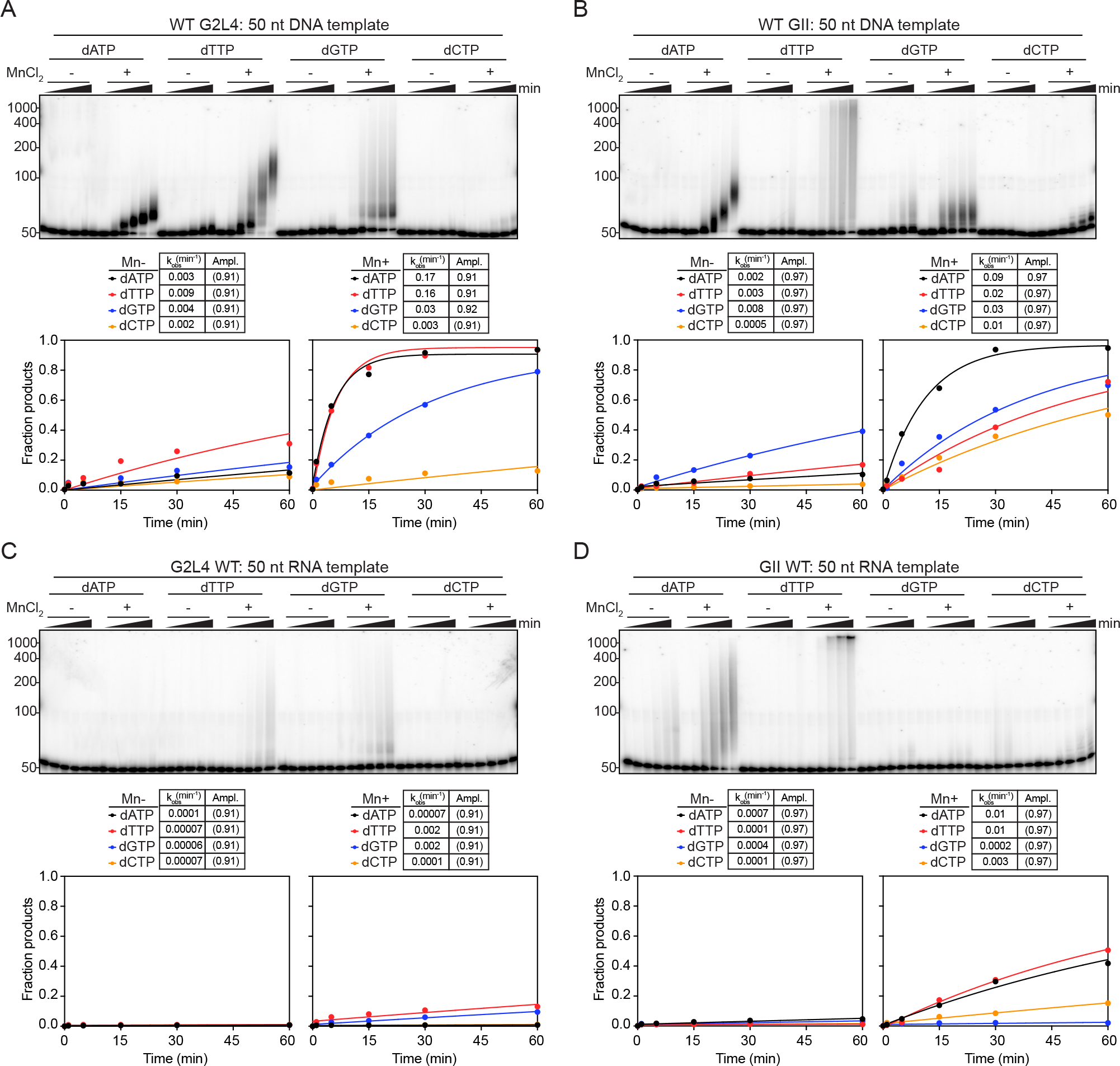
Terminal transferase time courses for WT G2L4 and GII RTs Terminal transferase time courses were done as described in Methods using 5’-labeled 50-nt DNA or RNA substrates in the presence or absence of 1 mM MnCl2. (A and B) Terminal transferase assays with WT G2L4 or GII RT using the DNA substrate. (C and D). Terminal transferase assays with WT G2L4 and GII RTs using the RNA substrate. For all panels, the numbers to the left of the gels indicate the positions of size markers in a parallel lane. Tables above the plots show the rate constants (*k*obs) and amplitudes (Ampl.) for production of all labeled products >50 nt obtained by fitting the data to a first-order rate equation. Values of Ampl. in parentheses indicate that the am- plitude was fixed at the given value because the reaction did not reach an endpoint during the measurement time (see Methods).

**Figure S12.**
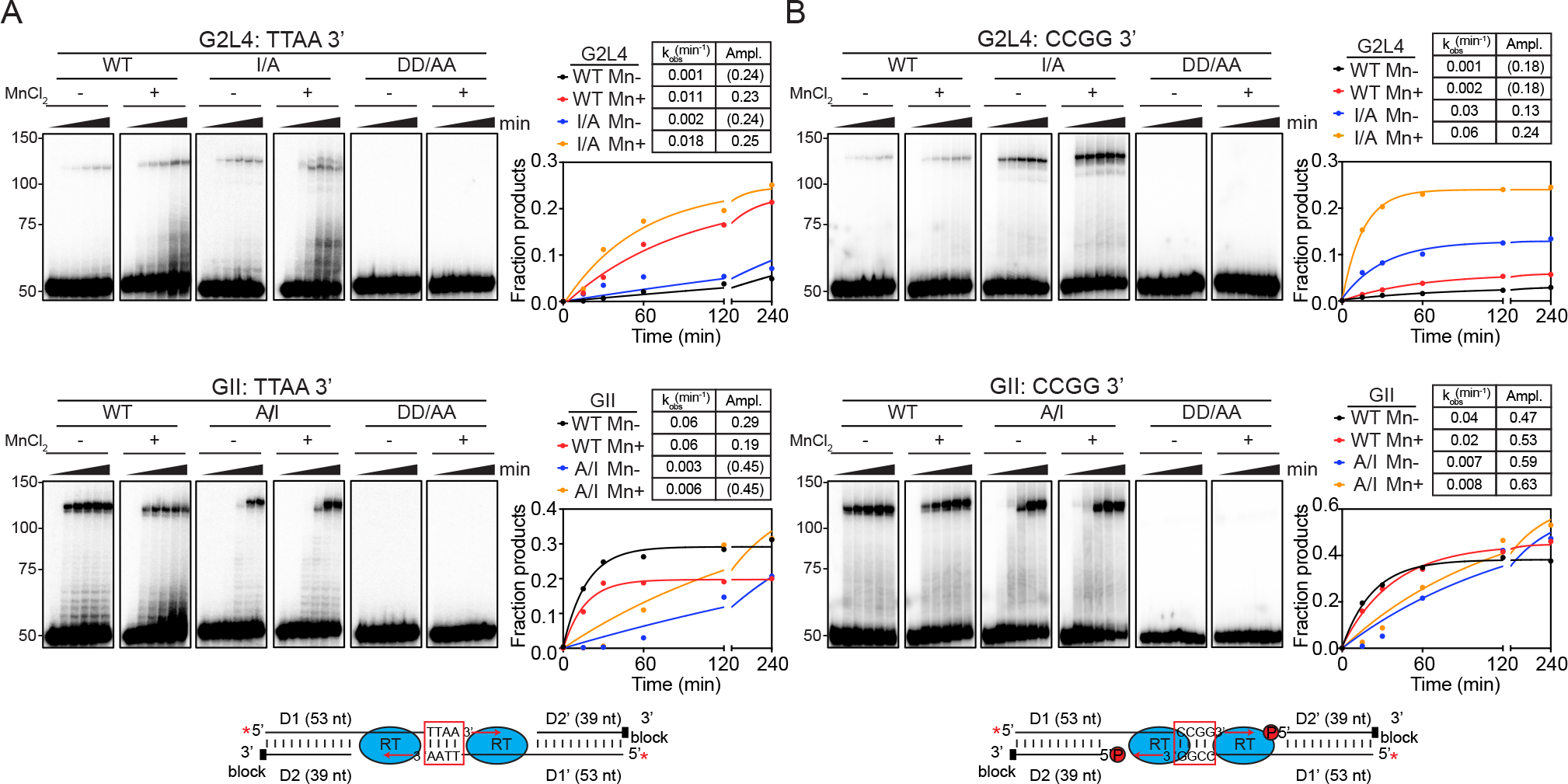
MMEJ time courses for WT G2L4 and GII RTs and effect of adding a 5’ phos- phate to the D2 oligonucleotide corresponding to the resected 5’ end at a double-strand break MMEJ reaction time courses were done as described in Methods using partially double-stranded 5’ labeled (red star) DNA substrates with 3’ overhangs having complementary TTAA or CCGG sequences at their 3’ ends in the presence or absence of 1 mM MnCl2. (A) MMEJ reactions using DNA substrates with 3’ overhangs having complementary 3’ TTAA sequences. (B) MMEJ reac- tions using DNA substrates with 3’ overhangs having complementary 3’ CCGG sequences as in Figure 5A but with a 5’ phosphate (red circled P in the schematic) at the 5’ end of the D2 oligonu- cleotide corresponding to resected 5’ end at a double-strand break. For all panels, the numbers to the left of the gels indicate the positions of size markers in a parallel lane. The plots show the fraction of substrate that is converted to products running between the 100- and 150-nt size mark- ers. Tables above the plots show rate constants (*k*obs) and amplitudes (Ampl.) obtained from fitting the data to a first-order rate equation. In parentheses indicate that the amplitude was fixed at the given value because the reaction did not reach an endpoint during the measurement time (see Methods).

**Figure S13.**
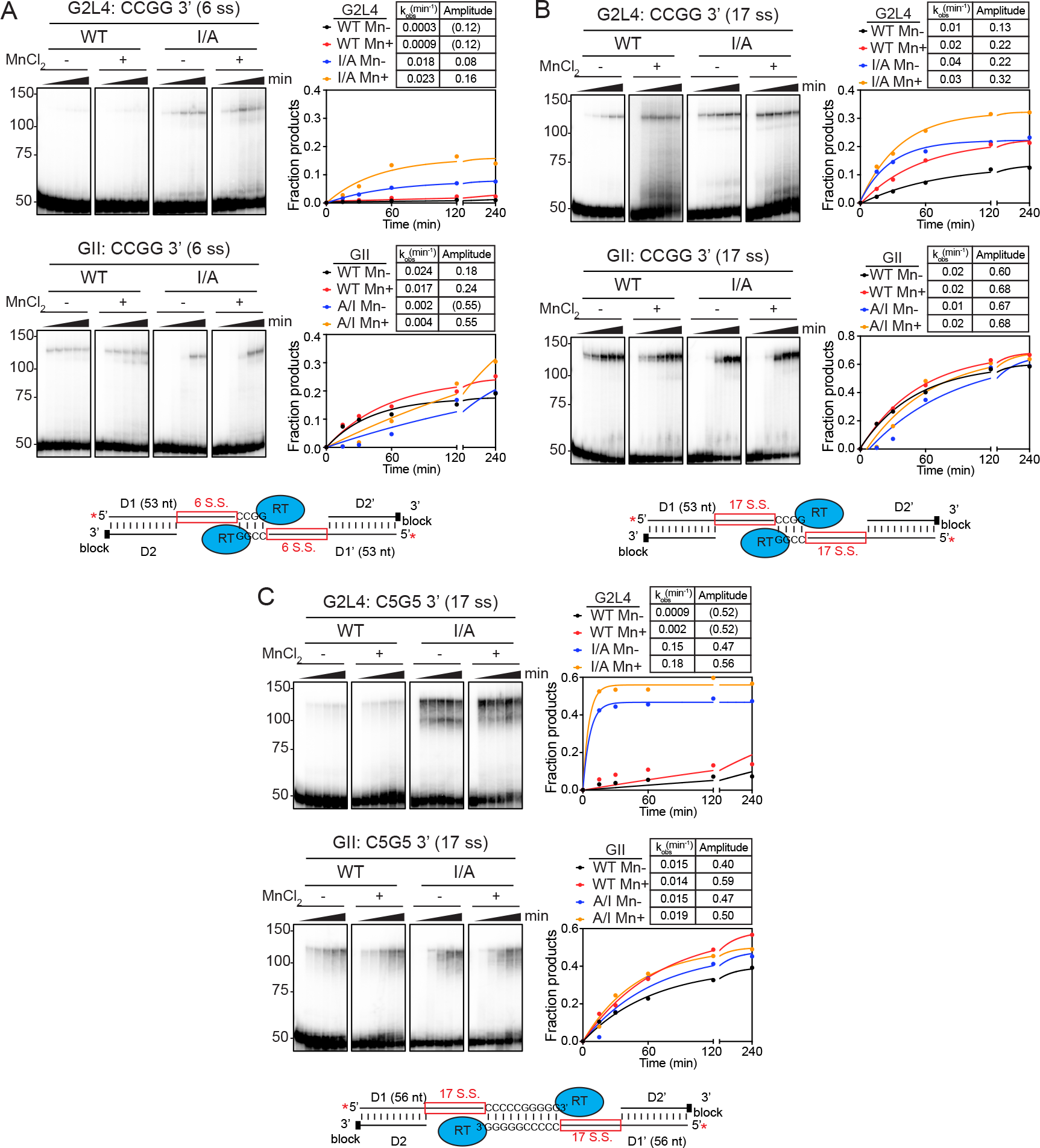
MMEJ time courses for WT and mutant G2L4 and GII RT with different length microhomologies and 3’ overhangs MMEJ reaction time courses were done as described in Figure S12 and Methods using partially double-stranded 5’-labeled (red star) DNA substrates having different length microhomologies and 3’ overhangs (see schematics at bottom of each panel). (A) MMEJ assay with CCGG (4 bp) mi- crohomology with 6-nt single-stranded gaps. (B) MMEJ assay with CCGG (4 bp) microhomologywith 17-nt single-stranded gaps. (C) MMEJ assay with CCCCCGGGGG (10 bp) microhomology with 17-nt single-strand gaps. For all panels, the numbers to the left of the gels indicate the posi- tions of size markers in a parallel lane. The plots show the fraction of substrate that is converted to products running between the 100- and 150-nt size markers. Tables above the plots show rate constants (*k*obs) and amplitudes (Ampl.) obtained from fitting the data to a first-order rate equation. Values of Ampl. in parentheses indicate that the amplitude was fixed at the given value because the reaction did not reach an endpoint during the measurement time (see Methods).

**Figure S14.**
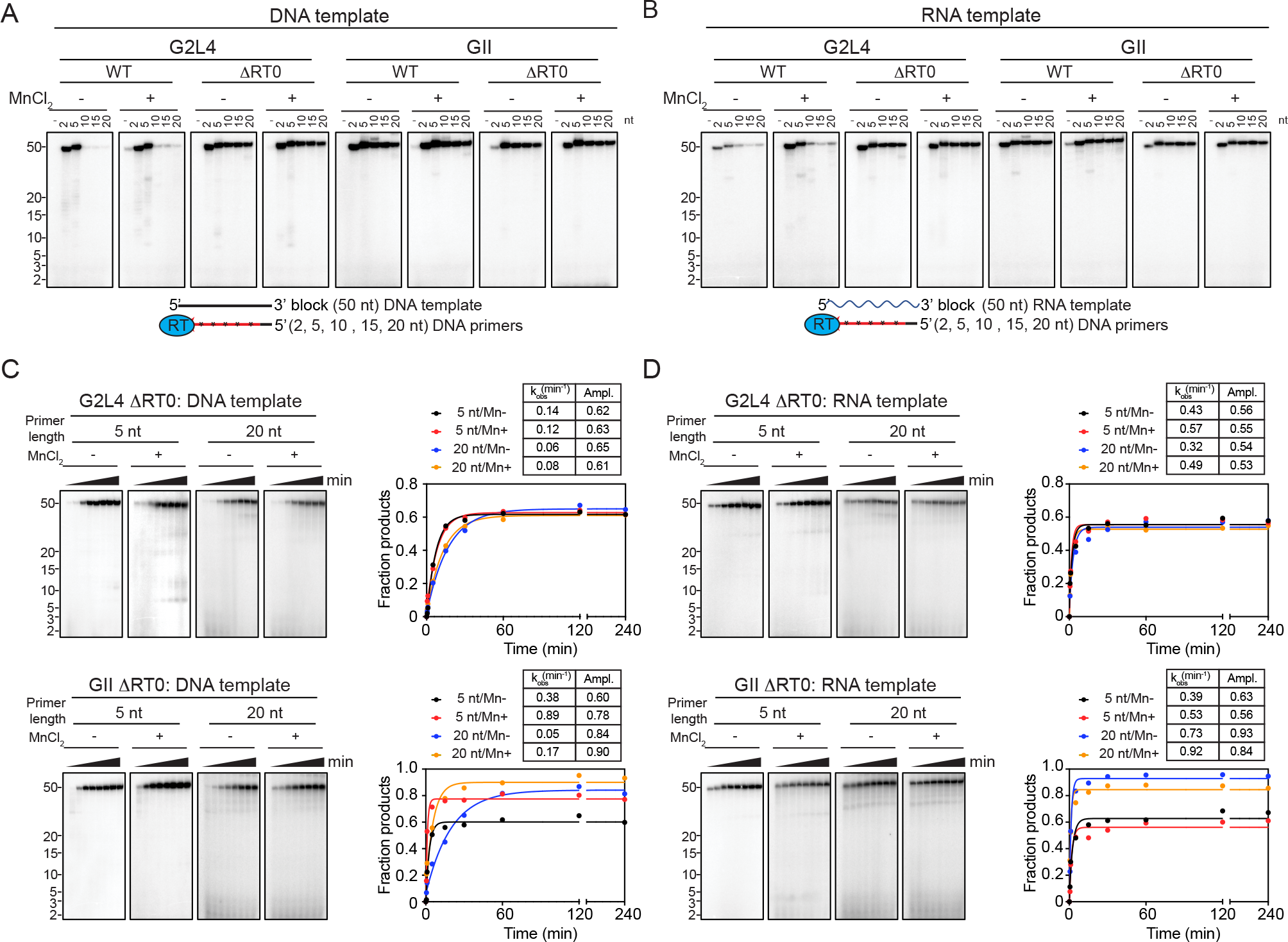
Effect of deleting the RT0 loop on the primer extension activity of G2L4 and GI RTs on DNA and RNA templates with different length DNA primers Primer extension assays with WT and *Δ*RT0 mutant G2L4 and GII RT were done as described in Figure S9 and Methods with 3’-blocked 50-nt DNA or RNA templates pre-annealed with different length DNA primers (100 µM 2 nt; 50 µM 5 nt; 500 nM 10, 15, 20 nt primers, respectively). (A and B) Primer extension assays for WT and *Δ*RT0 G2L4 and GII RTs with DNA and RNA tem- plates with no primer or annealed 2, 5, 10, 15, and 20 nt DNA primers (C and D) Primer extension time course experiments with G2L4 and GII ΔRT0 mutants with DNA or RNA templates with annealed 5-nt and 20-nt DNA primers with and without 1 mM MnCl2. For all panels, the numbers to the left of the gels indicate the positions of size markers in a parallel lane. Tables above the plots show the rate constants (*k*obs) and amplitudes (Ampl.) for production of labeled 50-nt DNA product obtained from fitting the data to a first-order rate equation.

**Figure S15.**
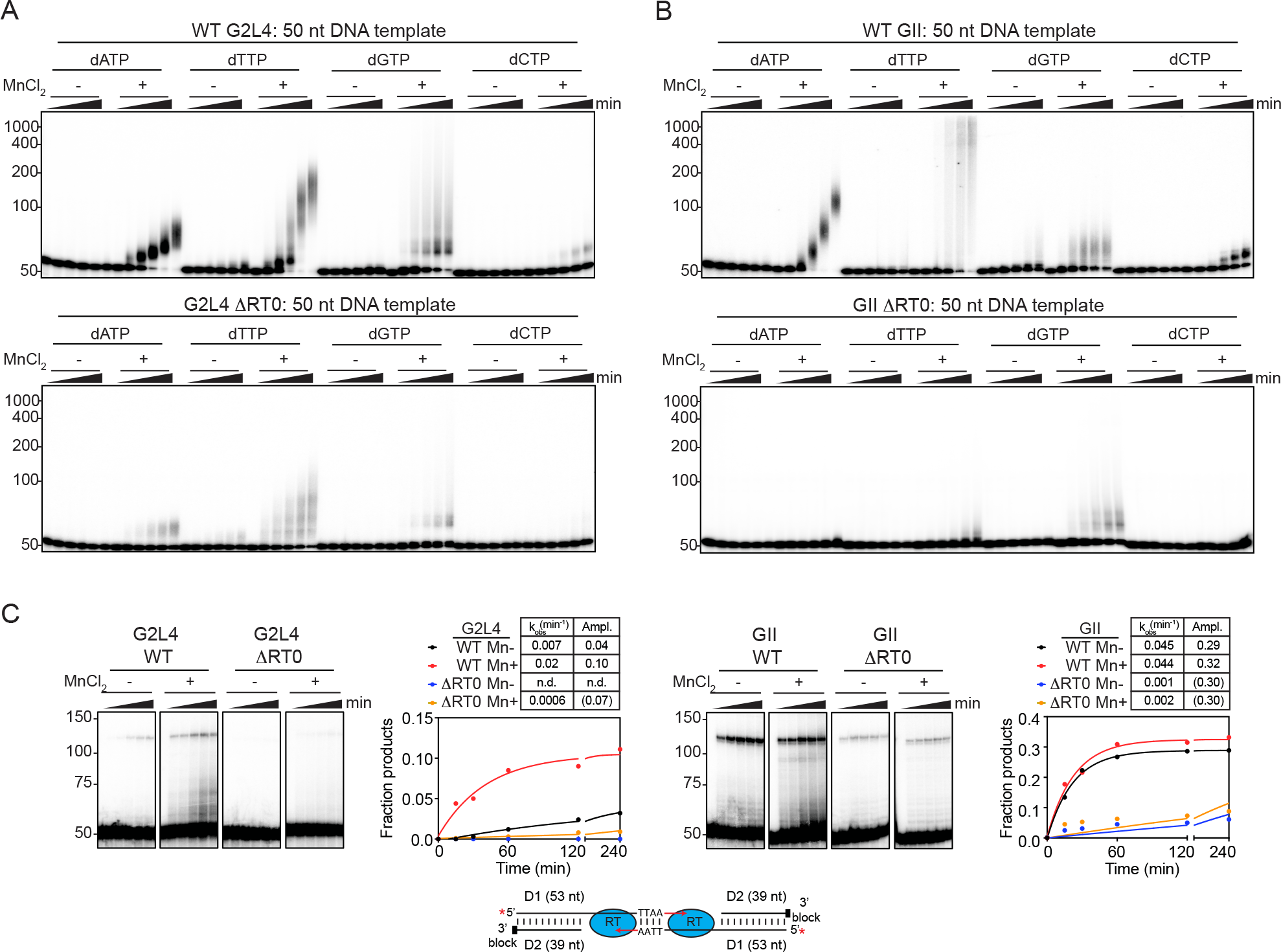
Effect of deleting the RT0 loop on terminal transferase and MMEJ activities of G2L4 and GII RTs Terminal transferase and MMEJ assays were done as described in Figure S11, Figure S13, and Methods. (A and B) Terminal transferase reactions time courses for WT (top) or *Δ*RT0 (bottom) G2L4 or GII RT using a 5-labeled 50-nt DNA substrate. Plots for the reaction time courses for production of labeled products >50 nt are shown in Figure 6C. (C) MMEJ assay by WT or *Δ*RT0 G2L4 and GII RTs using 5’-labeled DNA substrates having a 3’ overhang with 3’ TTAA microho- mology. The plots show the fraction of substrate that is converted to products running between the 100- and 150-nt size markers. For all panels, the numbers to the left of the gels indicate the posi- tions of size markers in a parallel lane. Tables above the plots show the rate constants (*k*obs) and amplitudes (Ampl.) obtained from fitting the data to a first-order rate equation. Values of Ampl. in parentheses indicate that the amplitude was fixed at the given value because the reaction did not reach an endpoint during the measurement time (see Methods).

**Figure S16.**
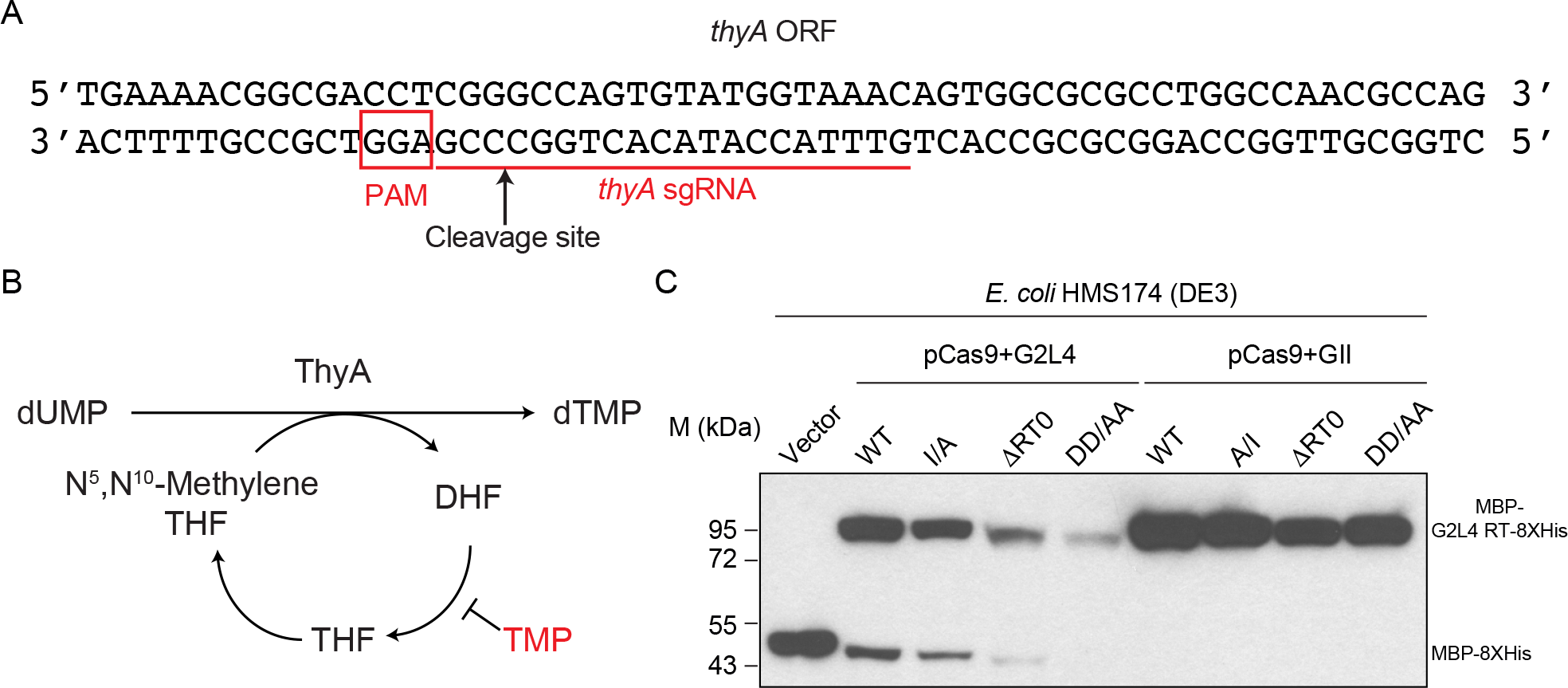
Cleavage of the *thyA* gene by the Cas9/*thyA* sgRNA complex (A) Sequence of a segment of the *E. coli* HMS174 (DE3) *thyA* gene showing the CRISPR/Cas9 cleavage site (arrow) and *thyA* guide RNA (underlined) and PAM (boxed) sequences. (B) *E. coli* thymidylate synthase pathway showing the basis for trimethoprim selection for *thyA* mutations. ThyA catalyzes the reductive methylation of 2’-deoxyuridine-5’-monophosphate (dUMP) to 2’- deoxythymidine-5’-monophosphate (dTMP) by using 5,10-methylenetetrahydrofolate (mTHF) as the methyl donor and reductant and yielding dihydrofolate (DHF) as a by-product. Trimethoprim (TMP) blocks the conversion of dihydrofolate to tetrahydrofolate (THF), which is needed for other cellular processes, resulting in cell growth arrest. (C) Immunoblot showing expression lev- els of WT and mutant G2L4 or GII RTs and MBP-8XHis from the vector control after induction of pCas9+G2L4 and GII RT plasmids in *E. coli* HMS174 (DE3). Immunoblots were done as in Figure S8.

**Table S1.**
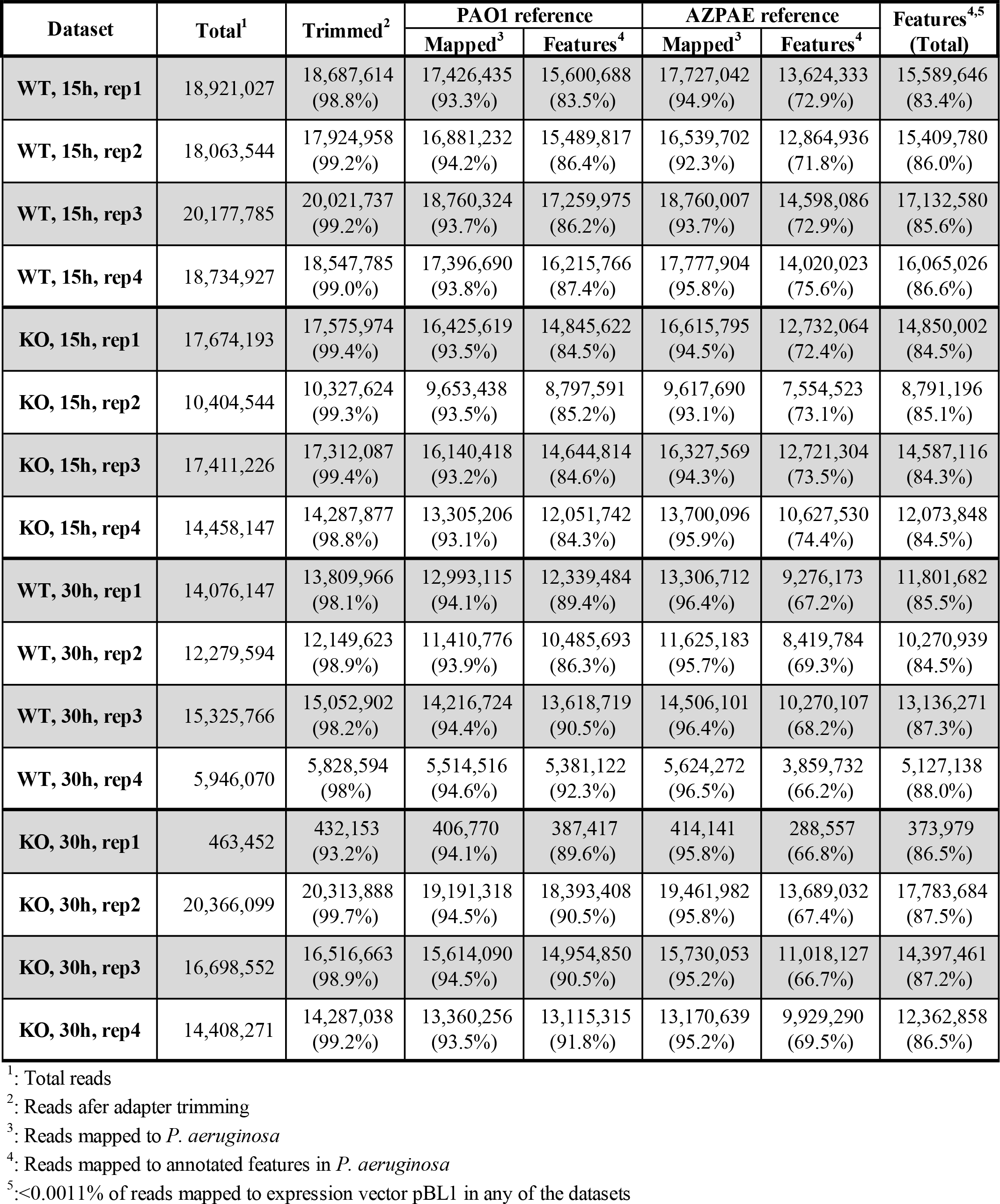
TGIRT-seq datasets and read mapping statistics

**Table S2.**
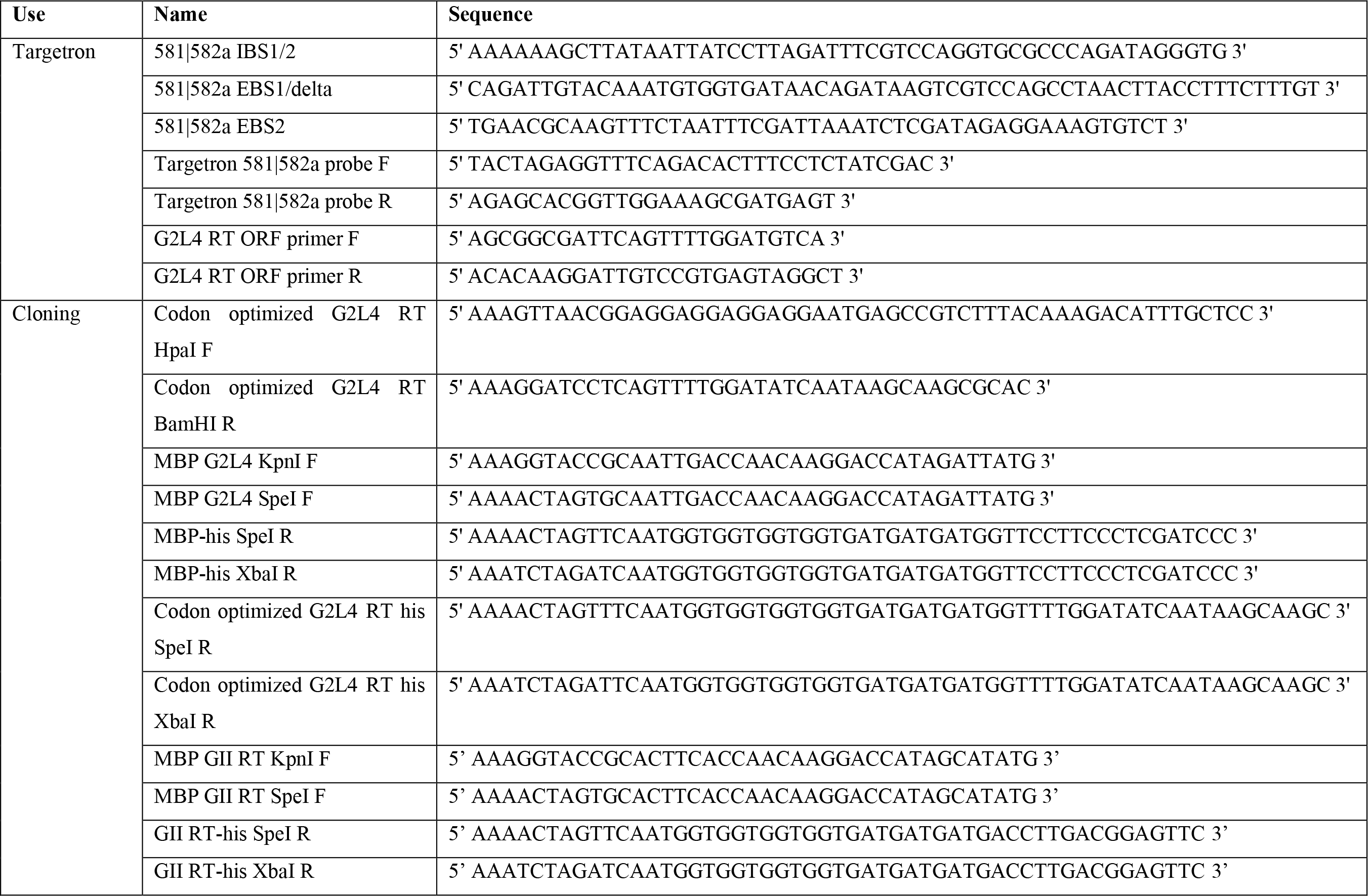
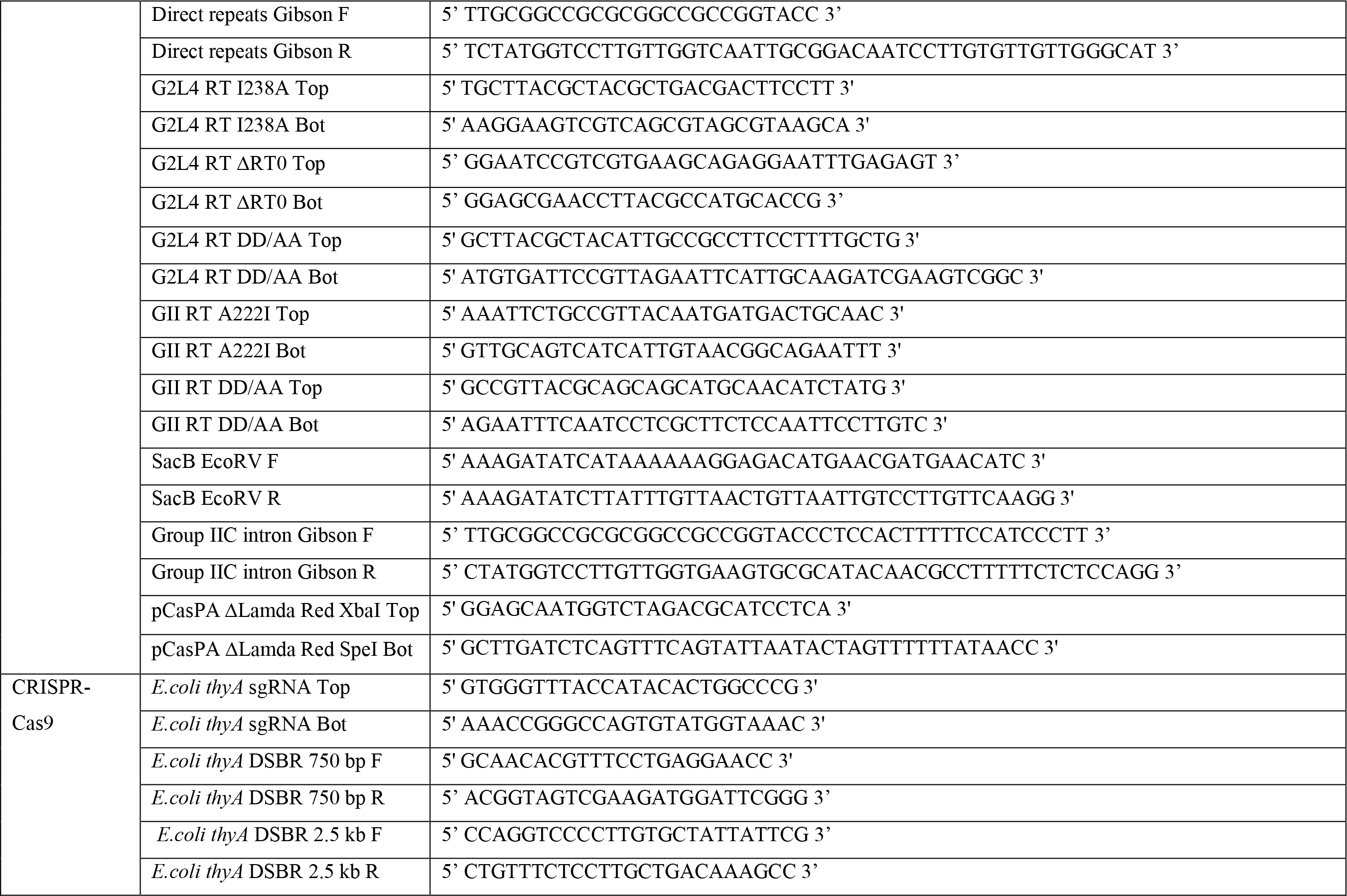
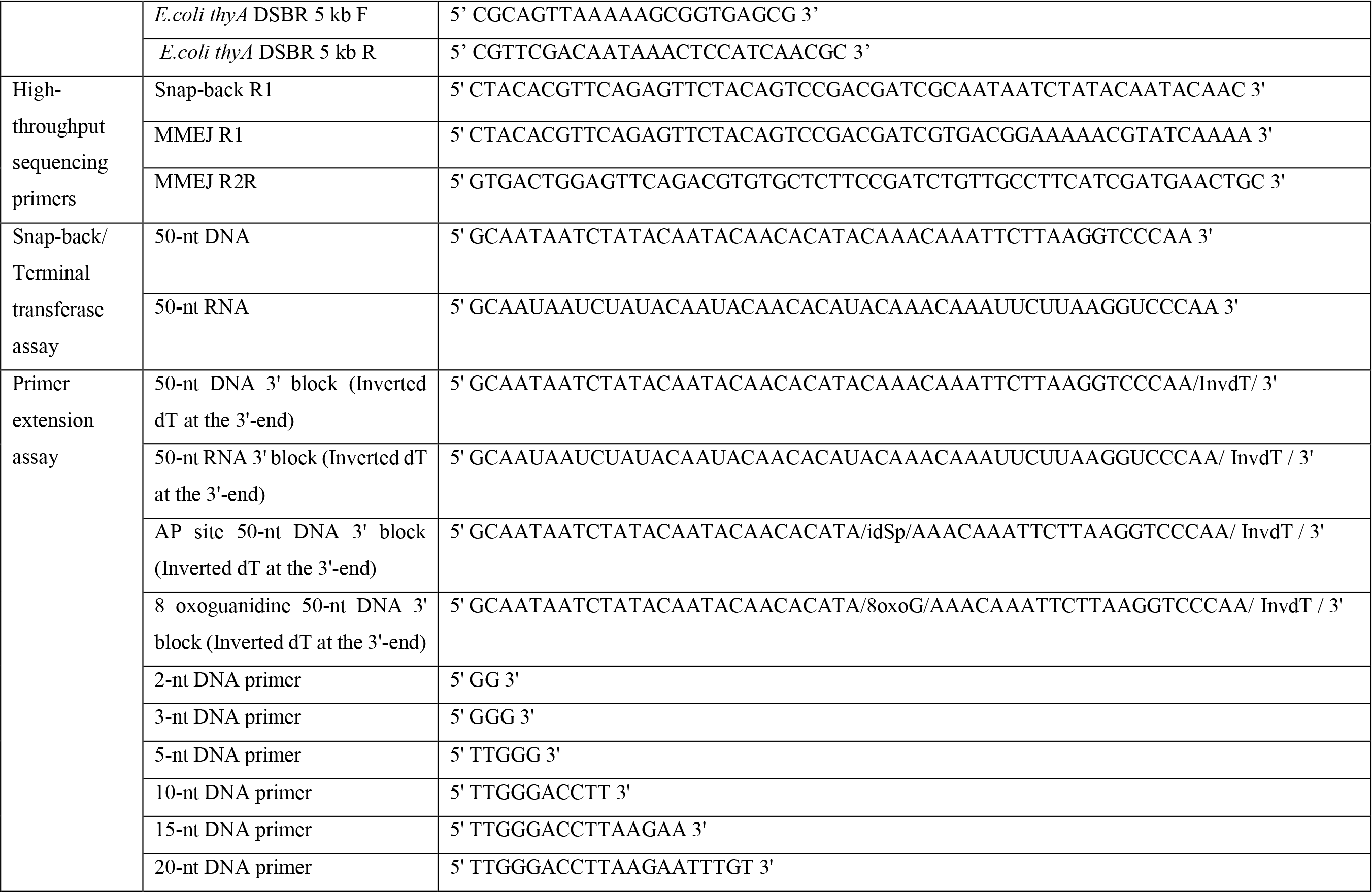
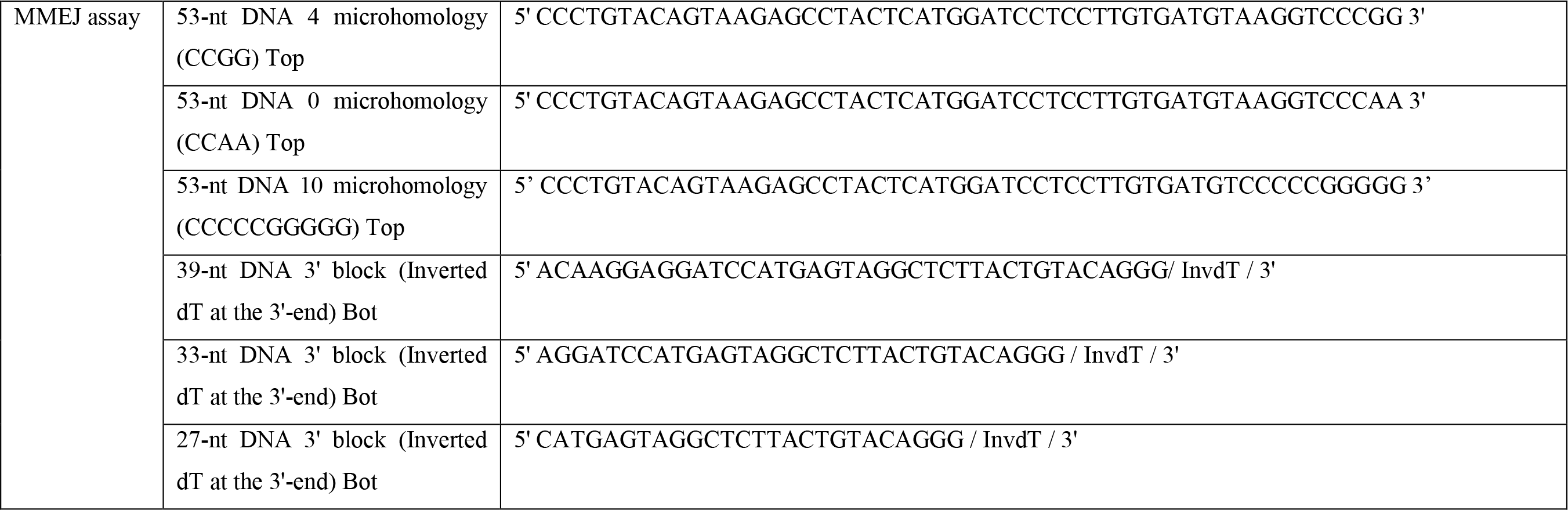
Oligonucleotides

**Table S3.** Recombinant plasmids

| Name | Source |
| --- | --- |
| pBL1 | Yao and Lambowitz, 2007 |
| pACD2-GsI-IIC-ΔORF+T7 | Mohr et al., 2018 |
| pBL1-G2L4 RT targetron | This study |
| pBL1-MCS | This study |
| pBL1-MBP-8xHis | This study |
| pBL1-MBP-G2L4 RT-8xHis | This study |
| pBL1-DR-MBP-G2L4 RT WT-8xHis | This study |
| pBL1-MBP-G2L4 RT DD/AA-8xHis | This study |
| pBL1-MBP-GII RT WT-8xHis | This study |
| pBL1-Intron-MBP-GII RT-8xHis | This study |
| pBL1-MBP-GII RT DD/AA-8xHis | This study |
| pMal-c5X | New England Biolabs (NEB #N8108) |
| pMal-G2L4 RT | This study |
| pMal-G2L4 RT I/A | This study |
| pMal-G2L4 RT DRT0 | This study |
| pMal-G2L4 RT DD/AA | This study |
| pMal-GII RT | Mohr et al., 2013 |
| pMal-GII RT A/I | This study |
| pMal-GII RT ΔRT0 | Stamos et al., 2017 |
| pMal-GII RT DD/AA | This study |
| pBluescriptII KS(+) | Agilent (212207) |
| pKS-SacB | This study |
| pACRISPR | Addgene (plasmid # 113348; Chen et al., 2018) |
| pACRISPR *thyA* sgRNA | This study |
| pCasPA | Addgene (plasmid # 113347; Chen et al., 2018) |
| pCas9SX | This study |
| pCas9+MBP-8x His | This study |
| pCas9+MBP-G2L4 RT-8xHis | This study |
| pCas9+MBP-G2L4 RT I/A-8xHis | This study |
| pCas9+MBP-G2L4 RT ΔRT0-8xHis | This study |
| pCas9+MBP-G2L4 RT DD/AA-8xHis | This study |
| pCas9+MBP-GII RT-8xHis | This study |
| pCas9+MBP-GII RT A/I-8xHis | This study |
| pCas9+MBP-GII RT ΔRT0-8xHis | This study |
| pCas9+MBP-GII RT DD/AA-8xHis | This study |

